# ECLARE: multi-teacher contrastive learning via ensemble distillation for diagonal integration of single-cell multi-omic data

**DOI:** 10.1101/2025.01.24.634799

**Authors:** Dylan Mann-Krzisnik, Anjali Chawla, Gustavo Turecki, Corina Nagy, Yue Li

## Abstract

Integrating multimodal single-cell data such as scRNA-seq and scATAC-seq is key for decoding gene regulatory networks. Still, integration remains challenging due to issues related to feature harmonization and limited quantity of paired data. To address these challenges, we introduce `ECLARE`, a novel framework combining multi-teacher ensemble knowledge distillation with contrastive learning for integrating unpaired single-cell multi-omic data. Briefly, `ECLARE` trains teacher models on paired datasets to guide a student model for aligning unpaired data, leveraging a refined contrastive objective and optimal-transport-based loss for precise cross-modality alignment. In computational benchmarking, experiments demonstrate `ECLARE`’s competitive performance in cell pairing accuracy, multimodal integration and biological structure preservation, indicating that multi-teacher knowledge distillation provides an effective means to improve a diagonal integration model beyond its zero-shot capabilities. In biological case studies, we demonstrate `ECLARE`’s applicability using unpaired snRNA-seq and snATAC-seq datasets in major depressive disorder (MDD). Firstly, our results revealed transcription factors and target gene combinations differentially regulated in depression with sex- and cell-type specificity. These findings further reveal gene regulatory interactions in excitatory neurons that are highly relevant to MDD neuropathology, such as those involving *EGR1, SOX2*, and *NR3C1*. Secondly, we show that `ECLARE` can learn continuous data manifolds useful for deciphering longitudinal biological processes in neurodevelopment and disease, revealing altered neurodevelopmental programs as potential regulators of depression in females, strongly associated with *EGR1* target genes. Altogether, we propose `ECLARE` as a robust solution for diagonal integration of unpaired multimodal single-cell data that enables the study of altered gene regulation in disease.

## Introduction

The development of multimodal single-cell technologies has enabled the acquisition of molecular data across multiple genomic modalities simultaneously ^1^. For example, SNARE-seq ^2^, SHARE-seq^3^ and the proprietary 10x Multiome simultaneously profile the transcriptome with single-cell RNA sequencing (scRNA-seq) and open chromatin regions or peaks with single-cell Assay for Transposase-Accessible Chromatin sequencing (scATAC-seq) within the same cell. This provides opportunities to investigate context-specific gene regulatory networks ^4^. However, paired multi-omic data are much less abundant than their unimodal counterparts and most combinations of single-cell modalities have not yet been successfully paired into multi-omic protocols. Hence, multi-omic protocols alone cannot currently reach the data volume currently achieved by single-cell atlases ^5;6^ nor can they fully account for the vast diversity of single-cell modalities currently available to study organ function and disease ^7^.

To increase data scale and modeling power to detect cis-regulatory programs between conditions (e.g., cases vs controls) or cell types, we may pool paired multi-omic data together. While gene features from scRNA-seq are universal across datasets, combining peaks from the scATAC-seq modalities across studies requires an arbitrary decision to either intersect or merge the peaks by their chromosomal coordinates. Neither is ideal: taking the intersection loses study-specific peaks and merging creates a large, redundant peak set. Another approach is to align peaks onto a set of *cis*-regulatory regions (cCREs) defined by a compendium study such as the ENCODE Project Consortium ^8^ or Human Enhancer Atlas ^6^. However, the ability to align peaks to cCREs is hampered by the variability in peak calling algorithms as well as the lack of universal mapping between peaks and cCREs ^9^, requiring further processing to promote alignability ^10^.

Meanwhile many methods have been developed to integrate *unpaired* single-cell multi-omic data ^11–16^. This task, known as *diagonal integration* ^1^, involves aligning cells from different modalities (e.g., scRNA-seq and scATAC-seq) that lack both shared cell identities and shared feature spaces. However, many limitations with previous methods remain unresolved. For instance, methodologies relying on gene activity scores such as Seurat ^17^ and ArchR ^18^ require transforming chromatin accessibility features to forcibly align with transcriptomic gene features. SCOT ^15^ uses the Gromov-Wasserstein Optimal Transport to align modalities but does not explicitly provide integrated representations that can be transferred to unseen data. Single-cell Contrastive Language–Image Pre-training (scCLIP) ^16^ performs CLIP-style contrastive learning (CL) using cell-type–based pseudo-pairs constructed from unpaired single-cell datasets without making use of the true paired multiome. scDART^13^ and GLUE^12^ incorporate prior knowledge about feature associations (e.g., between genes and peaks) to guide integration, which can be highly effective but may also constrain the discovery of novel or context-specific regulatory relationships if the prior knowledge is incomplete or biased. More recently, single-cell foundation models such as Geneformer^19^, UCE^20^, scGPT^21^, and scFoundation ^22^ have been tested for their diagonal integration capabilities, although falling behind more integration-focused methods^23^.

In this study, we propose **E**nsemble knowledge distillation for **C**ontrastive **L**earning of **A**TAC and **R**NA **E**mbeddings – `ECLARE` – a diagonal integration method for aligning unpaired RNA and ATAC cells using multimodal contrastive learning and multi-teacher *knowledge distillation* ^24–27^ (KD). While contrastive learning, knowledge distillation, and optimal transport are established paradigms, `ECLARE`’s conceptual novelty lies in its unique synthesis of these components to solve the diagonal integration problem: it leverages an ensemble of teachers trained on distinct paired datasets to guide a student model on unpaired data, further refined by a novel OT-based alignment loss (OT-CLIP) that enforces cross-modality matching during distillation without requiring global feature harmonization across all source datasets. We first demonstrate the benefit of KD for pairing RNA and ATAC, yielding superior pairing accuracy, integration and cell-type clustering metrics. Using `ECLARE` to pair RNA and ATAC separately generated from two recent studies ^28;29^, we aim to investigate differential gene regulatory networks (GRNs) in major depressive disorder (MDD). We also demonstrate that `ECLARE` can learn longitudinal patterns in neurodevelopmental data and show that key transcriptional regulators of inhibitory VIP neurons are expressed along developmental paths. These analyses on MDD and neurodevelopmental data lead the way to a third biological case study where `ECLARE` is used to co-embed MDD with neurodevelopmental data. Here, we identify neurodevelopment-related transcription modules and elucidate longitudinal processes associated with MDD. Our results on MDD GRN and co-embedding analyses reveal dysregulated regulatory interactions in MDD involving excitatory neurons and oligodendrocyte marker genes, and *EGR1* as a key TF of interest associated with later developmental signatures in MDD.

## Results

### ECLARE overview

`ECLARE` seeks to learn a shared embedding manifold that integrates unpaired scRNA-seq and scATAC-seq data via two stages of training (Fig. 1; Methods). First, we independently train a set of teacher models on paired RNA & ATAC multi-omics *source* data (Fig. 1a). Using the InfoNCE loss ^30^, a given teacher model is trained to force alignment between matching RNA and ATAC cells leading to an integrated latent space, where pairs of matching cells share highly similar coordinates. Second, we distill the knowledge from this ensemble of trained teacher models into a student model using the *target* dataset for which in silico integration is desired (Fig. 1b). Importantly, this target dataset is unpaired. Therefore, `ECLARE` relies on the ability of the teacher models to perform multimodal alignment without tuning on target data, i.e. zero-shot transfer. This multi-teacher distillation process relies on a KD loss (Fig. 1c) and a novel alignment loss based on Optimal Transport (OT) that we term *OT* -CLIP (Fig. 1d). Both the teacher-specific KD and alignment losses are combined across teachers to create ensemble multi-teacher KD and alignment losses via weighted averaging (Fig. 1b).

**Figure 1:**
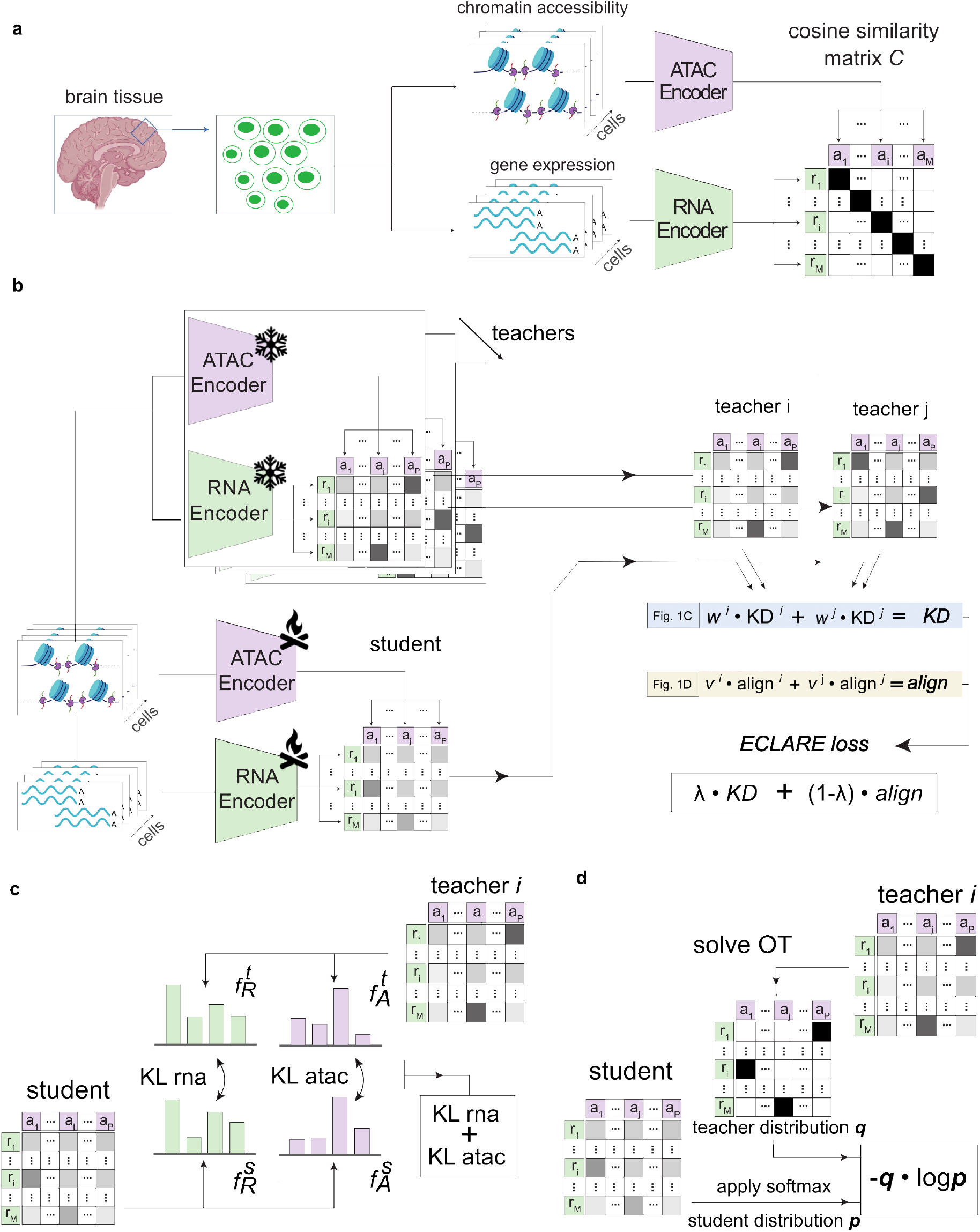
Overview of `ECLARE`. **(a)** Contrastive learning on paired data. Teacher neural network encoders are pretrained to align matched nuclei in a shared latent space on paired multiome data. **(b)** Multi-teacher knowledge distillation. A student model processes a target dataset while frozen teachers provide ATAC–RNA similarity matrices used in the KD and alignment losses. **(c)** KD loss. Row-/column-wise softmax distributions from student and teacher similarity matrices are compared using KL divergence. **(d)** Alignment loss (OT-CLIP). Optimal Transport applied to the teacher similarity matrix produces one-to-one RNA–ATAC matchings; cross-entropy between these matchings and the student similarity matrix defines the alignment loss. Teacher and student matrices are square, i.e. *M* = *P*.

### Contrastive learning provides a strong basis for multi-omic alignments

We compared CLIP with the existing state-of-the-art methods including multiVI ^11^, GLUE^12^, scDART^13^ and scJoint^14^ on evaluation metrics quantifying nucleus pairing (1-FOSCTTM), integration (iLISI) and clustering (ARI, NMI & ASW) using test paired data, which were not seen by any of the methods (Methods). We also adapted the vertical integration method MOJITOO ^31^ for diagonal integration, achieving high-quality integration of paired data and, given MOJITOO’s top performance in vertical RNA+ATAC integration reported by^32^, high potential for transferability to diagonal settings. For these benchmarking experiments, models were trained on a paired source dataset (e.g., **DLPFC_Anderson**) and evaluated on a separate paired target dataset (e.g., **Midbrain_Adams**) to evaluate zero-shot transfer. Given the paired nature of the datasets for these experiments, we considered 1-FOSCTTM as the key ground truth metric, which quantifies the quality of nuclei pairing across ATAC and RNA modalities with respect to ground truth pairings. CLIP provides high-quality pairings based on a median value of 0.88, outperforming the second-best model by a large margin (multiVI with 0.62; Fig. 2a.i). We also assessed the quality of the broader integration and alignment of nuclei by iLISI (Fig. 2a.ii). Most methods do not inherently force alignment between modalities, which resulted in poor iLISI scores. Only scDART has the median iLISI value exceeding that of CLIP. In terms of clustering by known cell types, CLIP is also superior based on ARI and NMI (Fig. 2a.iii-iv) and conferred slightly lower ASW compared to GLUE, scDART and scJoint (Fig. 2a.v). Together, these results favor the contrastive learning approach, making it a suitable basis for `ECLARE`.

**Figure 2:**
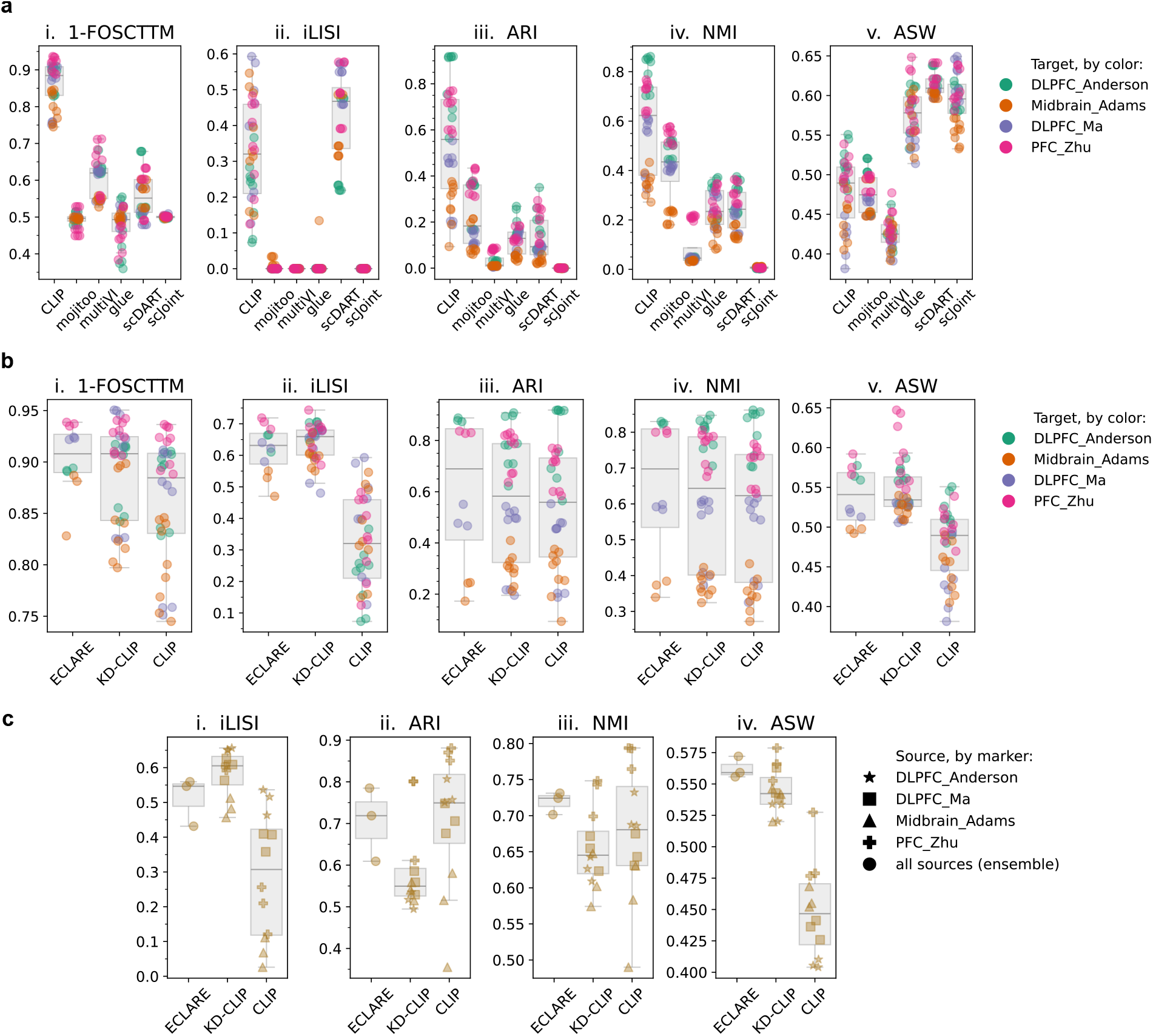
Integration and clustering metrics on benchmarking and knowledge distillation experiments. Subplots for each metric are further labeled with roman numerals i-v. **(a)** Benchmarking results on zero-shot alignment task with paired data to compare different baseline methods. Data points are color-coded by the target dataset evaluated for each experiment. **(b)** Comparison of `ECLARE`, KD-CLIP & CLIP methods on paired data, with data points color-coded by the target dataset evaluated for each experiment. **(c)** Comparison of `ECLARE`, KD-CLIP & CLIP methods on unpaired MDD data, with data points shape-coded by the source dataset(s) used to train individual models. For `ECLARE`, we refer to the source dataset as “all sources (ensemble)” since all source datasets are used.

### Knowledge distillation improves multi-omic alignment and cell-type clustering

We investigated on paired datasets whether incorporating single-teacher (KD-CLIP) and multi-teacher (`ECLARE`) KD improves the performance of the base CLIP model (Fig. 2b). The baseline CLIP achieves a median 1-FOSCTTM score of 0.88 on target data that is further increased by single-teacher and multi-teacher KD (Fig. 2b.i). Both of the latter achieve median scores around 0.91, although `ECLARE` demonstrates lower variability than KD-CLIP. In terms of multimodal integration, the median iLISI score of 0.31 for CLIP shows much room for improvement. Indeed, KD-CLIP and `ECLARE` show a marked increase in iLISI, respectively attaining values of 0.66 and 0.64 (Fig. 2b.ii). In terms of ARI and NMI clustering metrics, there is a cumulative increase when incorporating single-teacher (KD-CLIP) and then multi-teacher (`ECLARE`) KD from median values of 0.55 to 0.58 and 0.69 for ARI and from 0.63 to 0.65 and 0.70 for NMI (Figs. 2b.iii-iv). Lastly, we also observed improvements in ASW for both KD-CLIP and `ECLARE` compared to CLIP from 0.49 to 0.53 and 0.54, respectively (Fig. 2b.v), which are only slightly lower compared to GLUE, scDART and scJoint reported previously in Fig. 2a.v. These results show that the single-teacher KD strategy confers accurate multi-omic pairing and cell embeddings that preserve known cell-type structure. Moreover, the multi-teacher `ECLARE` strategy provides the best balance between modality-correction and biological preservation.

### ECLARE integrates unpaired nuclei from multi-omic MDD data

The majority of genetic variants associated with MDD are located in non-coding regions of the genome ^33^. This suggests that much is to be gained in combining both modalities to decode the *cis*-regulatory abnormalities in MDD ^34^. Earlier studies have separately generated snRNA-seq ^28;35^ and snATAC-seq ^29^ to study this mood disorder. As a case study, we sought to align these unpaired multi-omic nuclei to elucidate differential gene regulatory networks in MDD, with integration and clustering metrics shown in Fig. 2c. Due to the lack of ground-truth knowledge of nuclei pairing, we could not use 1-FOSCTTM to assess pairing accuracy and instead considered iLISI as the primary indicator of the quality of integration. Although KD-CLIP shows the greatest improvement on iLISI from 0.31 for CLIP to 0.61 (Fig. 2c.i), `ECLARE` outperforms KD-CLIP in terms of ARI, NMI and ASW (Figs. 2c.ii-iv). Therefore, `ECLARE` provides better integration than CLIP while also maintaining compact clusters concordant with cell-type annotations.

Lastly, we also assessed batch correction performance using k-nearest neighbor batch effect test (kBET - Fig. S1). On paired data, `ECLARE` and KD-CLIP achieve comparable median kBET acceptance rates (0.45), both above CLIP (0.42), while `ECLARE` exhibits a tighter spread across target datasets (Fig. S1a). On MDD data, `ECLARE` and KD-CLIP attain a median acceptance rate of 0.72 and 0.75, respectively, with a notably narrower distribution than CLIP, which reaches a lower value at 0.57 (Fig. S1b). Together, these results suggest that `ECLARE` yields embeddings whose batch mixing is more consistent across source–target configurations, rather than uniformly higher, consistent with its role as an ensemble distillation of single-teacher models.

### Cell-type–specific enrichment of MDD-associated gene programs in excitatory neurons and oligodendrocytes

To identify dysregulated genes in MDD, we employed the trained MDD student dual encoder, which conferred good integration and cell-type clustering (Fig. 2c; Fig. 3a), to pair the nuclei between modalities from the unpaired MDD multiomic datasets^28;29^. Nuclei were sampled by sex and cell type, projected into the `ECLARE` latent space, and paired one-to-one across RNA and ATAC modalities using a two-step optimal transport procedure to emulate paired data. Genes were then filtered using a statistical framework derived from sc-compReg, a computational method for comparing gene regulatory networks between conditions using single-cell data^37^. This sc-compReg-derived framework integrates gene expression, regulatory element accessibility, and transcription factor activity, which we followed with biological pathway enrichment analysis performed separately for each sex & cell-type combination. Refer to the Methods section for more details.

**Figure 3:**
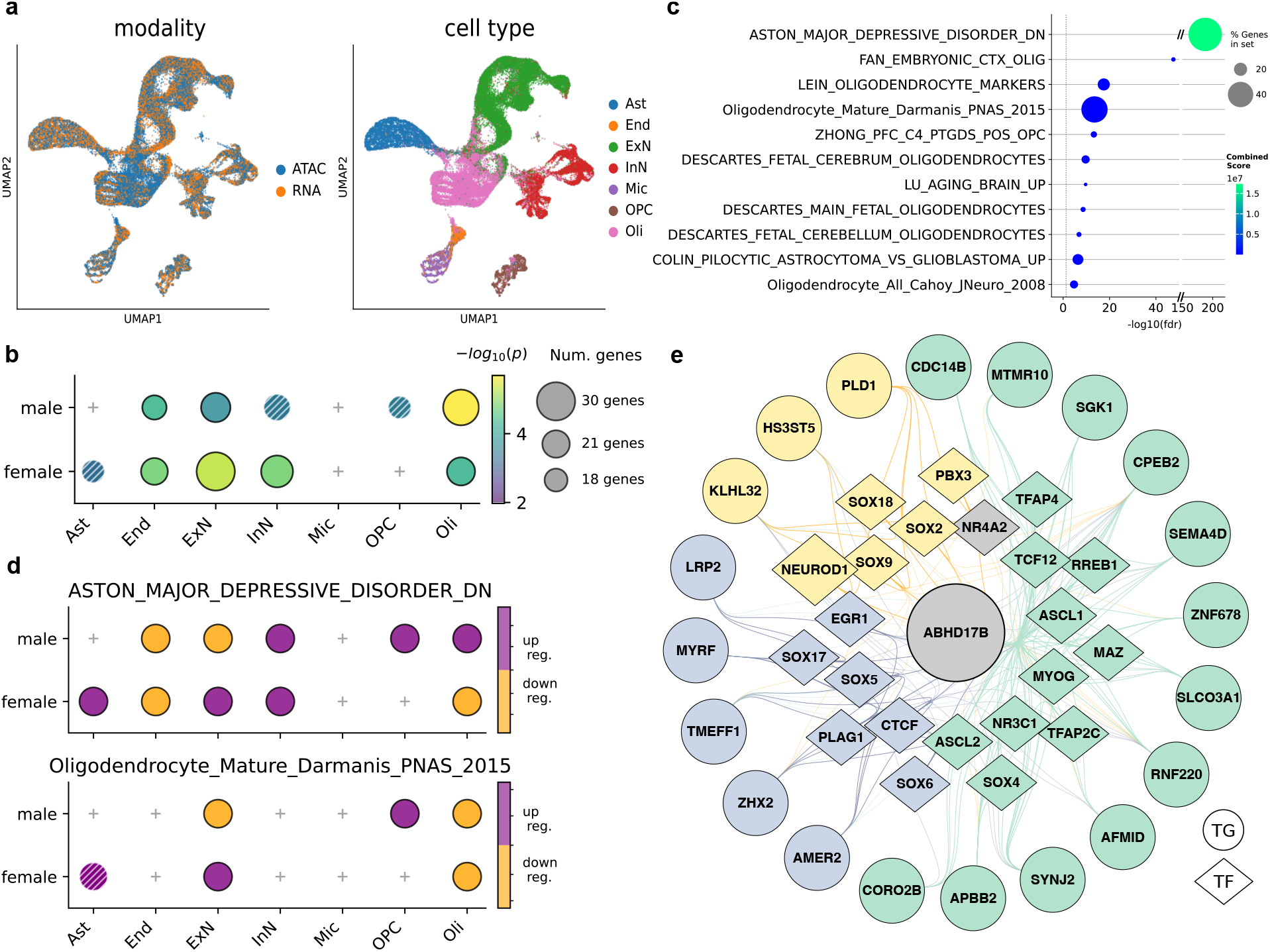
Downstream biological analyses following pairing of nuclei in `ECLARE` latent space. **(a)** UMAP visualizations of held-out MDD nuclei embedded using `ECLARE`, colored by modality and cell type. **(b)** Dotplot of aston major depressive disorder dn gene set enrichment, where dots are colour-coded by corrected p-value and dot size corresponds to the number of overlapping genes. Only dots associated with statistically-significant enrichment are shown (-log_10_(*p*) *>* 1.3, pre-Bonferroni). Dots for results not satisfying an additional Bonferroni correction for repeated comparisons (n=14) are hashed in white. **(c)** Dotplot arising from a two-step MDD enrichment analysis, with a vertical reference line added to locate the FDR-corrected p-value of 0.05. The horizontal axis is broken into two segments to accommodate the extreme significance of the aston major depressive disorder dn gene set without compressing the remaining pathways. **(d)** Weighted t-test between cases and controls based on module scores for pathways associated with aston major depressive disorder dn (top) or oligodendrocyte markers (bottom). **(e)** A differential GRN subnetwork, centered around the role of *ABHD17B* in female excitatory neurons. Circles represent target genes (TGs) and diamonds represent transcription factors (TFs). Nodes colored by cluster (gray nodes excluded from clustering analysis) and edges colored by the cluster of the TF involved in the edge’s TF-TG association. ExN: excitatory neuron, InN: inhibitory neuron, Oli: oligodendrocyte, OPC: oligodendrocyte precursor cell, End: endothelial, Ast: astrocyte, Mic: microglia.

We noticed widespread enrichment across sex & cell-type combinations for a gene set titled aston major depressive disorder dn, using the *Brain*.*GMT* resource^38^. TF target genes from excitatory neurons (ExN), endothelial cells (End) and oligodendrocytes (Oli) showed significant Bonferroni-corrected enrichment for this MDD-related gene set in both sexes (Fig. 3b, Bonferroni-corrected *p <* 0.05). Other gene sets related to neuropsychiatric disorders were identified, such as those associated with schizophrenia, bipolar disorder, and Alzheimer’s disease (Fig. S2). We validated the enrichment for aston major depressive disorder dn using GREAT^39^ and H-MAGMA^40^, and found this enrichment to be statistically significant and among the strongest associations across both sexes for excitatory neurons and oligodendrocytes, second only to oligodendrocyte precursor cells (OPC) from male donors (Fig. S3). Moreover, H-MAGMA based on MDD GWAS and HiC datasets revealed the strongest signals in oligodendrocytes (across both sexes) and OPC (male) (Fig. S4). Therefore, our results show oligodendrocytes and excitatory neurons to be strong candidates for depression. Next we investigated relationships between the MDD gene set and other *Brain*.*GMT* biological pathways by pooling filtered genes overlapping with the MDD gene set using a leading-edge-style, two-step enrichment analysis ^41;42^. Specifically, significant genes from sex & cell-type combinations overlapping genes from the MDD gene set were aggregated into a single gene set and provided as input to EnrichR. Once again, these results highlighted oligodendrocyte and OPC marker gene sets as being highly associated with MDD, accompanied by other pathways of interest relating to brain aging (lu aging brain up) and glioblastoma (Fig. 3c).

To validate our results, we performed differential gene expression analysis, which not only confirmed the enrichment results but identified the target cell types (Fig. 3d, top). We observed differential expression for oligodendrocyte-related genes present in both sexes for excitatory neurons and oligodendrocytes (Fig. 3d, top), suggesting a potentially shared role of excitatory neurons and oligodendrocytes in MDD, such as those related to myelination ^28;43^. blalock_alzheimers_disease_up and Gandal_2018_bipolardisorder_upregulated_cortex_also exhibit differential expression for excitatory neurons and oligodendrocytes for both sexes (Fig. S5).

Previous studies suggest converging evidence that myelin- and oligodendrocyte-lineage genes are dysregulated in cortex of depressed individuals^43^. Likewise, differential expression using snRNA-seq data in male MDD cases found transcriptional changes are concentrated in deep-layer excitatory neurons and immature OPCs, implicating potential communication between these cell types that involves FGF signaling, steroid hormone receptor chaperone cycling, immune functions and cytoskeletal regulation^28^. In line with this, enrichment for oligodendrocyte/OPC marker sets from our model further supports MDD-associated gene programs spanning both excitatory neurons and oligodendrocyte lineage cells, potentially involved in myelination and neuron–glia communication. A more recent study mapping the genetic landscape across 14 psychiatric disorders detected the strongest genetic signal in oligodendrocytes for the “Internalizing factor” class of psychiatric disorders, of which major depression is a member ^44^. The Internalizing factor also showed enrichment in excitatory neurons, further supporting our finding that both oligodendrocytes and excitatory neurons are key cell types involved in the neuropathology of MDD.

In sum, multiple enrichment analyses converge on excitatory neurons and oligodendrocyte lineage cells as key contributors to MDD across both sexes. Several oligodendrocyte and OPC marker gene sets show significant enrichment and differential regulation, suggesting involvement of oligodendrocyte-related gene programs in MDD.

### The gene regulatory effects of MDD-associated TFs are altered in excitatory neurons from female MDD donors

We performed a GRN pruning process that isolated differentially regulated GRN edges between TFs and their target genes based on sc-compReg, this time targeting individual TF-TG edges rather than aggregating at the TG level (see Methods). Because *NR4A2* serves as a cell-defining transcription factor for the specific deep-layer excitatory neurons most vulnerable to stress-induced molecular alterations in ^29^, we specifically examined its activity-dependent regulatory influence in these cells by looking for pruned TF-TG edges where *NR4A2* was a significant TF in excitatory neurons. We found that the interaction between *NR4A2* and *ABHD17B* is disrupted in excitatory neurons from female donors. We then searched for other TFs whose interactions with *ABHD17B* are disrupted in female excitatory neurons and further added their disrupted target genes as well. This process of first anchoring the search via *NR4A2* in excitatory neurons, then identifying *ABHD17B* as a key target and expanding back out from *ABHD17B* led to a GRN network that places *ABHD17B* as the central hub TG (Fig. 3e). We performed spectral co-clustering (*n_clusters*=3) on the set of TF-TG edges that constitute this GRN to identify broad biological categories of regulatory activity: the green cluster includes genes related to stress response signaling and synaptic homeostasis (TFs: *NR3C1, ASCL1* ; TGs: *SGK1, SEMA4D*) ^45–48^, the purple cluster includes synaptic plasticity and activity-dependent myelination (TFs: *EGR1, CTCF* ; TGs: *MYRF, LRP2*) ^49–52^, and the yellow cluster includes neurogenesis and neural progenitor maintenance (TFs: *SOX2, NEUROD1* ; TGs: *PLD1, HS3ST5*) ^53–56^. As a computationally inferred central hub, *ABHD17B* is hypothesized to integrate these regulatory axes by acting as a depalmitoylase that modulates the membrane localization of signaling proteins essential for synaptic remodeling, stress resilience, and neural maturation within MDD-vulnerable excitatory neurons ^57^. According to GTEx^58^ and psychSCREEN^33^, *ABHD17B* is highly expressed in brain & spinal cord (Fig. S6a), showing an increasing gradient of expression from superficial cortical layers to deep layers and white matter (Fig. S6b) that is more elevated in neurons and oligodendrocytes than in other major cell types (Fig. S6c). While this expression pattern aligns with the shared MDD-associated signal observed earlier in excitatory neurons and oligodendrocytes, these *in silico* findings remain hypothesis-generating and require future experimental validation. Moreover, according to the TF-enhancer-TG triplets associated with the TF-TG edges shown in Fig. 3e (GRN enhancers not shown), *NR4A2, EGR1, NR3C1* and *PLAG1* share a common enhancer region located at chr9:71768678–71769178 involved in the differential co-regulation of *ABHD17B*.

Consistent with this observation, Chawla et al. ^29^ also reported evidence for interactions between *NR4A2* and *EGR1* based on StringDB. *NR4A2* is an immediate-early gene known to be dysregulated by stress ^59^. Moreover, *NR4A2* + deep-layer excitatory neurons showed significant enrichment for MDD-associated genetic variants and chromatin remodeling ^29^. Also, *EGR1* was identified as a TF with altered chromatin footprinting in excitatory neurons ^29^ with known involvement in multiple neuropsychiatric conditions ^60^. On the other hand, *SOX2* is well established as a regulator of oligodendrocyte development and CNS remyelination ^61^. In the adult murine cortex, *SOX2* is predominantly expressed in astrocytes and oligodendrocytes ^62^, underscoring its relevance to glial support functions. Since *SOX2* regulates oligodendrocyte maturation and myelin repair, while *NR4A2–EGR1* interactions shape activity-dependent transcription in deep-layer excitatory neurons, the convergence of these TFs on shared targets suggests that MDD-associated dysregulation spans both neuronal and oligodendrocyte-lineage regulatory programs. In this framework, *NR3C1* could provide a canonical stress-hormone transcriptional entry point (including regulation of downstream stress-responsive genes such as *SGK1*) ^45^, *MYRF* could serve as a core regulator of the oligodendrocyte-lineage myelination program ^51^, and *NEUROD1* could modulate neurogenic differentiation and maturation ^54^ (Fig. 3e).

To summarize, we identified key TFs that are computationally inferred to differentially regulate *ABHD17B* in excitatory neurons from female donors. In combination with their TGs, these TFs govern response to stress, synaptic homeostasis, plasticity-driven myelination, and neural maintenance. While these findings provide a compelling hypothesis for MDD neuropathology, they remain predictive and warrant further experimental investigation.

### ECLARE captures developmental trajectory in unpaired data

Neurodevelopment in the human cortex is characterized by a series of regulatory gene programs and has been studied through the lens of single-cell gene expression and chromatin accessibility ^36;63;64^. Although clear insights have been drawn from investigating paired multiomics data from human cortex at different stages of development^63;64^, performing similar analyses on unpaired data is more challenging^36^. We thus turned to the integration of unpaired data collected in the context of studying the developmental trajectories of brain cell types^36^. This integration task would demonstrate how `ECLARE` captures transitions between cell types along the developmental axes. To this end, we used **PFC_Zhu** ^63^ and **PFC_V1_Wang** ^65^ paired datasets to train teachers, which then guided the `ECLARE` student to integrate the unpaired **CTX_Velmeshev** ^36^ dataset (Table 2). Note that we limited our analysis to excitatory neurons and inhibitory interneurons, as these abundant neuronal populations exhibit well-resolved developmental trajectories and lineage branching patterns in Velmeshev et al. ^36^.

By visualizing integrated nuclei in the form of force-directed layouts using Scanpy’s implementation of ForceAtlas2 (FA), we observe a separation of major excitatory and inhibitory neuronal lineages, as well as a finer aggregation into cell types (Fig. 4a, top row). Quantitatively, `ECLARE` achieves superior performance compared to scJoint and GLUE on this unpaired **CTX_Velmeshev** data in terms of biological clustering and multi-omic integration (Fig. 4b), thereby providing better nuclei embeddings for downstream multi-omic analysis (Fig. S7).

**Figure 4:**
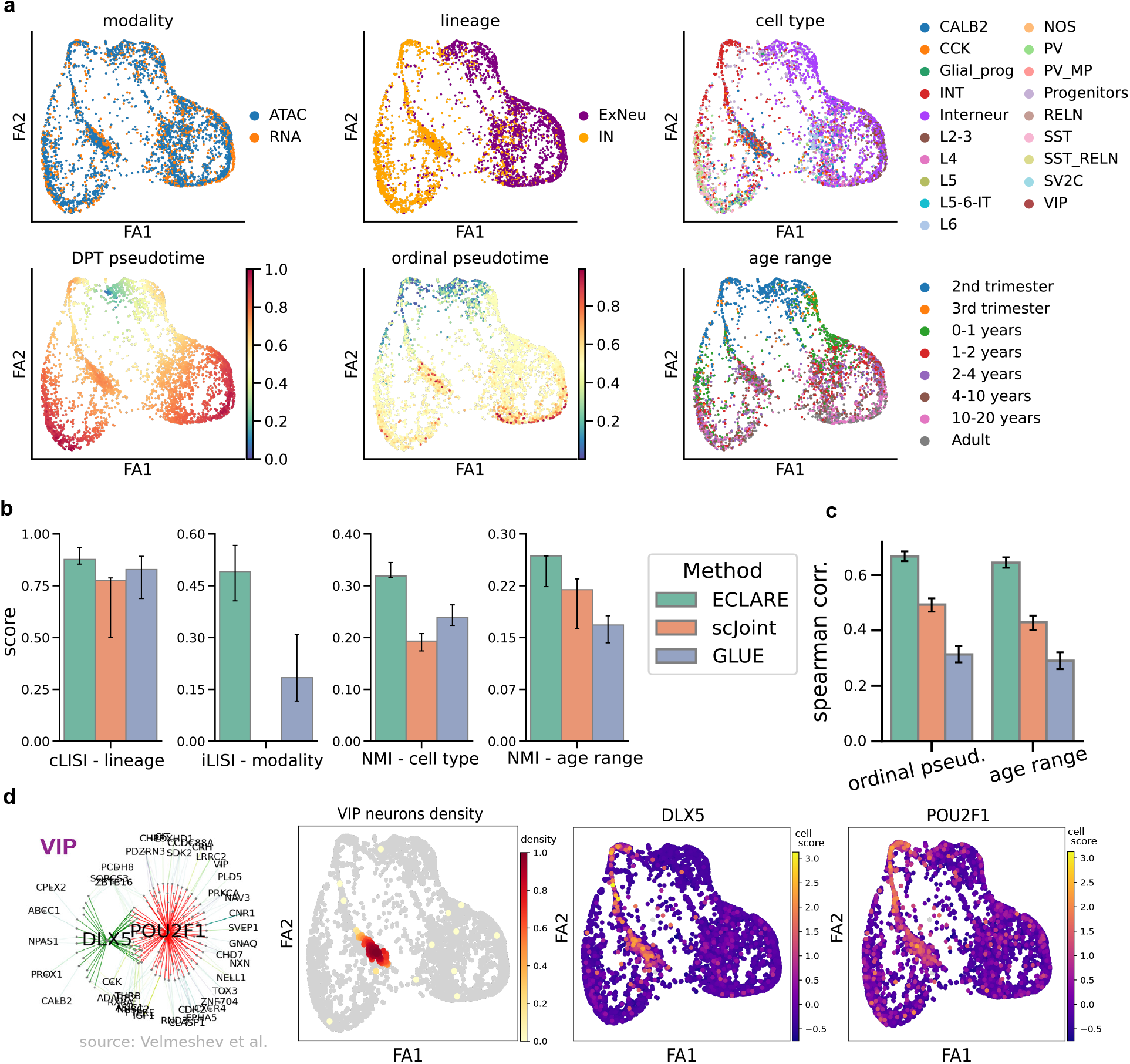
Integration of unpaired developmental data. **(a)** `ECLARE` embeddings visualized with force-directed embedding using ForceAtlas2 (FA). **(b)** Biological clustering and multi-omic integration metrics derived from latent embeddings for `ECLARE`, scJoint and GLUE. Error bars were obtained via bootstrapping nuclei (n=500). **(c)** Spearman correlations between DPT pseudotime and either ordinal pseudotime or age range. Error bars were obtained via bootstrapping nuclei (n=500). **(d)** Cell scores for VIP eRegulons inferred by Velmeshev et al. ^36^ using SCENIC+. The first panel is taken directly from the original publication to highlight the importance of *DLX5* and *POU2F1* eRegulons for VIP neurons. The second panel shows the density of VIP neurons within the `ECLARE` embedding. The third and fourth panels show cell scores for *DLX5* and *POU2F1* eRegulons, respectively.

We also performed unsupervised pseudotime inference using partition-based graph abstraction (PAGA)^66^ followed by diffusion pseudotime (DPT)^67^, as well as supervised pseudotime inference using an ordinal regression model based on CORAL^68^ that was trained on **PFC_V1_Wang** and applied onto **CTX_Velmeshev** (Fig. 4a, bottom row; Fig. S8). These inferred pseudotime values enabled us to treat DPT pseudotime as our *predicted* pseudotime and compare these predictions to a continuous reference variable (predicted ordinal pseudotime, Fig. S8) in addition to ground-truth ordinal labels (age range, Fig. S9). By computing Spearman correlations of DPT pseudotime with either ordinal pseudotime or age range, we observe that `ECLARE` produces cell embeddings in stronger agreement between predicted DPT pseudotime and both reference variables compared to scJoint and GLUE (Fig. 4c). These results demonstrate that `ECLARE` provides well-integrated embeddings that split the major neuronal lineages apart while also capturing the developmental axes with high fidelity.

We then set out to verify whether the developmental axes captured by `ECLARE` can reveal cell type-specific multi-omic patterns identified in the original publication for **CTX_Velmeshev** ^36^. To this end, we paired nuclei across RNA and ATAC modalities within the integrated latent space using the two-step OT procedure. We used the SCENIC+ eRegulon signatures from Velmeshev et al. ^36^ identified for the VIP cell type to perform cell-scoring on the paired nuclei. Indeed, cell scores of VIP-related *DLX5* and *POU2F1* eRegulons are enriched in high-density VIP regions of the `ECLARE` embedding (Fig. 4d). Moreover, these eRegulons are also enriched in the developmental branch stemming from the root and terminating at the high-density VIP region.

The enrichment of *DLX5* and *POU2F1* eRegulon activity in high-density VIP regions and along the corresponding developmental branch is consistent with known interneuron differentiation programs. *DLX5* is a key regulator of GABAergic interneuron development, expressed in CGE-derived progenitors and maintained in mature VIP neurons ^69;70^. Its eRegulon activity likely reflects progression through DLX-driven differentiation states. *POU2F1*, while not interneuron-specific, shows enriched motif accessibility in developing human neurons ^71;72^, suggesting it supports broader transcriptional programs active during early VIP neuron differentiation. Together, their eRegulon enrichment along the VIP trajectory marks transcriptional transitions characteristic of CGE-derived interneuron maturation.

### Knowledge distillation enables co-embedding of developmental and MDD data

Although diagnosed later in life, MDD is shaped by early developmental windows extending from prenatal life into early adulthood with known cellular mechanisms. These mechanisms relate genetic variants with neuronal hyperexcitability induced by childhood adversity and environmental exposure to stress ^73^. Moreover, regulatory effects of MDD GWAS variants using snATAC-seq data revealed disruption of many neurodevelopment-related TF binding sites in excitatory neurons ^29^. Previous studies have used multi-omics data acquired at different developmental stages to interrogate molecular associations between gene expression and chromatin accessibility features related to MDD variants ^63;65^. However, such findings are not confirmatory due to the nature of the data, i.e. the lack of clinical diagnoses during earlier developmental stages. We investigated whether co-embedding of developmental and MDD data using our `ECLARE` framework could guide the organization of the learned embedding towards longitudinal processes in MDD (such as gene programs related to development) as opposed to merely clustering around cell-type annotations (e.g. Fig. 3a). Specifically, we trained a developmental teacher based on paired developmental data (**PFC_Zhu**) and performed single-teacher KD (i.e. KD-CLIP) to train an MDD student model based on this developmental teacher (Fig. 5a.i-ii). By leveraging the single-teacher strategy, we co-embed the paired developmental data and MDD data via the same MDD student model (Fig. 5a.iii-iv). To simplify our analysis while ensuring a sufficient number of nuclei for statistical inference, we focused our analyses on excitatory neurons (ENs), which we prioritized in light of known disruptions of neurodevelopment-related TF binding sites in this cell population ^29^.

**Figure 5:**
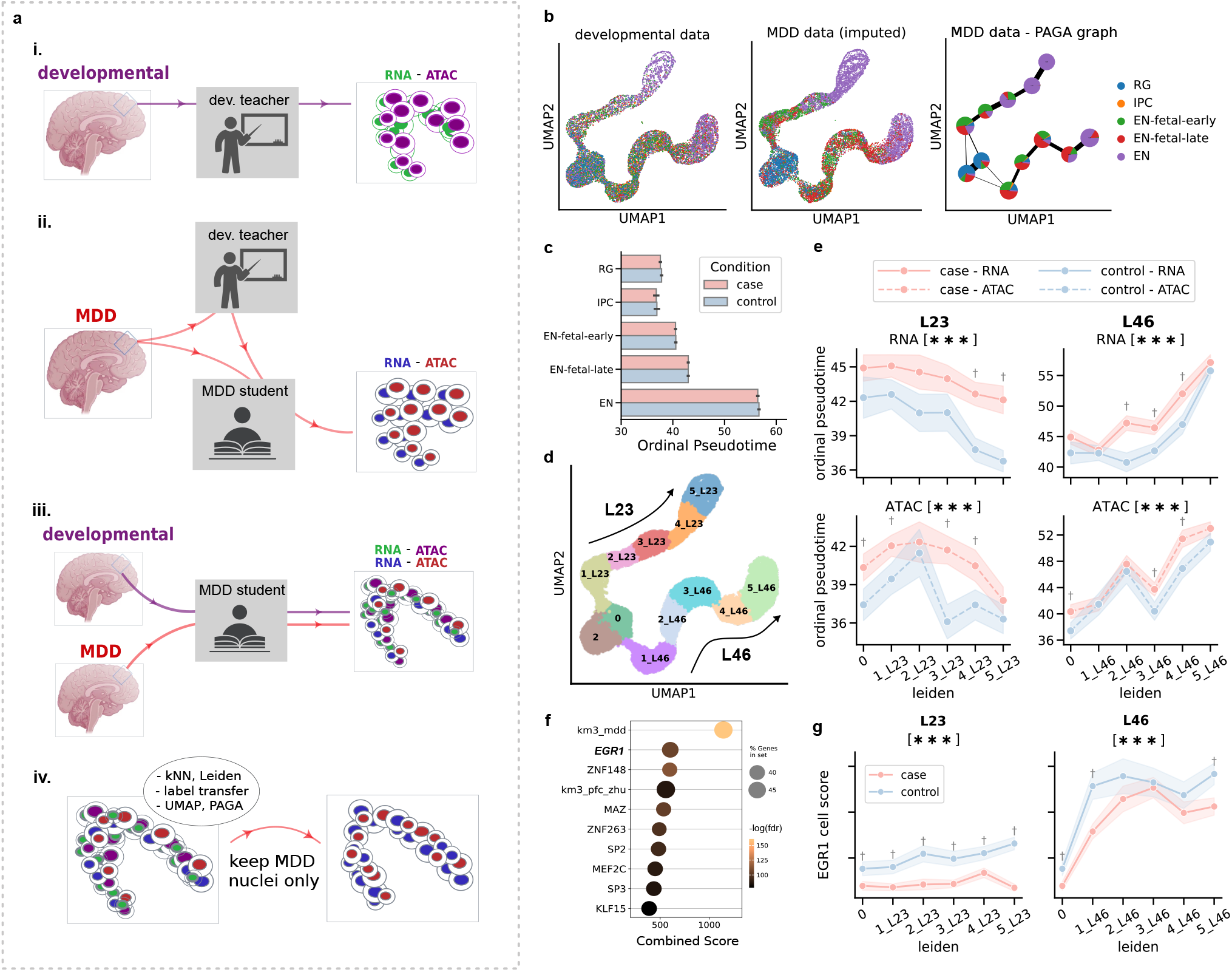
Uncovering longitudinal processes in MDD via co-embedding with developmental data. **(a)** Four-step process for co-embedding nuclei from MDD and developmental datasets: **i**. train a teacher model using developmental data (**PFC Zhu**); **ii**. train a student model using the developmental teacher model and MDD data; **iii**. embed nuclei from both developmental and MDD datasets using the same MDD student model; **iv**. after performing processing steps of interest on co-embedded nuclei (label transfer, PAGA, etc.), discard nuclei from developmental data to only keep MDD data nuclei. **(b)** UMAP of co-embedded developmental and MDD data based on imputed developmental cell-type labels, alongside PAGA graph for MDD data. **(c)** Mean ordinal pseudotime of MDD data per imputed cell-type label and MDD condition label. The y-axis contains the developmental cell types from early to latest. RG: radial glia; IPC: intermediate precursor cells; EN: excitatory neurons. **(d)** UMAP visualization of MDD nuclei colored by Leiden cluster and labeled according to progression along two longitudinal branches L23 and L46, both originating at cluster 0. **(e)** Female-specific analysis of case vs control differences in ordinal pseudotime per Leiden cluster organized along the two longitudinal branches. Results are organized by modalities (rows) and longitudinal branches (columns), where branch-level significance is annotated in sub-panel title and cluster-level significance is annotated above data points. *\* * \** denotes *p <* 0.001 whereas *†* denotes *p <* 0.05. **(f)** EnrichR-based enrichment of differentially-expressed genes in excitatory neurons from female donors, based on gene sets derived from brainSCOPE TF regulons and longitudinal gene clusters. Emphasis added on *EGR1* to ease visual retrieval. **(g)** Case and control multi-omic cell regulatory scores for *EGR1* eRegulons across Leiden branches, restricted to nuclei from female donors. The significance of results is reported as in panel **e**.

### MDD is associated with later developmental signatures in female donors

To characterize the organization of the learned embedding with respect to the alignment of MDD data nuclei to developmental data nuclei and their ordinal pseudotime values, we used co-embedded nuclei (Fig. 5a.iv) to impute developmental cell-type annotations onto the MDD data using k-Nearest Neighbor (kNN) label-transfer (Fig. 5b). Indeed, we confirm that MDD nuclei are organized along two branches that follow the expected progression between imputed cell types: *i*. radial glia (RG), *ii*. intermediate precursor cells (IPC), *iii*. early fetal ENs (EN-fetal-early), *iv*. late fetal ENs (EN-fetal-late) and *v*. mature post-fetal ENs (EN). This progression is also consistent with the increase in ordinal pseudotime values across imputed cell-types, which were assigned based on ATAC and RNA CORAL models trained on **PFC Zhu** (Fig. 5c; Fig. S10). Mean values were combined across modalities, branches and sexes, but separated by condition. Note that MDD nuclei were well balanced by donor age such that any differences in ordinal pseudotime could be attributed to CORAL outputs rather than being confounded by donor age (Fig. S11).

In light of the lack of case-vs-control differences in broadly-aggregated ordinal pseudotime values (Fig. 5c), we performed a more detailed dissection by separating ordinal pseudotime values per modality, branch and sex. For this dissection, the imputed cell-type labels were discarded and replaced by Leiden clusters specified along two longitudinal branches (Fig. 5d). These two branches were labeled by the majority cell subtypes comprising each branch based on MDD RNA data annotations ^28;35^, giving rise to the L23 and L46 branches. Of note, the L46 branch contains a higher density of **PFC_Zhu** than the L23 branch whereas the density distribution for MDD nuclei is uniform across branches, indicating that the L46 branch contains a higher density of co-embedded nuclei between datasets (Fig. S12).

By computing the mean ordinal pseudotime per Leiden cluster, we observed in female donors that the ordinal pseudotime for both modalities is consistently greater in cases compared to controls for all branches and modalities, with some significant cluster-specific differences as well (Fig. 5e). This consistent increase in ordinal pseudotime in MDD female cases suggests a shift toward later developmental signatures in this cohort, echoing reported convergent effects of aging and psychiatric disease ^74^ and the use of epigenetic aging clocks in neuropsychiatry research^75^. Results for male donors are not shown due to statistical insignificance (Fig. S13).

### Targets of *EGR1* are differentially expressed along longitudinal processes

To reveal whether neuropsychiatric risk variants are enriched in particular developmental stages, Zhu et al. ^63^ mapped risk variants to developmentally-associated gene clusters and their linked peaks (i.e. Fig. 5 in Zhu et al. ^63^). We repeated a similar set of analyses on excitatory neurons, where the **PFC_Zhu** and MDD datasets were used separately to cluster genes into four (pseudo)temporally-enriched clusters using k-means (km) clustering. These pseudotemporal clusters were labeled for each dataset separately based on the relative position of their genes’ maximal expression along the pseudotime axis, from earliest (**km1_mdd_&_km1 pfc_zhu**) to latest (**km4_mdd_&_km4_pfc_zhu**) expression (Fig. S14). We then examined whether any of these pseudo-temporal gene clusters were over-represented by TF regulons (i.e. sets of target genes affected by the same TF) identified as differentially expressed using pyDESeq2^76^. In female donors, we found that DEGs overlapped significantly with the regulons of *EGR1* as well as both **km3_mdd** and **km3_pfc_zhu** pseudotemporal clusters (Fig. 5f). Moreover, genes overlapping **km3_mdd** and the set of *EGR1* regulons are statistically over-represented in the ‘GO Biological Process 2021’ pathways associated with neuronal activity, including terms such as ‘regulation of neurotransmitter receptor activity’ and ‘regulation of neuronal synaptic plasticity’ (Fig. S15). Taken together, the fact that DEGs are over-represented in clusters of pseudotemporally-correlated genes (i.e. **km3_mdd_&_km3_pfc_zhu**) and comprise of many targets of *EGR1* (Fig. 5f) suggested that the differential expression of *EGR1* targets may follow a longitudinal pattern. Moreover, our previous analysis using sc-compReg on paired MDD nuclei pointed to *EGR1* as a key TF that exhibits differential gene regulation on its target genes in the context of MDD (i.e. Fig. 3e).

To further characterize the differential enrichment of *EGR1* as a potential TF mediator of MDD, we verified that the regions containing EGR1 binding motifs were differentially accessible using chromVAR^77^ at the nominal p-value *<* 0.05. We also performed cluster-specific differential analysis using a subset of peaks corresponding to each pseudotemporal “km” gene clusters. Besides the *EGR1* motifs, significant motifs for *AR* and *NR3C1* (or glucocorticoid receptor, GR) were identified, based on peaks linked to **km2_mdd** genes (Fig. S16f). *EGR1* is a direct transcriptional regulator of the *NR3C1* gene and early-life environment can program the methylation state of this *EGR1* -binding site in the *Nr3c1* promoter in rodents, altering lifelong GR expression ^78^. Moreover, our previous MDD GRN analysis identified *NR3C1* /*GR* as a key TF involved in MDD in excitatory neurons from female donors (Fig. 3e), in addition to our enrichment results based on sc-compReg candidate genes and *Brain*.*GMT* gene sets identified chebotaev gr targets up as being differentially regulated in MDD (Fig. S2). Finally, the TRRUST v2 database^79^ identifies the *AR* gene as one of two targets of *EGR1* that is associated with mental depression.

Because the DEG enrichment results point towards longitudinal expression patterns among *EGR1* targets (Fig. 5f), we next examined whether expression of *EGR1* target genes and accessibility of their linked regulatory elements exhibit coordinated pseudotemporal variations in MDD. To this end, we paired ATAC and RNA nuclei from MDD data and computed multi-omic cell scores for an *EGR1* regulon derived from the neurodevelopmental SCENIC+ eRegulons provided by Velmeshev et al. ^36^. Indeed, we observe in female donors a case-vs-control separation in *EGR1* cell scores in both branches (Fig. 5g), which is absent when analysing male donors (Fig. S17). Of note, the increase of *EGR1* cell scores along branch L46 follows the increase in ordinal pseudotime along this same branch (Fig. 5d).These observations are consistent with previous findings that *EGR1* is differentially regulated in MDD with variations between age groups ^80–83^.

In sum, by training a student model guided by a developmental teacher model and co-embedding developmental and MDD data via the `ECLARE` framework, we identified two longitudinal branches for excitatory neurons that expose condition-based divergences in ordinal pseudotime. These divergences are accompanied by longitudinal patterns in *EGR1* cell scores demarcating MDD cases and controls in excitatory neurons from female donors.

## Discussion

Integrating scRNA-seq and scATAC-seq promises to identify context-specific gene regulatory networks (GRN) by linking TFs with open chromatin regions and their downstream genes. This will elucidate dysregulated GRN in disease, but progress is hindered by computational methods that can leverage the limited paired multiome data to integrate the vast amount of unpaired multiome data. In this study, we developed `ECLARE` as a means of leveraging paired datasets via contrastive learning (CL) and knowledge distillation to align unpaired modalities by optimizing optimal transport (OT) and CL loss (Fig. 1). We demonstrated the effectiveness of `ECLARE` by first establishing that a simple MLP trained with CL loss serves as a good base model for zero-shot alignment tasks, achieving superior alignment and clustering metrics than other integration methods (Fig. 2a). We then demonstrated the improvement of multi-teacher KD over the zero-shot alignment on both paired (Fig. 2b) and unpaired MDD (Fig. 2c) datasets.

To showcase `ECLARE` in real-world applications, we carried out three related case studies centering around MDD and brain development & aging. First, in aligning unpaired MDD multiomes, we observed broad patterns of MDD-specific enrichment involving excitatory neurons and oligodendrocytes (Fig. 3b-d). Then, based on sc-compReg differential regulation analysis, we observed that multiple TFs including *EGR1* show differential regulation of *ABHD17B* in excitatory neurons sourced from female subjects (Fig. 3e). These TFs govern response to stress, synaptic homeostasis, plasticity-driven myelination, and neural maintenance. Second, we demonstrated that `ECLARE` can learn integrated representations of unpaired data collected at different stages of development and aging in human cortical tissue (Fig. 4). We incorporated the concept of ordinal pseudotime as a means of examining whether the learned representations conform to longitudinal processes, making novel use of the CORAL framework for single-nucleus data. Compared to scJoint and GLUE, `ECLARE` learns representations that better capture developmental and aging-related structure in human cortex. Lastly, the above two biological case studies on MDD and development paved the way for the third case study on a co-embedding analysis where developmental data were used by `ECLARE` to integrate MDD data anew and expose longitudinal patterns relating to MDD (Fig. 5). Focusing on excitatory neurons from female donors, we show that `ECLARE` learns an embedding in which MDD cases are shifted toward later developmental signatures relative to controls. From these pseudotemporal analyses on MDD data, we further highlighted *EGR1* as a transcription factor of interest, whose target genes and linked peaks exhibit coordinated (pseudo)temporal activity and differential expression in excitatory neurons from the female donors.

A key limitation of the current framework is its reliance on pairwise feature alignment between source and target datasets. While `ECLARE` avoids the severe feature loss of global all-to-all intersection, the required pairwise intersection still discards some dataset-specific regulatory elements. Several other limitations concerning data heterogeneity and practical applicability remain. First, projecting adult disease samples onto a fetal/adolescent developmental reference introduces a substantial domain shift. This co-embedding may be susceptible to batch effects or reference bias, as developmental and adult trajectories are driven by distinct transcriptomic programs. Second, the practical application of `ECLARE` requires selecting appropriate paired multi-omics training data. While we demonstrate that models trained on related tissues generalize well, performance may degrade if the target data lacks a suitable paired reference. In scenarios where paired training datasets of the same tissue origin are unavailable, users may need to rely on alternative prior-based methods or explore cross-tissue generalization strategies. Finally, our benchmarking and biological case studies were restricted to brain tissues, limiting the scope of our claims and findings to this tissue type.

To extend `ECLARE`, we will explore several future directions. First, we can weight different teachers as a function of cell-type or class imbalances, and implement more sophisticated architectures such as Transformer^16^ to increase model capability and transferability. Second, `ECLARE` can be adapted to other multi-omics datasets that further incorporate other modalities such as methylomics, proteomics and spatial data. It is also worth investigating whether `ECLARE` would further benefit from a mosaic integration approach, where datasets are only partially unpaired and thus contain overlapping cells and/or features ^1^. Third, other training paradigms can be incorporated to further boost the robustness of `ECLARE`. For example, we can take into account distributional differences between the reference paired and unpaired target data during KD^84^. Lastly, GRN analysis can be expanded to identify specific peak-gene linkages. Similar to GLUE^12^, prior knowledge about feature associations such as in-cis regulatory interactions could be incorporated within the training paradigm for more interpretable results. Standard scATAC-seq processing libraries such as ArchR^18^ and Signac ^85^ can infer peak-gene links, facilitating the use of such prior information for developing better diagonal integration methods.

In conclusion, we demonstrate that KD is a useful paradigm for improving single-nuclei integration quality beyond zero-shot transfer on paired and unpaired multi-omic data. Importantly, the proposed `ECLARE` incorporating multi-teacher KD achieves the best overall multi-omic alignment and enables in-depth analysis of the differential and temporal GRN in MDD. With increasing single-cell multiomic data, we envision `ECLARE` to be an attractive solution for large-scale functional genomic analysis.

## Methods

### The ECLARE framework

We aim to achieve diagonal integration of unpaired single-cell/single-nucleus (sc/sn) RNA-seq (sc/snRNA-seq) and sc/snATAC-seq data. For this task we introduce `ECLARE`, an overview of which is provided in Fig. 1. Note that we will refer to cells and nuclei interchangeably.

#### Contrastive pretraining of individual teacher models

The first step of `ECLARE` (Fig. 1a) is to independently train a set of teacher models on their respective paired RNA & ATAC multi-omics data (i.e. the *source* datasets). These teacher models are trained based on the CLIP framework by minimizing the InfoNCE loss ^30^:

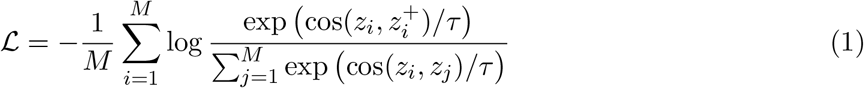

where *z*_*i*_ ∈ {*a*_*i*_, *r*_*i*_} represents the embedding of a given anchor cell from one modality (i.e. RNA cell *r*_*i*_ or ATAC cell *a*_*i*_), 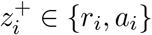 represents the matching cell embedding from the other modality, *z*_*j*_ represents any cell from the modality opposite to the anchor (including 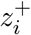), ‘cos’ represents the cosine similarity, *τ* the temperature hyperparameter (*τ* =1 for our experiments), and *M* the total number of cells for a given batch. The InfoNCE loss optimizes the encoders so that each matched pair has higher similarity relative to all mismatched pairs (i.e., the diagonal dominates under softmax-normalization). In this way, a given teacher model is trained to learn the alignment between matching RNA and ATAC cells leading to an integrated latent space, where pairs of matching cells share highly similar coordinates. Importantly, the encoders for each teacher can operate on a different set of features (i.e., genes and peaks) from each dataset and no feature harmonization is needed across source datasets.

#### Multi-teacher knowledge distillation

The second step of `ECLARE` is to distill the knowledge from this ensemble of trained teacher models into a student model using a separate *target* dataset (Fig. 1b). Importantly, this target dataset can be unpaired; this second step thus relies on the ability of the teacher models to perform multimodal alignment in a zero-shot fashion without relying on cell-pairing information. This knowledge distillation (KD) step is performed by projecting the cells or nuclei from the target dataset through the encoders of the teacher and student models to construct cell-cell similarity matrices for each model. From these similarity matrices, two loss terms are computed for each student-teacher pair: a KD loss (Fig. 1c) and an alignment loss (Fig. 1d). These two losses are combined into a single total loss via a convex mixture (Fig. 1b), similar to the mixture of soft-labels and hard-labels losses initially proposed for KD^24^ or the mixture of KD and InfoNCE loss from MobileCLIP^86^:

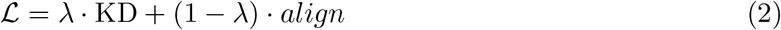

where *λ* is a hyperparameter that controls the trade-off between mimicking the teachers’ representations (the KD loss) and explicitly enforcing cross-modality alignment via optimal transport (the alignment loss). A higher *λ* prioritizes structural preservation from the teachers, while a lower *λ* emphasizes refining the cross-modal matching. Note that since the teacher model is used to project both source dataset during training and target dataset for inference, pairwise feature alignment is still required between source and target datasets. We perform this pairwise ATAC peak set alignment using intersection. While this intersection-based approach inevitably discards some dataset-specific regulatory elements, it is a necessary trade-off to enable zero-shot transfer from teacher to student. However, because `ECLARE` only requires pairwise alignment (source-to-target) rather than global all-to-all alignment across all source datasets simultaneously, it mitigates the severe feature loss that typically occurs when intersecting many datasets. We acknowledge that the reliance on pairwise feature alignment remains a limitation of the current framework.

For the KD loss (Fig. 1c), softmax is computed for all similarity matrices to obtain probabilities across rows and columns, such that the Kullback-Leibler (KL) divergence can be computed between the rows and columns of the student similarity matrix with those of each teacher similarity matrices. The KL divergences for RNA (Eq. 3) and ATAC (Eq. 4) are then summed to obtain a single KD loss per teacher (Eq. 5):

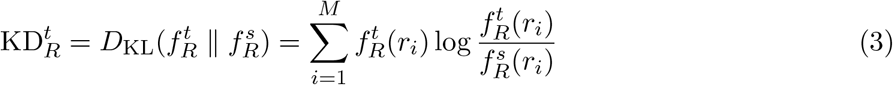

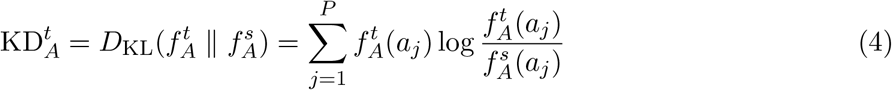

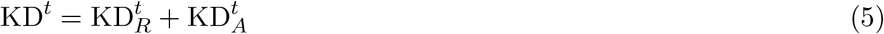

where 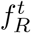 and 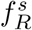 are the distributions obtained by computing row-wise softmax on the teacher and student cosine similarity matrices, 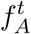 and 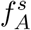 are the distributions obtained by computing column-wise softmax on the teacher and student cosine similarity matrices, and *M* and *P* are the number of RNA and ATAC nuclei (generally, *M* = *P* during mini-batch training).

For the alignment loss (Fig. 1d), the teacher similarity matrix is refined by solving a classical exact Optimal Transport (OT) problem using the *ot*.*solve()* function from the Python OT library (`POT` ^87^). This is done by minimizing the Wasserstein distance between the scRNA-seq cells and scATAC-seq cells based on the cosine similarity computed by the teacher CLIP model:

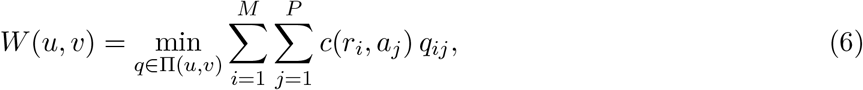

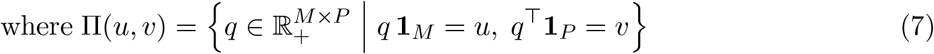

where *c* corresponds to the cosine distance defined as *c* = 0.5[1 − cos(*x, y*)]. The constraints in Eq. (7) indicate that the elements of *q* (the *transport plan*) are positive-definite real numbers, and that summing across rows and columns matches the marginals *u* and *v*, which we both set as uniform distributions for simplicity. Also, *r*_*i*_ and *a*_*j*_ correspond to embedded RNA and ATAC cells.

Because we solve a balanced, unregularized OT problem using a linear program solver on square matrices with uniform marginals ^88^, this process sparsifies the similarity matrix into a scaled permutation matrix (i.e. the transport plan *q*) that defines unique pairs of cells across modalities. This teacher transport plan is then used as ground truth to compute the InfoNCE loss from the student similarity matrix, normalized via the softmax operation. By treating the teacher transport plan *q* as a bivariate teacher distribution and the student RNA and ATAC softmax distributions 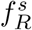 and 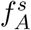, the alignment loss is computed with cross-entropy:

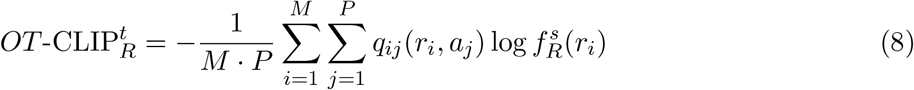

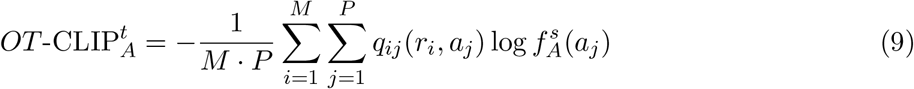

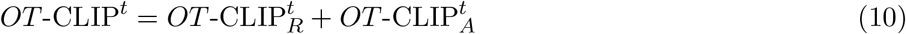

In other words, we compute the InfoNCE loss similarly to CLIP, but compute cross-entropy for entries of the student similarity matrix that correspond to the teacher transport plan rather than simply using the diagonal entries. Since we solve for an OT problem to obtain the teacher transport plan and then compute a loss similar to that of CLIP, we call this loss variant the *OT* -CLIP loss.

#### Computing teacher-specific weights

Both the teacher-specific KD and alignment losses from Eqs. (5) & (10) are combined across teachers to create ensemble multi-teacher KD and alignment losses via weighted averaging (Fig. 1b).

For KD, we use the teacher similarity matrices *C*^*t*^ ∀ *t* ∈ {1, …, *T*} to quantify how well a given teacher is capable of aligning nuclei across modalities, where teachers with higher cosine similarity values in *C*^*t*^ see their KD loss upweighted, and vice-versa. Specifically, consider *C*^*t*^ and the transport plan *q* derived from *C*^*t*^ via Eq. (6). The values in *C*^*t*^ that intersect with the predicted matches in *q* – i.e. wherever *q >* 0 – can be extracted as follows:

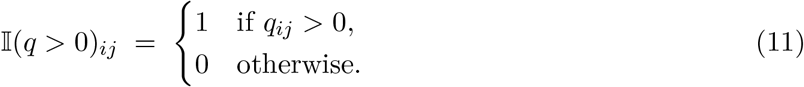

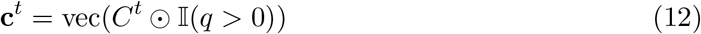

where **c**^*t*^ represents a vector of cosine similarity values for the pairs predicted by teacher *t*. On the one hand, we can obtain weights based on the average value across the *M* pairs of nuclei as follows:

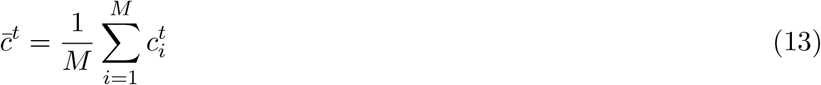

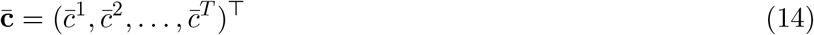

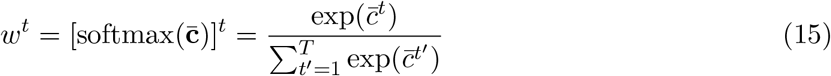

which provides a single weight for each teacher that is shared between all pairs of nuclei. On the other hand, the averaging across pairs from Eq. (13) can be withheld, giving rise to a weight for each teacher at each pair, indexed by e.g. RNA nuclei *i* ∈ {1, …, *M*}:

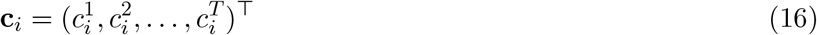

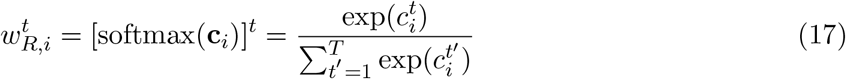

such that 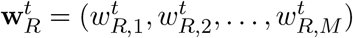. In the same fashion, one can derive weights by indexing the ATAC nuclei *j* ∈ {1, …, *P*}, leading to teacher weights 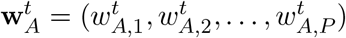.

Because a teacher weight is defined for each RNA and ATAC nucleus, the KD losses defined in Eqs. (3)–(5) can instead be reweighted as follows:

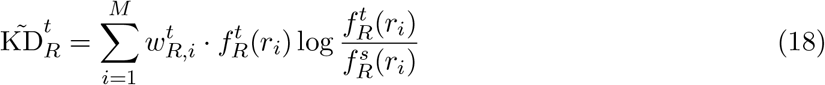

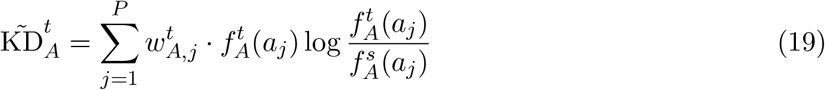

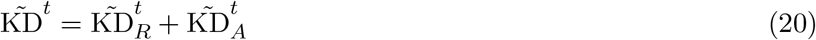

We adopt this nucleus-level teacher weighting in `ECLARE` and obtain the ensemble multi-teacher KD loss as follows:

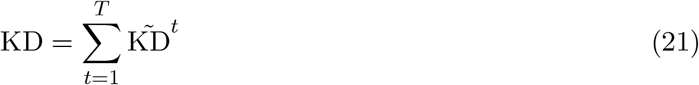

Lastly, the *OT* -CLIP alignment losses also require their teacher-specific weights. For computing these weights, we extract for each teacher the linear Wasserstein distance associated with solving the OT problem (Eq. 6) that is provided by the solver in the POT library, giving rise to a Wasserstein distance *l*^*t*^ for teacher *t*. Teacher weights are then obtained by computing *v*^*t*^ = softmax(1 − *l*^*t*^) for teachers *t* ∈ {1, …, *T*}. Note that teacher reweighting of the alignment losses is performed for the whole batch of nuclei, such that the unweighted teacher alignment losses from Eq. (10) are weighted and summed:

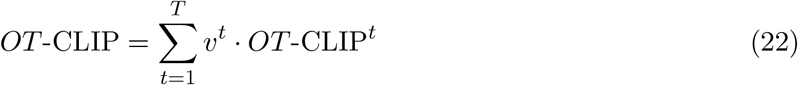

Finally, the ensemble KD and *OT* -CLIP losses from equations Eqs. (21) & (22) are combined into the total loss of Eq. 2.

#### CLIP, KD-CLIP and ECLARE model training

We performed hyperparameter tuning using Optuna^89^ with TPE sampler and median pruner to find optimal parameters for the training parameters and architectures for CLIP, KD-CLIP and `ECLARE` models. We designed a compound metric to use as the objective, corresponding to a weighted average of iLISI, ARI and NMI metrics computed on held out nuclei in the CLIP embedding:

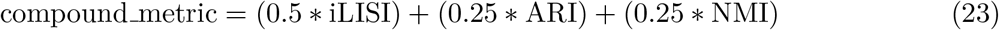

Refer to the “Evaluation metrics” section for information on these metrics. This compound metric incorporates measures of multi-omic integration (iLISI) and biological clustering (ARI & NMI).

The hyperparameter tuning experiments involved training CLIP models on the primary paired datasets (Table 1), zero-shot projection of MDD data nuclei into the CLIP models’ respective embeddings and then computing the compound metric on these projected nuclei. Once the ideal hyperparameters for the CLIP models were identified, we performed the same procedure on the `ECLARE` MDD student model, where the CLIP teachers now have fixed parameters. One noteworthy parameter introduced by the student model is the *λ* parameter used for computing the total loss in Eq. 2.

**Table 1:** Hyperparameters and Tested Values.

| Hyperparameter | Distribution / Tested Values | Selected Values |
| --- | --- | --- |
| `num_units` | Categorical: {128, 256, 512} | 256 |
| `_teacher_num_layers` | Categorical: {1, 2, 3} | 1 |
| `_student_num_layers` | Categorical: {1, 2, 3} | 2 |
| `dropout_p` | Uniform (Float): [0.1, 0.9] | 0.3 |
| `distil_lambda` | Categorical: {0, 0.1, 0.25, 0.5, 0.75, 0.9, 1} | 0.1 |

**Table 2:**
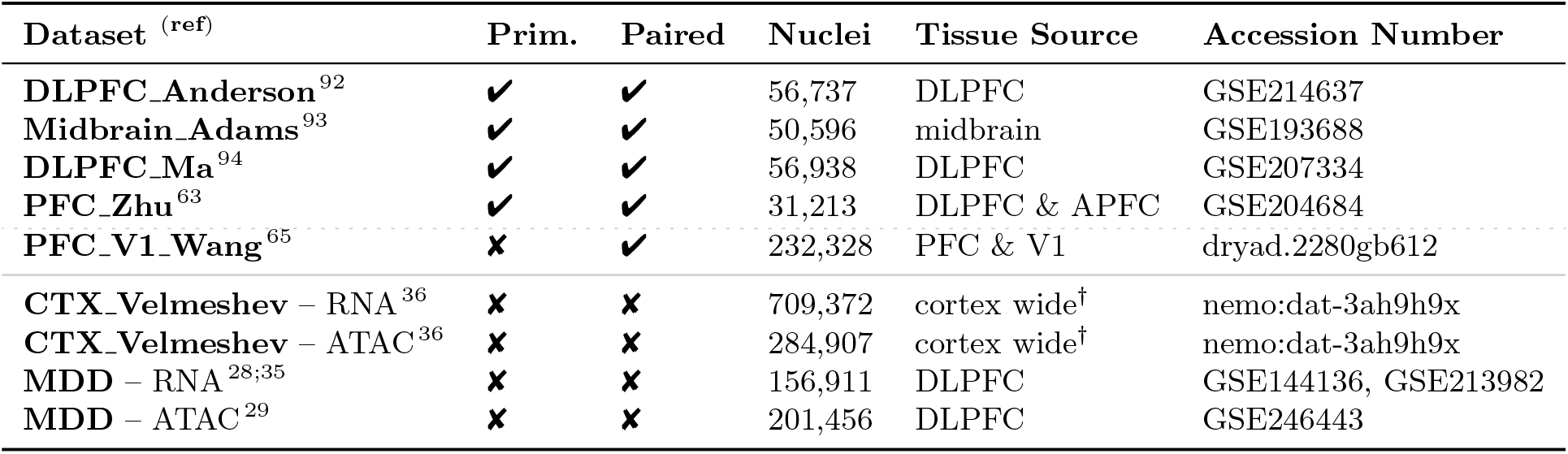
Summary of datasets. The “Prim.” column identifies paired datasets that compose of the main paired datasets used for benchmarking experiments. The solid horizontal dividing line delineates paired (above) from unpaired (below) datasets. The dashed horizontal dividing line delineates primary paired datasets from the paired **PFC_V1_Wang** dataset. DLPFC: dorsolateral prefrontal cortex, APFC: anterior prefrontal cortex, V1: primary visual cortex, PBMC: peripheral blood mononuclear cells. ^*†*^ includes prefrontal, cingulate, temporal, insular & motor cortices

The tested and selected values are listed below in Table 1, which are the default parameters used throughout this report. For simplicity, we prioritized keeping the same values for teacher and student models, with the exception of the number of hidden layers since this parameter showed maximal benefit when allowed to differ for teacher and student models. Note that all models adopt a standard MLP architecture with ReLU units and dropout layers.

All models were trained in PyTorch ^90^ using AdamW optimization with a learning rate of 0.001, weight decay of 0.01 and batch size of 800 nuclei per modality.

### Single-nucleus multi-omics data

We make use of several multi-omics snATAC-seq and snRNA-seq datasets from brain tissue that are summarized in Table 2. Of these datasets, five are publicly-available *paired* datasets acquired using snMultiome or similar technologies and thus provide direct matching information between nuclei across both modalities. Of these five paired datasets, the first four listed in Table 2 serve as the primary paired datasets used throughout benchmarking experiments. The fifth paired dataset **PFC V1 Wang** is only used for the developmental biological analysis, not for benchmarking nor for training MDD student models. The last four datasets listed in Table 2 correspond to *unpaired* datasets. RNA and ATAC **CTX Velmeshev** datasets are used as the target for the developmental biological analysis. The RNA and ATAC **MDD** datasets are from case and control subjects of a prior study on MDD for which no direct pairing information is available, thus requiring diagonal integration to perform downstream analysis.

#### Data preprocessing

For each dataset listed in Table 2, we performed the standard preprocessing steps of extracting protein-coding genes from RNA data, removing features with zero counts, library size normalization (*target_sum*=1e4), log(1+*x*) transformation and min-max scaling (*max_value*=10). Also, for experiments focused on model training, benchmarking and quantitative assessment of performance with metrics, we used highly-variable feature selection for both RNA and ATAC data using Scanpy’s *highly_variable_genes* function to reduce the number of input features and consequently the total number of parameters in CLIP models. This feature selection step was performed before min-max scaling, and was used to set the maximum number of genes to 10,000 and maximum number of peaks to 100,000. To further limit the number of input features, we only kept peaks that are within a 1Mbp window around protein-coding genes.

#### Pairwise alignment of multi-omic datasets

Feature alignment between source and target datasets is required to enable forward passes with a teacher model. Thus, each source-target combination requires a pairwise feature alignment for both RNA gene features and ATAC peaks. Hence, we keep the intersection of genes and peaks shared across source and target datasets. Whereas the gene intersection can be straightforwardly identified based on gene name, ATAC peaks requires the *intersection* utility from pybedtools ^91^ based on the peaks’ intervals. Afterwards, every peak from the source dataset that is part of this intersection is relabeled based on its homologous peak in the target dataset. In the case that multiple source peaks map shared the same target peak, only one of the source peaks was retained to remove duplicates.

### Multi-omics Integration Experiments

To demonstrate the effectiveness of `ECLARE` for multi-omic integration of paired and unpaired data, we performed experiments to assess i) whether training a pair of RNA and ATAC encoders using contrastive learning on paired data similar to CLIP outperforms other baseline methods in terms of zero-shot alignment of unseen data, ii) whether integrating KD improves upon the basic zero-shot approach, iii) whether KD further benefits from a multi-teacher approach compared to a single-teacher one, and iv) whether these observations are applicable to unpaired multi-omic data.

#### Evaluation metrics

We assess the performance of `ECLARE` and other integration methods using metrics that capture nuclei pairing accuracy, degree of multimodal integration and conservation of biological information. We provide an overview of these metrics here, although the interested reader can find more details in Luecken et al. ^95^ and Cao and Gao ^12^. All metrics are computed based on embedded features, and most metrics are computed using the scib-metrics library ^95^.

#### FOSCTTM

The fraction of samples closest to the true match (FOSCTTM) quantifies for each cell from one modality (e.g. RNA cell *r*_*i*_) the fraction of non-matching cells from the other modality (*a*_*j*_ ∀ *j*≠ *i*) that are closest to the matching cell (*a*_*i*_). This process is repeated for all cells from both modalities and is followed by global averaging of individual FOSCTTM scores. Because FOSCTTM requires ground-truth information on the true matches (or pairings) across modalities, it can only be evaluated on paired datasets. Note that we report 1-FOSCTTM, such that higher 1-FOSCTTM represents better pairing accuracy.

#### iLISI

The integration local inverse Simpson’s Index (iLISI) metric quantifies the amount of mixing occurring within local cell neighborhoods with regards to a fixed label associated with each cell. For example, the most common label for iLISI is the batch label such that high iLISI values correspond to strong mixing between cells from different batches, a good indication of batch effect removal. In the current study, we use the modality of origin as labels such that high iLISI corresponds to strong mixing between cells from different modalities, a good indication of multimodal integration. We use the KNN variant of iLISI using 30 neighbors, and opt for scaled iLISI to bound its value between 0 and 1.

#### ARI, NMI & ASW

ARI and NMI are reference-based metrics that compare learned cluster labels to reference cell-type labels, differing in how similarity is computed: ARI assesses pairwise label consistency after adjusting for chance, while NMI measures mutual information between label distributions, normalized by entropy. ASW, unlike ARI and NMI, evaluates cluster compactness using only the data-driven labels, without reference labels. All three metrics rely on Leiden clustering with 30 neighbors.

#### Baseline methods

The base model that serves as the building block of our `ECLARE` framework is a lightweight variant of scCLIP^16^. scCLIP trains ATAC and RNA encoders in a contrastive manner by maximizing similarity between a nucleus’s RNA and ATAC embeddings, but it relies on artificially matched atlas-level data based on shared cell-type annotations. By contrast, we use genuinely paired multi-omic data and opt for simpler MLP encoders to maintain true cell-matching fidelity. We simply refer to this MLP-base model as **CLIP**. We also implemented an ablated version of `ECLARE` where the multi-teacher setup is replaced by a single-teacher one to perform KD. We refer to this single-teacher variant as **KD-CLIP**. Other baseline methods sourced from other studies are listed and described below.

**MOJITOO**^31^ trains and filters canonical correlation analysis (CCA) components based on paired data having undergone PCA (RNA) or LSI (ATAC) dimensionality reduction. MOJITOO is not a diagonal integration method by default and does not inherently require gene activity scores; however, for our diagonal integration benchmarking setup, we used gene activity-derived ATAC features to construct a shared gene-level space for cross-modality comparison. **MultiVI** ^11^ trains a variational autoencoder model for each modality while also promoting alignment between variational distributions for joint modeling of counts data. **GLUE** ^12^ also performs multimodal autoencoding while guiding the data reconstruction to mimic the feature-by-feature associations (e.g. genes-by-peaks) encoded within a knowledge graph. **scDART**^13^ performs diagonal integration with an MLP that directly maps an ATAC input layer onto an RNA input layer and forces manifold alignment between the embedded representations arising from the feature sets of both modalities. **scJoint**^14^ is a two-step method where an integrated embeddings space is first used to transfer cell-type labels from RNA cells onto ATAC cells, followed by a refinement step to consolidate the ATAC embeddings afforded by the transferred labels.

All competing baseline methods were provided with the same preprocessed input data (i.e., the same feature sets and normalization where applicable) to ensure comparable input conditions. However, unlike `ECLARE` which underwent hyperparameter tuning via Optuna, the baseline methods were run using their default hyperparameters as recommended by their respective authors. This difference in tuning strategy should be noted when interpreting the benchmarking results.

#### Benchmarking study

For our benchmarking study, we compare single-cell multimodal integration methods on their ability to perform zero-shot alignment on unseen data. The motivation here is to see whether the base CLIP model behind `ECLARE` outperforms other published multi-omics integration methods. In this way, any improvements beyond this base model would showcase the effectiveness of multi-teacher KD even for models with strong transferability. Note that we only used the four primary paired datasets for this benchmarking study (Table 2).

We first compare CLIP with the baseline methods MOJITOO, MultiVI, GLUE, scDART and scJoint using the evaluation metrics on the 4 paired human brain datasets described in Table 2. Each dataset is used as either a source dataset for training or a target dataset for evaluation. Thus, the task consists of performing all 12 unique combinations of source-to-target evaluations. Moreover, each experiment from each of the 12 source-to-target combinations is repeated 3 times by resampling the source and target datasets, similar to cross-validation folds. Hence, a total of 36 experiments are performed for each method. For each experiment, the evaluation metrics are applied onto RNA and ATAC nuclei that have been embedded by the method in question.

We then make similar comparisons between CLIP, KD-CLIP and `ECLARE` on the same 4 paired datasets. Note that for a given target dataset, `ECLARE` consumes all the other 3 source datasets simultaneously due to its multi-teacher mechanism, thus reducing its total number of experiments from 36 to 12. These experiments serve to compare the performance of single-teacher and multi-teacher KD compared to performing zero-shot alignment with CLIP.

After performing experiments using paired data as target datasets, we repeat experiments with CLIP, KD-CLIP and `ECLARE` by treating the unpaired MDD data as the target dataset and using the 4 paired datasets as source datasets. Here, the number of source-to-target combinations reduces from 12 to 4 for CLIP and KD-CLIP and from 4 to 1 for `ECLARE`. Due to the 3 validation folds, a total of 12 experiments are performed for CLIP and KD-CLIP and 3 experiments for `ECLARE`. Note that we do not report 1-FOSCTTM for these experiments due to the lack of ground-truth nuclei pairing information.

### Gene regulatory network analysis on MDD data

Once `ECLARE` is trained on nuclei from case and control subjects from the MDD dataset, it can be used to project ATAC and RNA nuclei into a common embedding space and derive nuclei pairings by solving the OT problem formalized in Eq. (6) and identifying unique pairs contained within the transport plan. These paired nuclei are then used to study how MDD leads to dysregulations in gene regulation and investigate the pathways impacted by dysregulated genes. We selected the MDD student model that resulted in the highest compound metric based on the experiments from the previous section.

#### Pairing nuclei in ECLARE latent space

To pair unpaired ATAC and RNA nuclei using `ECLARE`, nuclei were sampled at the sex and cell-type level and projected through the trained dual encoders. Based on these `ECLARE` embeddings, a two-step process based on OT was used to first ensure that an equal number of nuclei is retained from both modalities and then map these nuclei one-to-one across modalities. OT is performed a first time to identify and remove the excess nuclei from the modality with such excess nuclei. The excess nuclei to be removed are identified based on the OT transport plan, where nuclei from the excess modality with the lowest transport weights to the other modality are removed. Then, OT is performed a second time — now with equal numbers of nuclei from both modalities — to identify the optimal pairings across modalities. The OT cost matrices were computed based on diffusion distances ^96^ in Euclidean form:

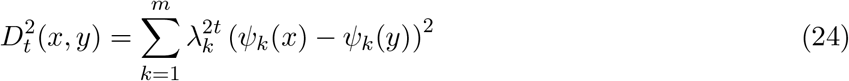

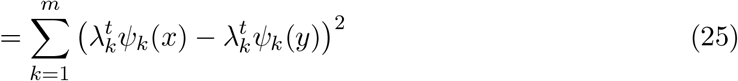

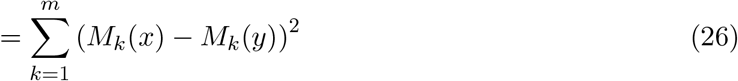

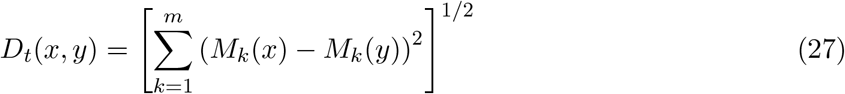

where *ψ*_*k*_(.) represents the projection onto the *k* -th diffusion component and 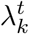 represents the singular value of *ψ*_*k*_ (i.e. *obsm[“X diffmap”]* and *uns[“diffmap evals”]* in Scanpy, respectively). The diffusion coefficient *t* was set to *t* = 1.

Based on the one-to-one nuclei pairings across modalities, the rows from one of the modalities (here, snATAC-seq) are sorted with respect to these pairings such that snATAC-seq and snRNA-seq now have an equal number of nuclei with rows corresponding to paired nuclei.

#### Pseudo-bulking with SEACells

Many of the standard analyses for studying gene regulation based on single-cell ATAC-RNA data involve correlating chromatin accessibility features with gene expression ^18;85;97^. However, correlating highly sparse and noisy single-cell data leads to highly spurious results when studying gene-gene interactions ^98^ and is likely to be exacerbated by cross-modal correlations, especially because scATAC-seq data are even sparser than scRNA-seq data. According to Squair et al. ^98^, one of the best remedies to counter spurious correlations is to aggregate cells into pseudobulks. This aggregation mitigates spurious associations driven by technical noise and sparsity, and helps to better account for biological variability across cells and samples ^98^. However, pseudo-bulking usually refers to aggregating all cells from the same cell type, which would prevent generating multiple cell aggregates for the same cell type. We thus opted to aggregate cells using the SEACells framework ^99^, which uses diffusion modeling to find cells with similar molecular profiles while mitigating the bias of sampling from high-density areas. Importantly, Persad et al. ^99^ have shown that SEACells leads to more reliable peak-gene correlations than directly using the single-cell profiles, making it more reliable for downstream analyses ^100^.

We learn nucleus-to-SEACell assignments by running SEACells on the RNA nuclei, then applying the same assignments to the ATAC nuclei (recall that RNA and ATAC nuclei have been paired). The distances and kernel affinities are computed based on PCA components of the RNA data. The number of SEACells (*n*) formed by aggregating *N* nuclei was set to *n* = max(*N/*100, 15), meaning that each SEACell is associated with 100 nuclei on average, for a minimum of 15 SEACells. During aggregation, gene expression features are averaged across nuclei to normalize by the number of nuclei associated with each SEACell.

#### Differential regulation analysis with sc-compReg

To perform differential gene network analysis, we follow the sc-compReg procedure ^37^. This procedure involves the computation of a transcription factor regulatory potential (TFRP), which quantifies how TF expression and the accessibility of regulatory elements (REs - e.g. ATAC peaks) affects the expression of target genes (TGs). For TF *i*, TG *j*, SEACell pseudobulk *t* and RE *j*, TFRP is computed as follows:

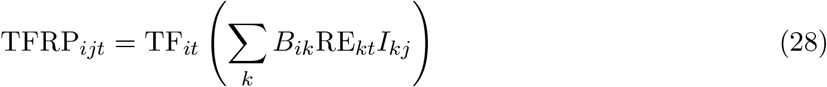

where *B*_*ik*_ corresponds to the binding strength of TF *i* on RE *k* and *I*_*kj*_ corresponds to the interaction strength between RE *k* and TG *j*. We obtain *B*_*ik*_ by computing the log-odds scores between TFs from JASPAR 2020 & 2024 databases and sequences within REs defined by ATAC peaks with the position-specific scoring matrix using pyJASPAR^101^ and PyRanges^102^. *I*_*kj*_ was obtained using the following equation:

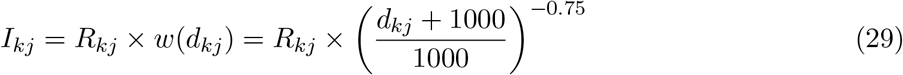

where *R*_*kj*_ is the Spearman correlation coefficient between accessibility values in RE *k* and TG *j*, and *d*_*kj*_ is the linear genomic distance between RE *k* and TG *j*. As shown in Eq. 29, this distance is weighted by a distance power decay function *w*(.), which is the formula used by the *dist power decay* function in ‘scglue’ library^12^ that we employed.

Although TFRP is calculated for each TF-TG pair (i.e. TFRP_*ijt*_) and directly used for downstream analyses in the original sc-compReg paper ^37^, we derive a single TFRP value per TG by aggregating TFRP_*ijt*_ values (Eq. 28) across TFs, i.e.

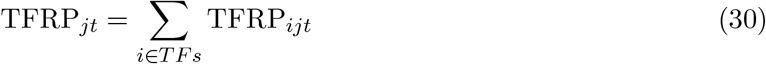

Having a single TFRP value per TG for each SEACell enables more interpretable gene enrichment analyses, as explained later on.

In sc-compReg, the TFRP is used as an estimator of the expression of its associated target gene TG. Specifically, TFRP and TG are related by a linear regression model as follows:

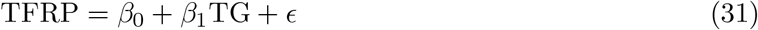

where *β*_0_ and *β*_1_ are standard regression bias and coefficient parameters, and *ϵ* ∼*N* (0, *σ*^2^). By fitting these parameters, estimates of TFRP (denoted 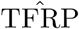) can be computed based on TG and used to compute conditional Gaussian maximum likelihoods (*β*_0_ and *β*_1_ grouped into *β*):

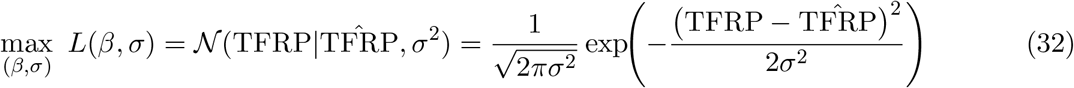

Likelihoods derived from linear regression are computed in this manner on condition-specific nuclei (i.e. MDD cases and controls) as well as nuclei pooled across both conditions, giving rise to *L*_*case*_(*β*_*case*_, *σ*_*case*_), *L*_*ctrl*_(*β*_*ctrl*_, *σ*_*ctrl*_) and *L*_*pooled*_(*β*_*pooled*_, *σ*_*pooled*_). From these condition-specific and condition-agnostic likelihood values, a likelihood ratio (LR) is computed to quantify the degree to which MDD alters the association between TFs & REs and TGs (the ‘max’ are dropped for clarity):

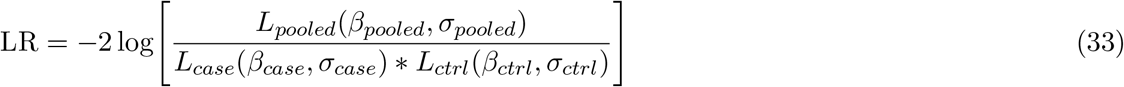

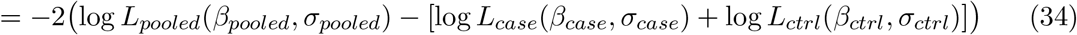

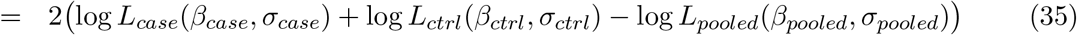

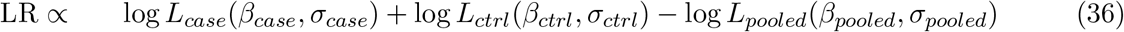

The computed LR is then tested for statistical significance based on an LR null model. For establishing a null distribution of LR values, Duren et al. ^37^ demonstrate that the standard chi-square distribution (with three degrees of freedom) is not a suitable null model and is thus replaced by fitting a gamma distribution to the computed LR values corresponding to the lower 50th percentile of the empirical distribution of LR values. We thus adopt this procedure to derive a null model from a fitted gamma distribution and use it to determine TGs whose LR exceeds the 95th percentile. This filtering procedure thus produces a gene set containing TGs that are differentially regulated in MDD cases versus controls.

Note that the statistical modeling and filtering of LR values is performed on sex-specific and cell type-specific sets of nuclei. In this way, we can study the sex and cell-type determinants of expression dysregulation caused by MDD. Moreover, we limit the number of TF-RE-TG combinations by focusing on GRNs outlined in the Brain Single-Cell Omics for PsychENCODE (brainSCOPE) resource provided by the PsychENCODE Consortium^103^. Leveraging this resource avoids analyzing the excessive number of unique TF-RE-TG combinations (*n* ≈ 10^12^) to focus on cortical gene regulatory programs (*n* ≈ 1.5 × 10^5^), thus also limiting the number of spurious comparisons. Although brainSCOPE provides cell type-specific TF-RE-TG combinations, we pool combinations across all cell types to generate a single set of candidate GRNs.

#### Enrichment Analysis of filtered gene sets

We use the gene set derived from the sc-compReg procedure to perform gene enrichment analyses. We perform EnrichR analysis using GSEApy ^104;105^ based on pathways defined in *Brain*.*GMT*, a curated database of brain-related functional gene sets ^38^. We further trimmed out functional gene sets derived from non-cortical areas (notably, hippocampus and nucleus accumbens) as was done in a meta-analysis of early life stress effects on prefrontal cortex in MDD ^106^. This enrichment analysis is performed for each sex & cell-type combination, resulting in 14 sets of enrichment results of filtered *Brain*.*GMT* pathways.

Based on the analyses described just above, we identified a *Brain*.*GMT* pathway of interest labeled aston_major_depressive_disorder_dn for which enrichment was significant across many sex & cell-type combinations. For each sex & cell-type combination, we extracted the genes from this pathway that overlapped with those derived from the sc-compReg procedure and aggregated these genes via union to form a single gene set. Note that only sex & cell-type combinations with significant enrichment for aston major depressive disorder dn were considered for this aggregation. Thus, this aggregated gene set represents genes that are differentially regulated in MDD in at least one of the eligible combinations. We repeated EnrichR with this aggregated gene set to identify pathways whose enrichment directly depends on genes contributing to enrichment of aston major depressive disorder dn across different sex and cell types. This second enrichment step, conceptually related to post-enrichment refinement strategies such as Leading Edge Metagene (LEM) analysis ^41^ and integrative approaches like ActivePathways ^42^, serves to distill biologically coherent pathways from genes repeatedly implicated across contexts, providing a more focused view of the mechanisms shared across sex and cell types.

#### Pathway-level differential expression

The gene enrichment analyses described in the previous section yield enriched pathways whose genes overlap with genes enriched in an MDD-specific pathway. We investigated differential gene expression at the pathway level by first using *scanpy*.*tl*.*score genes* to generate pathway expression scores for each SEACell pseudobulk. Recall that these pseudobulks were formed for each sex & cell-type combination for both case and control conditions separately. Hence, given an enriched pathway and sex & cell-type combination, a pathway score is computed for all pseudobulks from both conditions. We then performed a weighted t-test between pseudobulk pathway scores across conditions, where each pseudobulk is weighted by the number of nuclei mapped onto it. The weighted t-tests were performed using *DescrStatsW, get compare* and *ttest ind* from the statsmodels python library^107^.

#### Pruning GRNs by testing TF-TG pairs for differential regulation

To enable further downstream analyses, we pruned the brainSCOPE GRNs to identify TFs mediating differential regulation. For a given sex & cell-type combination, brainSCOPE GRNs were restricted to GRNs for which the TG overlapped with the genes identified based on the sc-compReg procedure. In this way, candidate TF-TG GRN pairs are first anchored on genes already identified as mediating differential regulation. Then, TF-TG pairs are further pruned by running the standard sc-compReg analysis ^37^, which computes LR scores for each TF-TG pair (Eq. 28) as opposed to performing our proposed gene-aggregated variant that yields LR scores at the TG level only (Eq. 30). Computing LR scores for each TF-TG pair allows the construction of a null distribution (via a fitted gamma distribution) and statistical filtering to prune the GRNs by retaining TF-TG pairs whose LR exceeds the 95th percentile. This filtering process is performed for all sex & cell-type combinations, yielding different sets of filtered TF-TG pairs.

After identifying the *NR4A2* -*ABHD17B* regulatory edge in excitatory neurons from female donors (Results), we extracted all TFs linked with pruned TF-TG edges involving *ABHD17B* associated with this same group of cells. We then also extracted all direct TGs of those newly-extracted TFs. This GRN – which includes *ABHD17B*, the TF regulators of *ABHD17B* as well as their other direct targets – was further reduced in size based on the k-core algorithm ^108^ (*k* = 5). Note that *NR4A2* and *NR3C1* were white-listed to ensure that they remain in the downsized GRN. We then clustered the TF-TG edges using sklearn’s *SpectralCoclustering* function on the binary TF-TG adjacency matrix with *k* = 3 clusters. *ABHD17B* was removed from the clustering operation due to its ubiquitous association with all TFs, and *NR4A2* was also removed in light of only having an edge with *ABHD17B*. Graph processing and visualization were performed using NetworkX ^109^ and HoloViews ^110^ libraries.

In summary, we filtered brainSCOPE GRNs by first anchoring TF-TG pairs on TGs already identified as being differentially regulated. We then extracted TF-TG pairs shared across all sex & cell-type combinations and further pruned TF-TG pairs based on regulon enrichment. Recall that brainSCOPE GRNs are composed of TF-RE-TG triplets, such that a given filtered TF-TG pair also includes its associated REs.

### Integration of unpaired developmental data

We performed `ECLARE` integration of the **CTX Velmeshev** dataset using **PFC Zhu** and **PFC V1 Wang** paired datasets to train teacher models. After aligning the RNA and ATAC features between source and target datasets, the teacher and student models were trained using nuclei from all cell types, but we restrict nuclei to those belonging to the inhibitory and excitatory lineages for downstream analysis. This subset of nuclei is then used to perform pseudotime inference and SCENIC+ eRegulon cell scoring, as explained below.

#### PAGA and diffusion pseudotime

To infer pseudotime values in an unsupervised manner, we performed Leiden clustering followed by partition-based graph abstraction (PAGA ^66^) to obtain a coarse denoised graph based on similarities between Leiden clusters. Except for *n neighbors* = 15, we used default parameters for the *pca, neighbors, leiden* and *paga* functions in Scanpy. The PAGA graph was then used to initialize the force-directed layout used for further visualization, which uses the ForceAtlas2 wrapper (FA). Following PAGA, we performed diffusion pseudotime (DPT^67^) based on a root nucleus defined as the nucleus closest to the centroid of 2nd trimester nuclei, as annotated by Velmeshev et al. ^36^. Other parameters for DPT were set to their default values. All steps related to PAGA and DPT were executed with Scanpy^111^.

#### Supervised pseudotime inference via ordinal regression

As demonstrated by Macnair et al. ^112^, pseudotime inference can be achieved in a supervised manner using ordinal regression modeling. We thus trained a MLP-based ordinal regression model using the CORAL framework^68^ with the **CTX_Velmeshev** dataset. Briefly, CORAL minimizes a cross-entropy loss of *K* -1 binary classifiers, where *K* corresponds to the number of unique ordinal labels (the “age range” variable in the case of **CTX_Velmeshev**, Fig. 4a). Each binary classifier estimates the probability that the true ordinal label *y*_*i*_ of a given sample exceeds the classifier’s associated rank threshold *r*_*k*_ according to

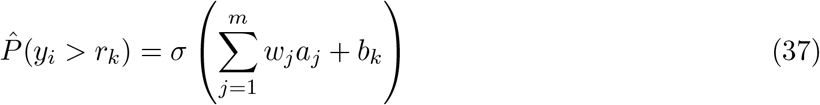

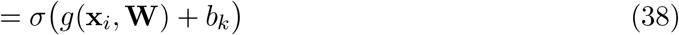

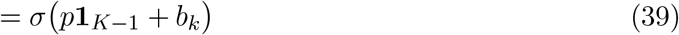

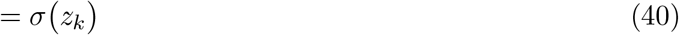

where *σ*(·) denotes the sigmoid function, *a*_*j*_ are the activations of the penultimate layer, 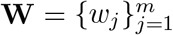 are shared weights, *b*_*k*_ is the bias term specific to the *k*-th threshold, *p* is the “raw ordinal pseudotime” value (more on this later), **1**_*K*−1_ is a vector-of-ones used to broadcast *p* into *K*-1 values and *z*_*k*_ is the classifier logit for the *k*-th threshold. Crucially, rank consistency is imposed between successive classifier outputs by enforcing weight sharing across all *K-1* classifiers, such that **W** is shared while *b*_1_ *> b*_2_ *>* · · · *> b*_*K*−1_. The combination of shared weights and monotonically decreasing bias terms ensures that the classifier logits *z*_*k*_ are themselves monotonically decreasing, thereby enforcing ordinal consistency across thresholds.

One separate CORAL model is trained for ATAC and RNA nuclei from the target **CTX_Velmeshev** dataset. ATAC and RNA mini-batches are sampled separately, leading to ATAC- and RNA-specific CORAL losses. For simplicity, these two modality-specific losses are averaged together into a single total loss that is minimized to optimize both CORAL models simultaneously. The architectures consist of an MLP with a fully connected hidden layer of dimension 256, followed by a CORAL layer consisting of another fully connected layer mapping the 256 hidden units to one hidden variable *p* (Eqs 37-39). This single latent variable—which we interpret as the raw ordinal pseudotime value inferred by CORAL—is then expanded into logits for the *K* -1 classification thresholds via the bias terms *b*_*k*_ that obey the rank-consistency constraint (Eq. 40). Here, *K* = 8 in light of the eight developmental stages annotated in the **CTX_Velmeshev** dataset, listed in the last sub-panel of Fig. 4a. CORAL models were trained in PyTorch to minimize classification cross-entropy (i.e. the custom *coral loss* implemented by the authors of CORAL) for 100 epochs using AdamW optimizer with a learning rate of 0.001 and weight decay of 0.01.

The raw ordinal pseudotime values generated for each nucleus by CORAL’s fully-connected layer were further processed after training to obtain the ordinal pseudotime values used in the results reported in Fig. 4a,c. Specifically, the raw values were normalized by standard deviation across all nuclei then transformed by the sigmoid function to constrain values between 0 and 1. These refinements were performed separately on RNA and ATAC nuclei.

#### Computing evaluation metrics on integrated CTX Velmeshev data

In addition to computing the previously-described iLISI and NMI metrics on the modality, cell type and age range variables, we also included the cell-type LISI (cLISI) score ^95^ on the lineage variable. The cLISI metric quantifies the degree of cell-type (un)mixing in local cell neighborhoods, where higher cLISI scores reflect cell neighborhoods that are more pure in terms of cell-type annotation. For the lineage variable, cLISI was favoured over NMI since there are only two lineages retained for analysis, i.e. excitatory neurons (ExNeu) and inhibitory interneurons (IN). Note that metrics were computed on a separate set of Leiden clusters created specifically for metric evaluation, where the resolution parameter was set to 0.25. In addition to these metrics, we also computed the Spearman correlation between unsupervised diffusion pseudotime (DPT) and supervised ordinal pseudotime & age range. All metrics and correlation values were computed on each method’s latent embedding of **CTX_Velmeshev** nuclei (`ECLARE`, scJoint and GLUE; Fig. S7). Error bars were obtained via bootstrapping using 500 resampled iterations, where nuclei were resampled with replacement and used to compute metrics and correlation values at each iteration.

#### SCENIC+ cell-scoring of VIP eRegulons

SCENIC+ is a pipeline that performs inference of GRNs based on single-cell ATAC-seq and RNA-seq data ^113^. Key building blocks for these GRNs are eRegulons, which center around a TF and the genes that it targets alongside the enhancer elements that the TF interacts with.

We extracted the eRegulons included in Supplementary Data 3 from Velmeshev et al. ^36^, which contains pre-computed gene regulatory network information including TFs, their TGs with associated importance weights, and target chromatin regions with region-to-gene linkage scores. To process these data with the SCENIC+ framework, we first converted our ATAC-seq data into a cisTopic object using the pycisTopic library^113;114^, performing Latent Dirichlet Allocation (LDA) topic modeling with 20 topics over 100 iterations to identify co-accessible chromatin regions. We then created a SCENIC+ object by integrating the RNA expression data (stored as an AnnData object), the cisTopic chromatin accessibility object, and the parsed eRegulon definitions, reconstructing the transcription factor regulatory modules with their original importance weights and correlation values.

Using the SCENIC+ cell-scoring functionality, we applied the AUCell algorithm to compute activity scores for each eRegulon in individual cells, generating both gene-based scores from RNA expression enrichment of TGs and region-based scores from ATAC accessibility enrichment of target regions. We specifically examined the *DLX5* and *POU2F1* eRegulons that were identified as VIP interneuron-specific regulatory programs. To visualize cell-type-specific regulatory activity, we plotted the normalized eRegulon scores on force-directed graph embeddings and generated density plots to highlight VIP neuron spatial distribution, enabling direct comparison of regulatory program activity across cell types and developmental stages.

### Co-embedding of developmental and MDD data

#### Training of KD-CLIP MDD student model

We trained an MDD student model based on single-teacher knowledge distillation using **PFC_Zhu** as the paired source dataset (Fig. 5a.i-ii). Both teacher and student models were trained exclusively on excitatory neurons. Model architecture and training strategy remained largely unchanged from what is described in the Methods section ‘CLIP, KD-CLIP and `ECLARE` model training’, with the exception of the latent embedding size that was reduced from 256 to 64 hidden units. This contraction in latent dimensions provided more compact and structured integration of **PFC_Zhu** and MDD data within the latent space. After training, nuclei from MDD dataset excitatory neurons were sampled by balancing uniformly for age and condition, then projected alongside **PFC_Zhu** nuclei into a common embedding using the student model (Fig. 5a.iii). For age balancing, we used binned donor ages based on the pandas *qcut* method (q=17 bins) ^115;116^. This binning provided an optimal trade-off between total number of nuclei, representation across donor ages and balancing of ordinal pseudotime values across case & control nuclei.

#### Cross-dataset label imputation, characterization of Leiden clusters and inferring ordinal pseudotime

From co-embedded RNA and ATAC nuclei, we built an adjacency graph spanning both datasets using Scanpy’s *neighbors* function and then performed kNN classification (k=10) to impute developmental cell-type labels onto the MDD data (Fig. 5a.iv). Specifically, for each nucleus from MDD data, its nearest-neighbors in the **PFC_Zhu** data were extracted to identify the most frequent cell-type label and then assigned to the queried nucleus from MDD data. The optimal parameters used for building this graph consisted of 8 input PCA components (explains 66% of variance) and *n neighbors*=50, with other parameters assuming their default values.

The cross-dataset graph was also used to infer Leiden clusters (*resolution*=0.6) and build a PAGA graph spanning both datasets using Scanpy. Also, visualizing the graph and its Leiden clusters exposes two major branches originating from a common root cluster. Hence, we assigned labels to these two branches based on the dominant sub-cell-type based on MDD RNA data annotations: one branch is dominated by layer 2/3 neurons (branch L23) whereas the other branch is dominated by layer 4/5/6 neurons (branch L46). Once labels were assigned to these two branches, Leiden clusters were labeled by their position along these branches, stemming from a common root cluster (cluster 0) and evolving throughout its respective branch, i.e. clusters 1 L23 to 5 L23 for branch L23 and 1 L46 to 5 46 for branch L46. Labels for other clusters neither falling into one of the two branches nor representing the root cluster were left unchanged and were not used in subsequent analysis (i.e. cluster 2).

We also trained RNA and ATAC CORAL models using the **PFC_Zhu** dataset to impute ordinal pseudotime values onto co-embedded nuclei. We used the same CORAL model architecture and training procedures as previously, with the exception of the conversion of raw ordinal pseudotime values into processed values. Instead of normalizing raw values by standard deviation and transforming with the sigmoid function, we performed histogram matching between raw values and age bins using *match histograms* from the skimage library^117^. This results in a distribution of ordinal pseudotime values that matches the age bins distribution, therefore generating values that are more akin to years-since-birth and thus more interpretable in the context of MDD.

#### Statistically testing differences in ordinal pseudotime

We statistically compared ordinal pseudotime values between MDD cases and controls along the clusters distributed across the L23 and L46 branches, with separate tests for male and female donors.

To quantify global differences for a given branch of interest, we first computed cluster-specific means and standard errors for both case and control nuclei for clusters belonging to that branch. These cluster-level summaries are then used to compute the difference in mean pseudotime between conditions for each cluster along the branch, along with the standard error of that difference. A weighted average of these cluster-level differences was computed via an intercept-only weighted least-squares using *WLS* class from statsmodels ^107^, where the weights were derived from cluster-level standard errors. The resulting mean and standard error were used to derive a Z-statistic from which a single-tail *p*-value was computed according to the hypothesis that mean pseudotime for cases is greater than that for controls. Note that this procedure is equivalent to a fixed-effect model.

In addition to testing branch-level differences in ordinal pseudotime values using this fixed-effect method, we also tested for cluster-specific differences using Welch’s t-test to highlight the clusters within each branch showing the highest case vs control differences.

#### Pseudotemporal clustering of gene sets

We followed the procedures from Zhu et al. ^63^ to infer clusters of genes based on their pseudo-temporal pattern of expression. We first pre-filtered genes based on the Moran’s I statistic that identifies genes whose expression is associated with pseudotime with statistical significance ^118^. Moran’s I was calculated for each gene using the PySAL/ESDA python library^119^ by providing processed pseudo-counts values and a cell connectivity graph, with 1000 random permutations to infer p-values for each gene. We derived a connectivity graph that directly reflects ordinal pseudotime by executing the *sc*.*pp*.*neighbors* function to nuclei embedded in CORAL’s latent space, using the CORAL model previously trained with **PFC_Zhu**’s RNA dataset. The nominal p-values were then subjected to multiple comparison correction using Benjamini-Hochberg, with the resulting q-values used for selecting genes at the *q <* 0.01 level of significance.

Once genes were pre-filtered using Moran’s I, gene expression profiles were smoothed along the ordinal pseudotime axis using the pyGAM library ^120^ to fit a Generalized Additive Model (GAM). We used a gamma distribution with log links and 9 knots to fit cubic splines through the ordered expression profiles for each gene separately. Then, the smoothed expression profiles underwent k-means clustering using scikit-learn ^121^ to cluster genes into four clusters labeled km1 to km4.

#### Detecting differentially-accessibly TF motifs with pychromVAR

The pychromVAR library is a python implementation of chromVAR^77^ and is part of the scverse ecosystem ^122^. As input to pychromVAR, we pseudobulked snATAC-seq counts data by imputed developmental cell type and by condition. We then used pychromVAR’s functions to add the peak’s sequences based on hg38 reference genome, add GC-bias information, add background peaks, fetch JASPAR2020 motifs (collection=‘CORE’, tax group=[‘vertebrates’]) and perform motif matching. Then, we computed accessibility deviations using a custom version of pychromVAR’s *compute deviations* method that only differs by the deviation scores the function outputs. Specifically, we ensured that the observed deviations (*Y*) as well as mean and standard deviation of background deviations — *µ*(*Y* ^′^) and *σ*(*Y* ^′^), respectively — were all returned, rather than only returning the Z-scored deviations, i.e. 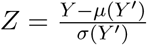. Having access to these quantities allowed us to compute the bias-corrected deviation *Y* − *µ*(*Y* ^′^), which is the quantity used by the authors of chromVAR^77^ to compute differential accessibility. As such, a bias-corrected deviation value was computed for each pseudobulk and each TF motif. Differential accessibility was then computed using a Welch’s t-test for unequal variances based on bias-corrected deviations for cases and controls. FDR correction was then applied onto the Welch nominal p-values based on the Benjamini-Hochberg procedure.

## Code & data availability

Code is made available at github.com/li-lab-mcgill/ECLARE. A derived data bundle containing the harmonized inputs consumed by the analysis pipelines is deposited on Zenodo (doi.org/10.5281/zenodo.14794845). All primary data analyzed in this study are publicly available through the accession numbers reported in Table 2; no new primary datasets were generated for this study. Per-figure reproduction recipes are documented in the repository’s “Reproducing manuscript figures” section and are pinned to the immutable git tags `v1.0-main` (Figs. 1, 3, 5), v1.0-fig2-benchmark and v1.0-fig2-og (Fig. 2), and `v1.0-fig4-dev` (Fig. 4); each tagged snapshot includes a top-level `REPRODUCE.md` with the exact commands. The repository is released under the MIT License.

## Acknowledgments

Y.L. is supported by Canada Research Chair (Tier 2) in Machine Learning for Genomics and Healthcare (CRC-2021-00547), Natural Sciences and Engineering Research Council (NSERC) Discovery Grant (RGPIN-2016-05174) and Canadian Institutes of Health Research (CIHR) Project Grant (PJT-540722). D.M.K. is supported by NSERC Canada Graduate Research Scholarship Doctoral (CGS D) scholarship. We acknowledge contributions from Doruk Ç akmakçi for helpful discussions pertaining to methods, results and biological interpretation.

## Author contributions

D.M.K. contributed to study design, analysis, writing and revision. Y.L. contributed to study design, writing and revision. A.C. contributed towards writing and revision. G.T. and C.N. contributed towards revision.

## Supplementary Materials

## Supplementary Methods

### Genomic region enrichment analysis using rGREAT

To functionally interpret sets of regulatory regions identified across sex and cell-type combinations, we performed genomic region enrichment analysis using the rGREAT package ^123^, an R interface to the Genomic Regions Enrichment of Annotations Tool (GREAT^39^). This analysis tests whether genomic intervals are non-randomly associated with curated gene sets by assigning regulatory domains to genes and evaluating enrichment based on genomic proximity rather than direct gene overlap.

Custom gene sets were provided using a cortical brain–specific GMT file from the *Brain*.*GMT* resource, with gene symbols mapped to Entrez Gene identifiers to ensure compatibility with GREAT. For each gene set, symbol-to-Entrez mapping was performed using the *org*.*Hs*.*eg*.*db* annotation database, retaining only uniquely mapped identifiers. The resulting list of Entrez-based gene sets was supplied to GREAT as user-defined annotations.

For each sex and each major cell type (astrocytes, endothelial cells, excitatory neurons, inhibitory neurons, microglia, oligodendrocyte precursor cells, and oligodendrocytes), we analyzed a corresponding BED file containing genomic coordinates of regulatory peaks of interest. BED files were converted into GRanges objects and analyzed independently to preserve cell-type– and sex-specific regulatory context.

Enrichment analysis was performed using the human genome build hg38. GREAT was applied to each set of genomic regions using default regulatory domain definitions, and enrichment statistics were computed for all supplied gene sets.

### Gene set enrichment using H-MAGMA

To further evaluate whether genes exhibiting differential regulatory activity are enriched for genetic risk associated with MDD, we performed gene set enrichment analysis using H-MAGMA^40^ (Hi-C–coupled Multi-marker Analysis of GenoMic Annotation), an extension of MAGMA that incorporates chromatin interaction profiles derived from Hi-C data to improve the assignment of noncoding genetic variants to their putative target genes. This analysis tests whether gene sets identified from our multi-omic regulatory framework are non-randomly associated with gene-level association statistics derived from genome-wide association studies (GWAS), thereby linking regulatory dysregulation to inherited genetic risk.

Genes prioritized by the sc-compReg–based differential regulatory analysis were used as input gene sets for H-MAGMA. Specifically, genes exhibiting significant likelihood ratio statistics—reflecting altered associations between chromatin accessibility, transcription factor activity, and gene expression across conditions—were aggregated into sex- and cell-type–specific gene sets and tested for enrichment against MDD GWAS–derived gene-level association statistics.

H-MAGMA analyses were conducted using precomputed GWAS resources accessed through the Psychiatric Genomics Consortium (https://doi.org/10.7488/ds/2458) based on the study of Howard et al. ^124^. Output statistics from H-MAGMA, including regression coefficients and corresponding significance values, were extracted and summarized separately for each sex and cell-type combination.

## Supplementary Figures

**Figure S1:**
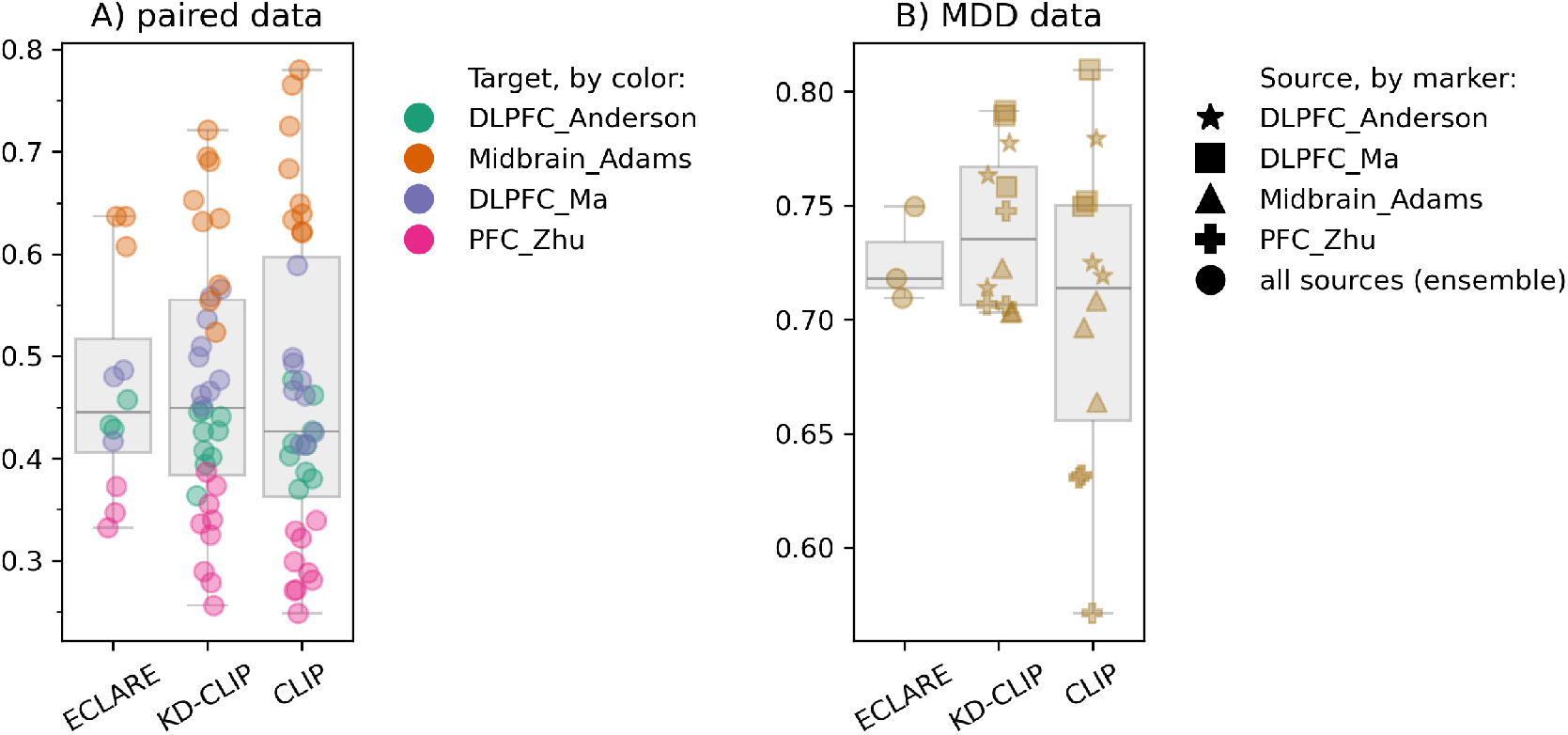
Batch correction performance assessed with acceptance rates of k-nearest neighbor batch effect test (kBET) for (**a**) paired data and (**b**) MDD data. The higher the kBET acceptance rate the better. On paired data, data points color-coded by the target dataset evaluated for each experiment. On unpaired MDD data, data points shape-coded by the source dataset(s) used to train individual models. For `ECLARE`, we refer to the source dataset as “all sources (ensemble)” since all source datasets are used.

**Figure S2:**
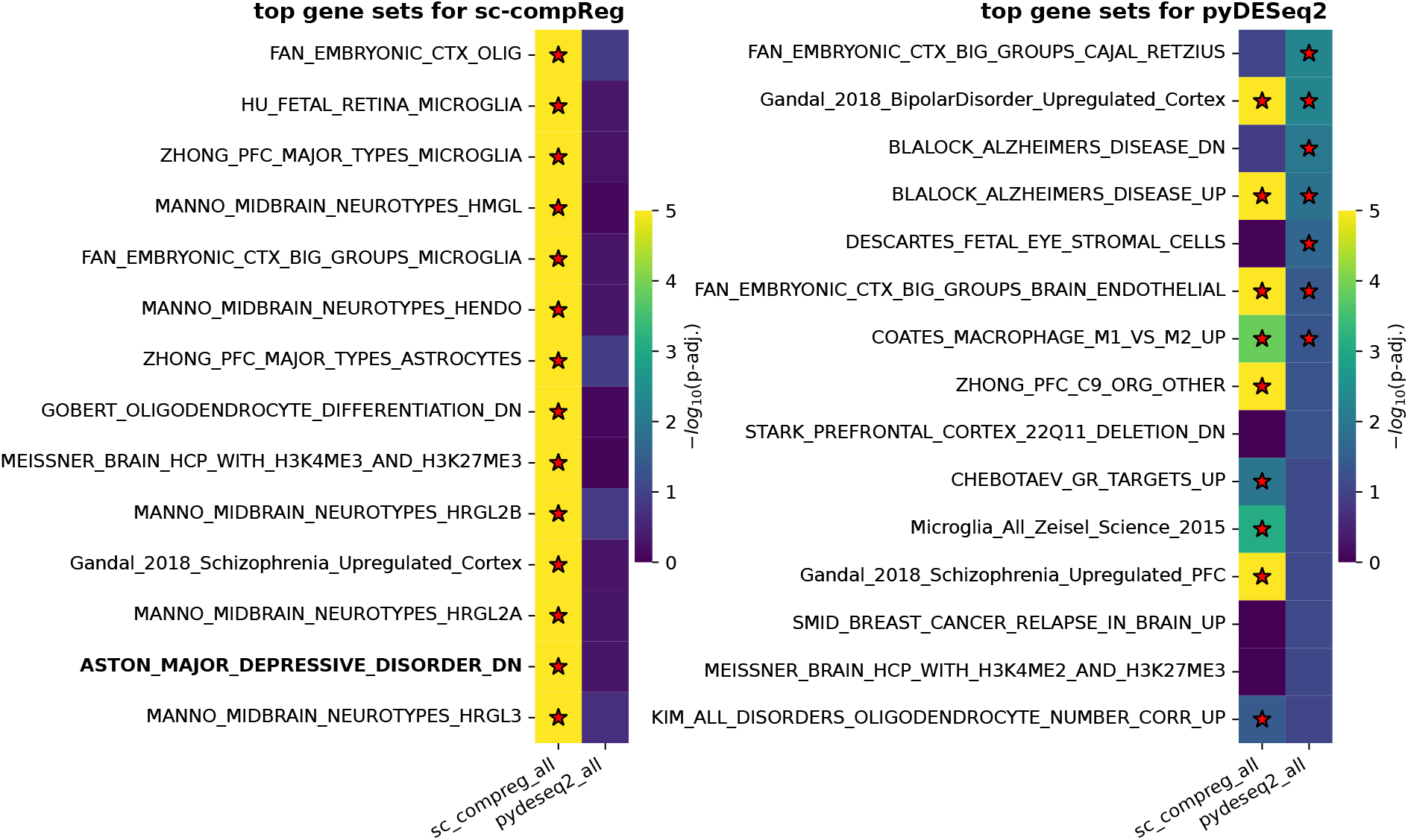
Enrichment results for Brain.GMT gene-sets. The top-15 enriched gene sets when using sc-compReg (left) or pyDESeq2 (right). In both cases, results are compared across both methods. Red stars denote gene sets that are significantly enriched according to EnrichR. Label for aston major depressive disorder dn in bold-face for ease of reference.

**Figure S3:**
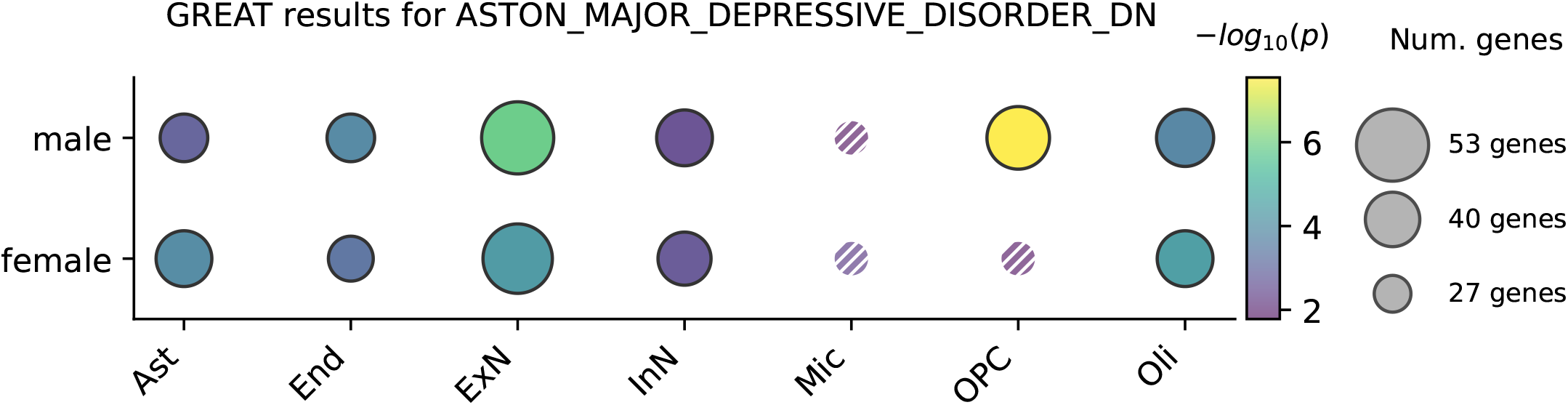
Results from Genomic Regions Enrichment of Annotations Tool (GREAT) enrichment analysis. Solid circles denote significantly-expressed gene sets after Bonferroni correction, hashed circles denote nominally significant gene sets failing to pass Bonferroni correction, and small gray crosses denote gene sets failing to attain nominal significance.

**Figure S4:**
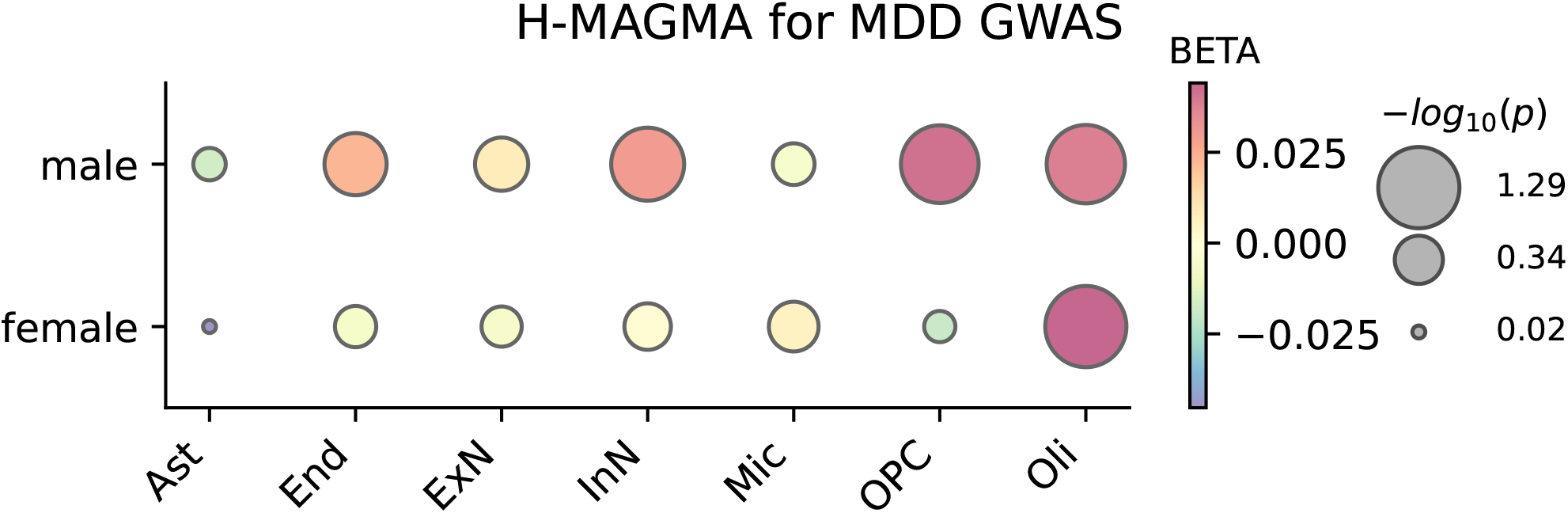
H-MAGMA results for MDD GWAS. Because all results fall below statistical FDR-corrected significance, size-coded enrichment levels are shown for all sex & cell-type combinations to enable comparison of relative differences in enrichment. Results are color-coded by the BETA regression fit parameter returned by the H-MAGMA analysis.

**Figure S5:**
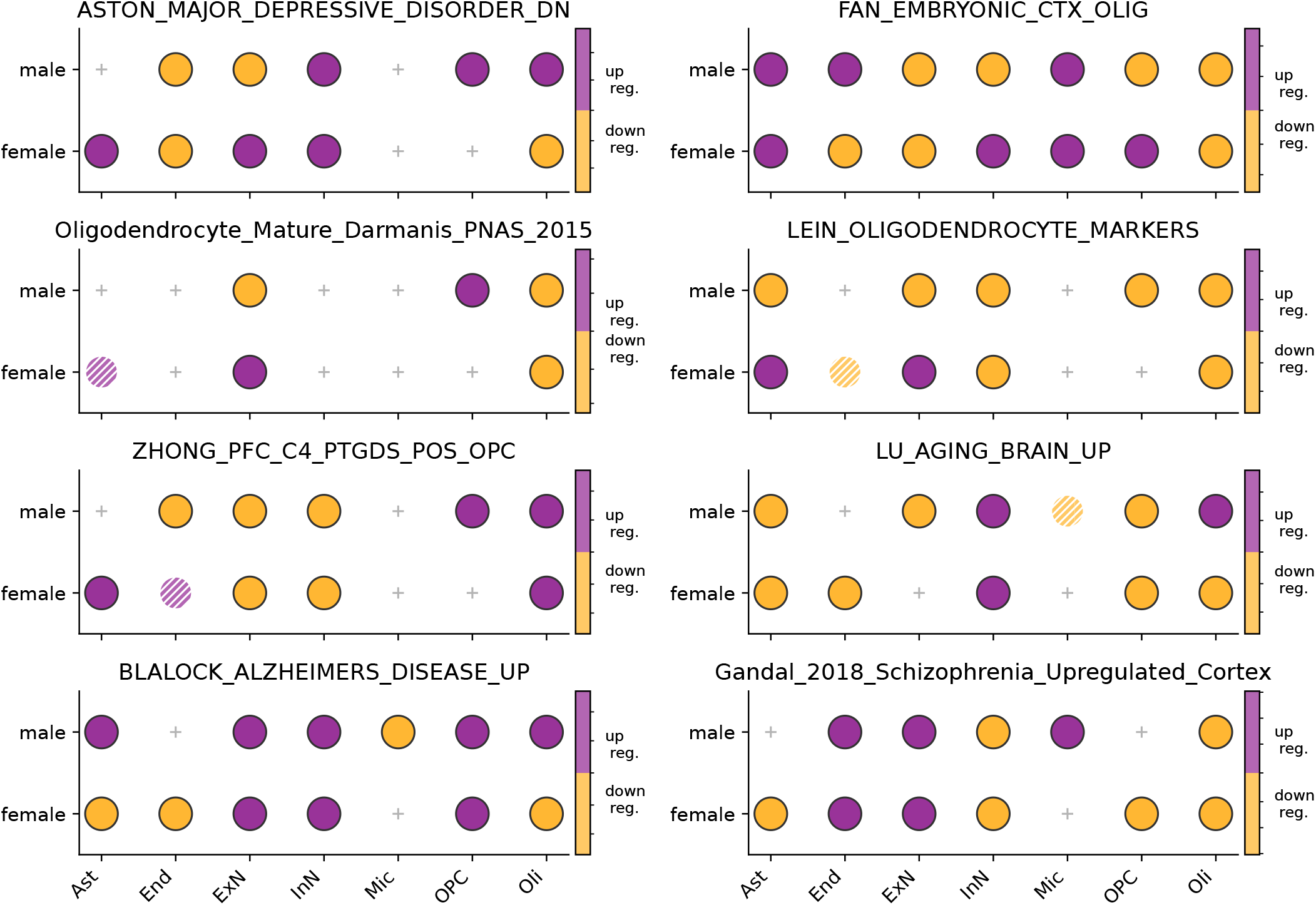
Differential expression of selected Brain.GMT gene sets at the gene set-level using module scores, following the MDD leading-edge analysis. Solid circles denote significantly-expressed gene sets after Bonferroni correction, hashed circles denote nominally significant gene sets failing to pass Bonferroni correction, and small gray crosses denote gene sets failing to attain nominal significance.

**Figure S6:**
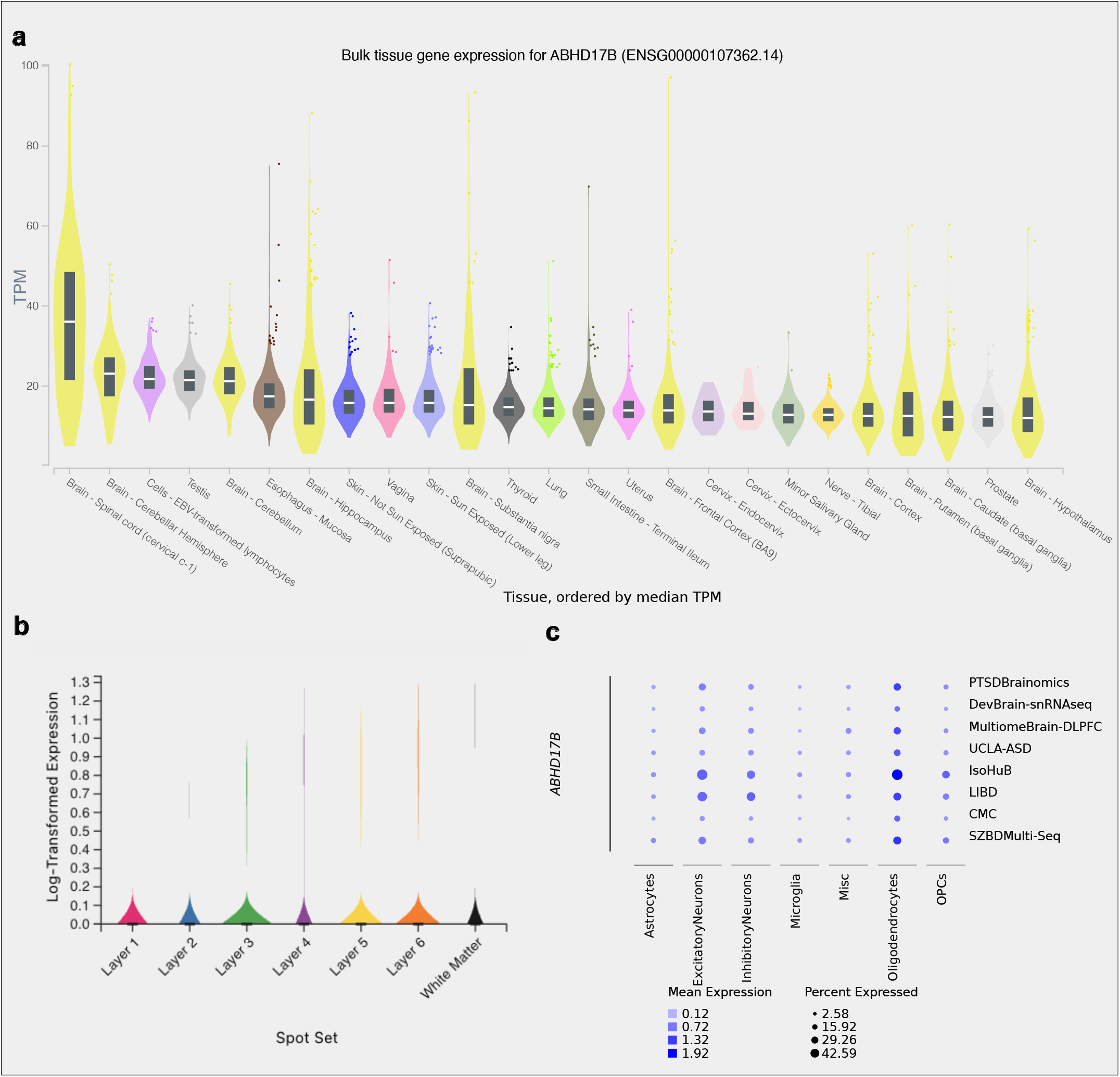
Evidence that *ABHD17B* is expressed in the brain. **(a**) Bulk tissue expression for *ABHD17B* for different tissues, ranked by median transcript-per-million (TPM). Violin plots for brain & spinal cord tissues are in yellow. Plot generated by GTEx online browser. **(b)**. Log-transformed expression of *ABHD17B* across cortical layers and white matter for *Brain 6522 - Anterior* sample in psychSCREEN online browser. **(c)**. Multi-dataset summary of *ABHD17B* expression across major cell types, taken from psychSCREEN online browser.

**Figure S7:**
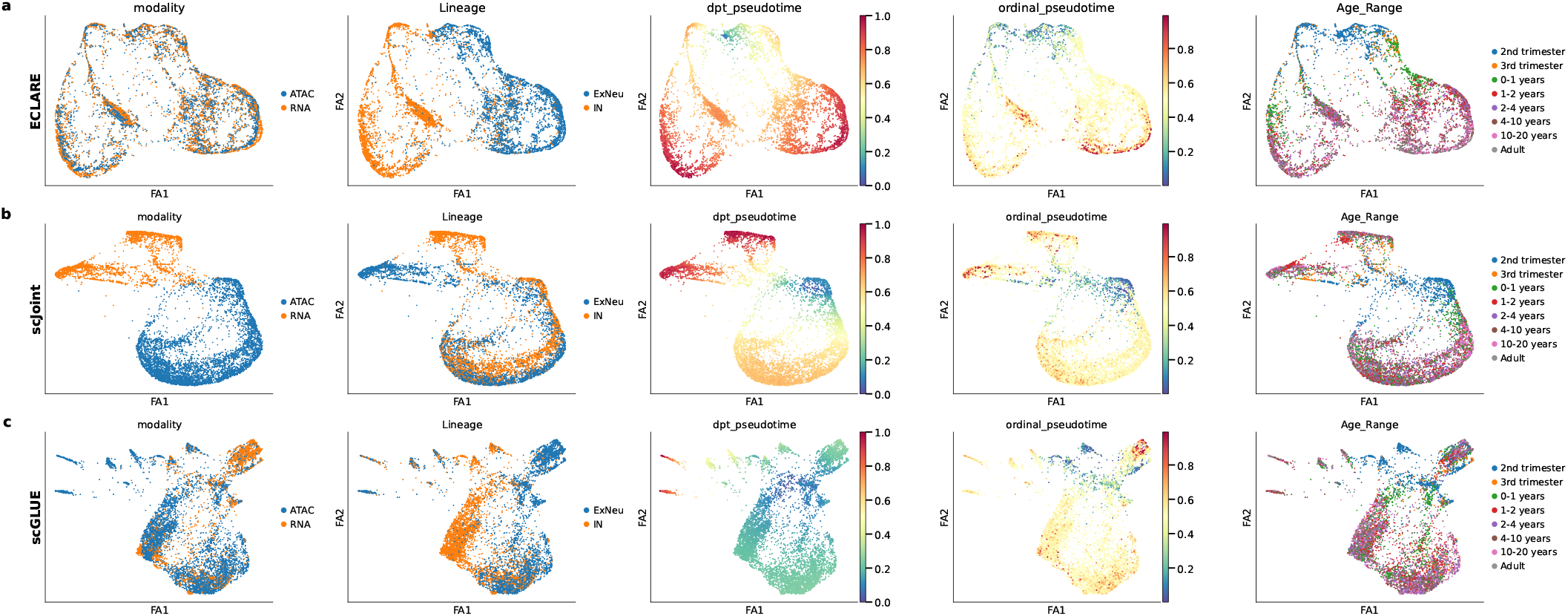
Embedded nuclei from excitatory neurons and inhibitory interneurons for **(a)** `ECLARE`, **(b)** scJoint and **(c)** GLUE. ‘dpt_pseudotime’ refers to diffusion pseudotime.

**Figure S8:**
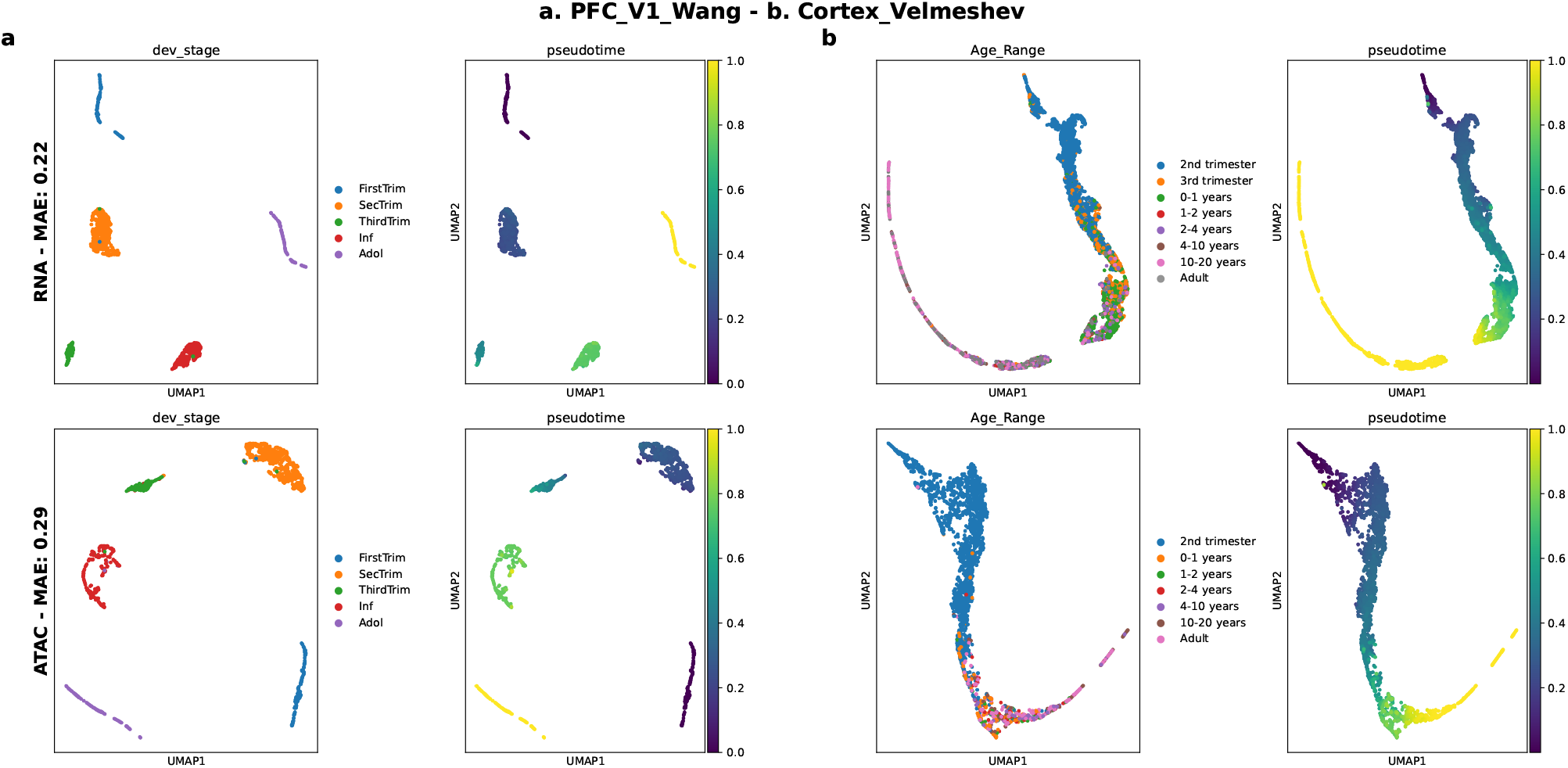
UMAP of nuclei embedded in CORAL latent space for snRNA-seq (top row) and snATAC-seq (bottom row). **(a)** Embedded neurons from source dataset on which CORAL model was trained on (**PFC_V1_Wang**) colored by developmental stage and inferred ordinal pseudotime. The mean absolute error (MAE) based on held-out source nuclei is reported in the y-axis titles. **(b)** Embedded neurons from target dataset (**CTX_Velmeshev**) colored by age range and inferred ordinal pseudotime.

**Figure S9:**
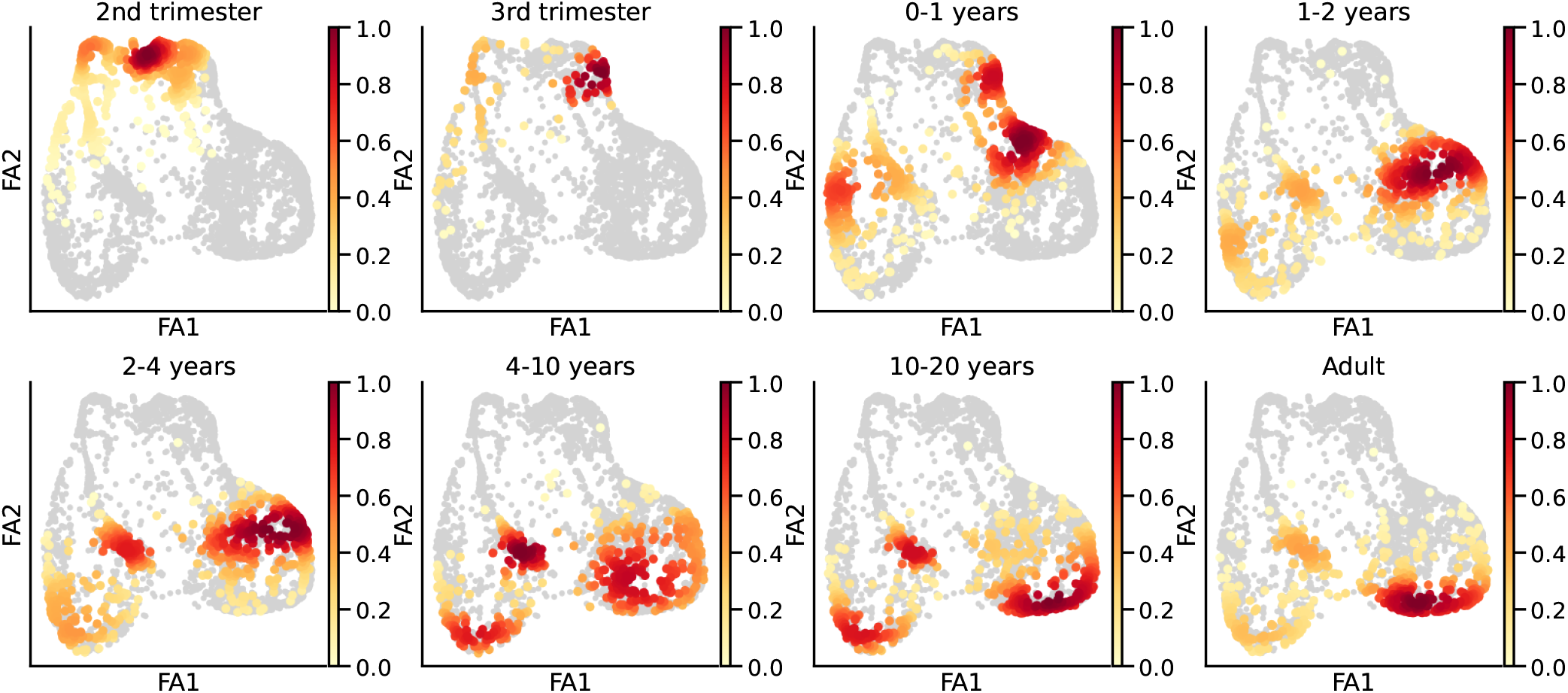
Density plot of **CTX_Velmeshev** data for each age range in an `ECLARE` embedding, plotted with force-directed layouts (FA).

**Figure S10:**
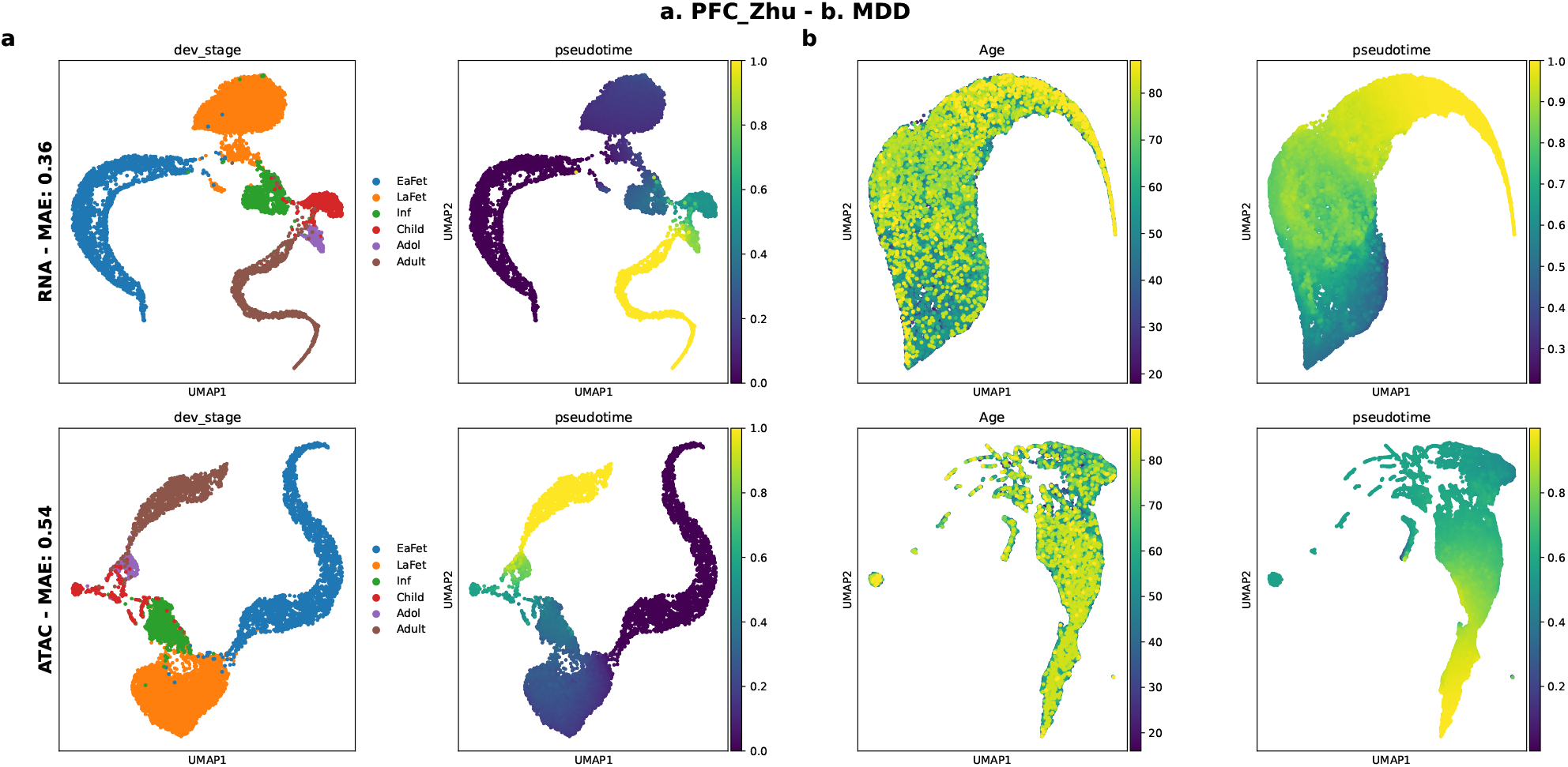
UMAP of excitatory neurons in CORAL latent space for snRNA-seq (top row) and snATAC-seq (bottom row). **(a)** Embedded neurons from source dataset on which CORAL model was trained on (**PFC_Zhu**) colored by developmental stage and inferred ordinal pseudotime. The mean absolute error (MAE) based on held-out source nuclei is reported in the y-axis titles. **(b)** Embedded neurons from target dataset (**MDD**) colored by donor age and inferred ordinal pseudotime.

**Figure S11:**
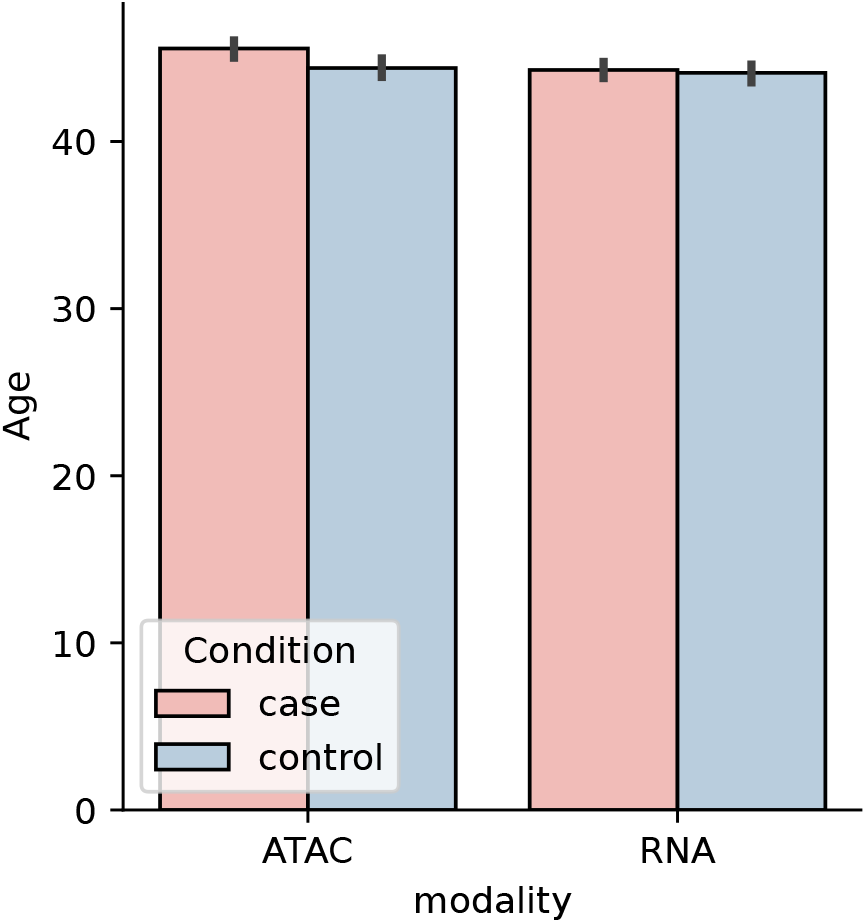
Balancing MDD nuclei for donor age. Mean donor age for each MDD condition label (case & controls) and modality (ATAC & RNA).

**Figure S12:**
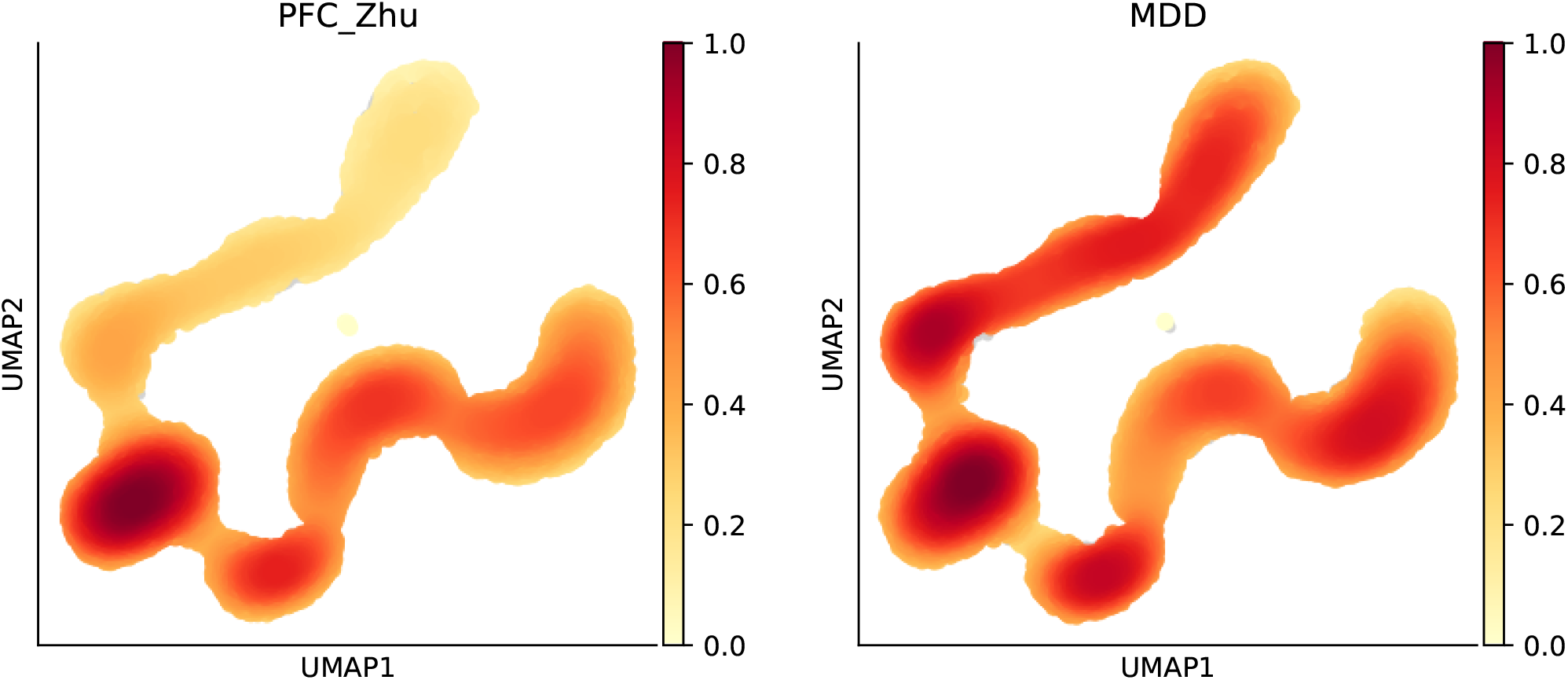
Density of co-embedded nuclei in ECLARE embedding. Nuclei separated by dataset, i.e. developmental **PFC_Zhu** data (left) and MDD data (right). Nuclei density obtained with *sc*.*tl*.*embedding density* and plotted with *sc*.*pl*.*embedding density* from Scanpy.

**Figure S13:**
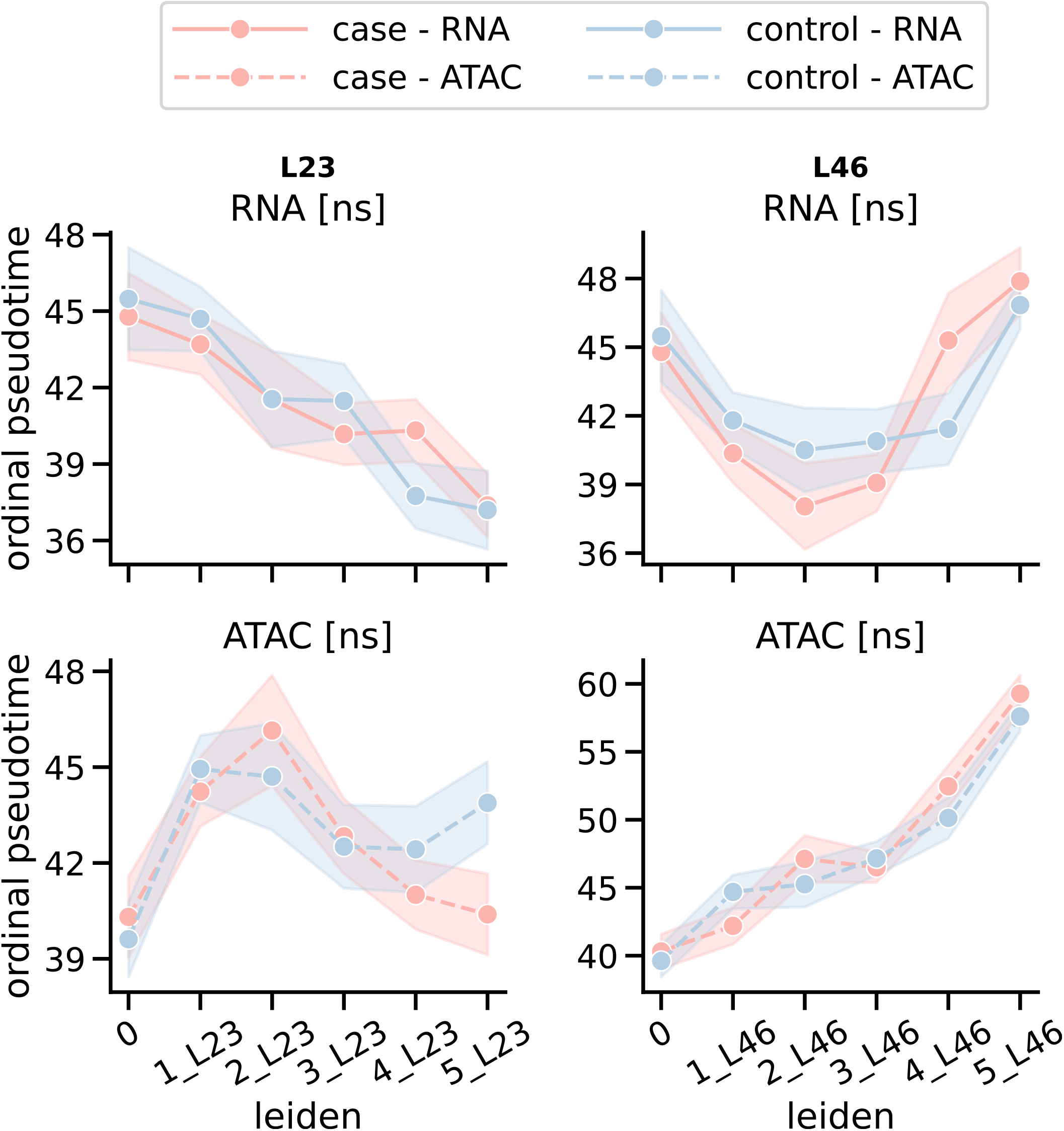
Male-specific analysis of case vs control differences in ordinal pseudotime per Leiden cluster organized along the two longitudinal branches. Results are organized by modalities (rows) and longitudinal branches (columns), where branch-level significance is annotated in sub-panel title and cluster-level significance is annotated above data points. ns: non-significant.

**Figure S14:**
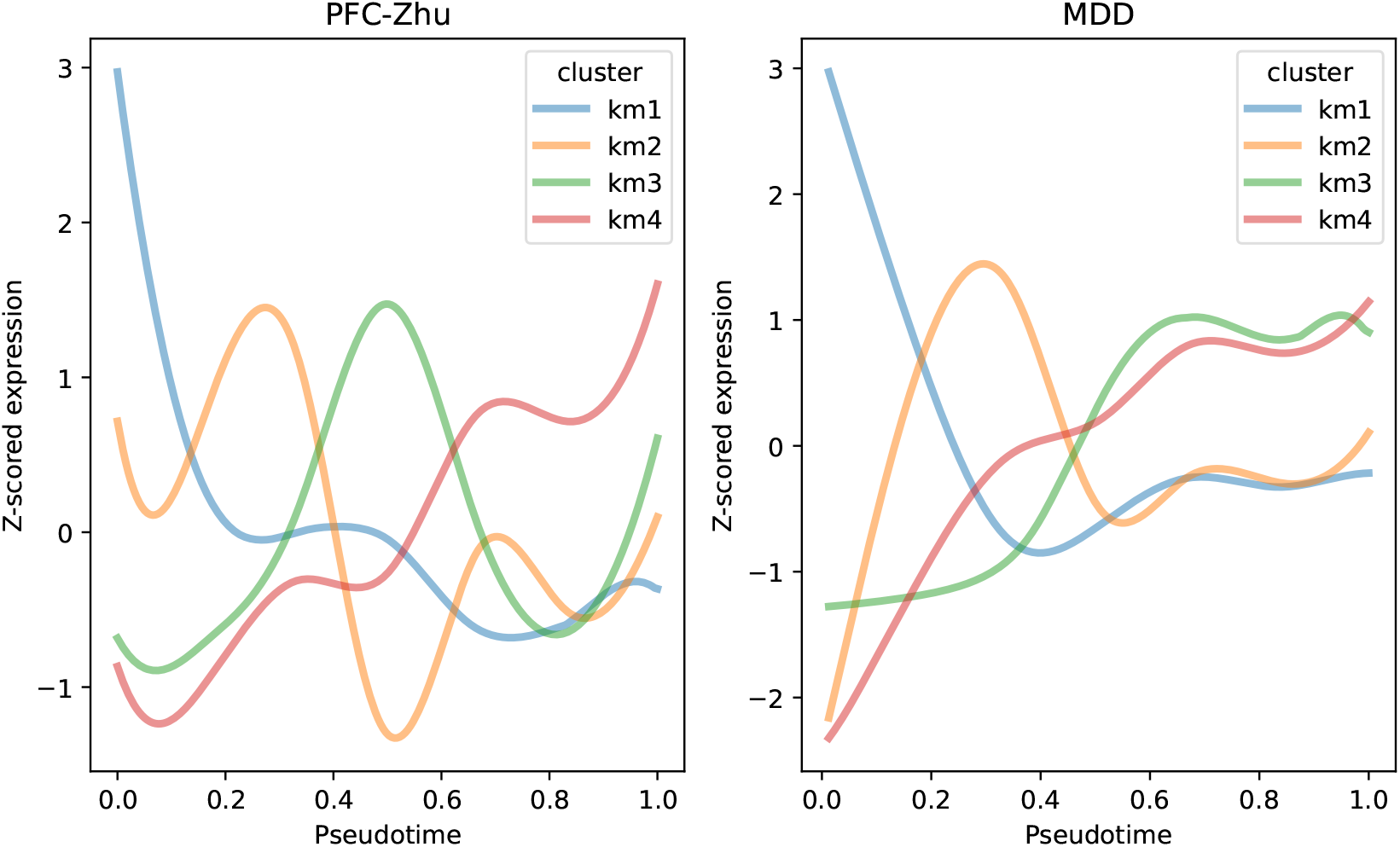
Mean smoother trajectories for pseudotemporal gene clusters (i.e. “km cluster”) derived separately from **PFC_Zhu** (left) and MDD (right) datasets. The x-axis represents scaled ordinal pseudotimes and y-axis represents z-scored gene expression.

**Figure S15:**
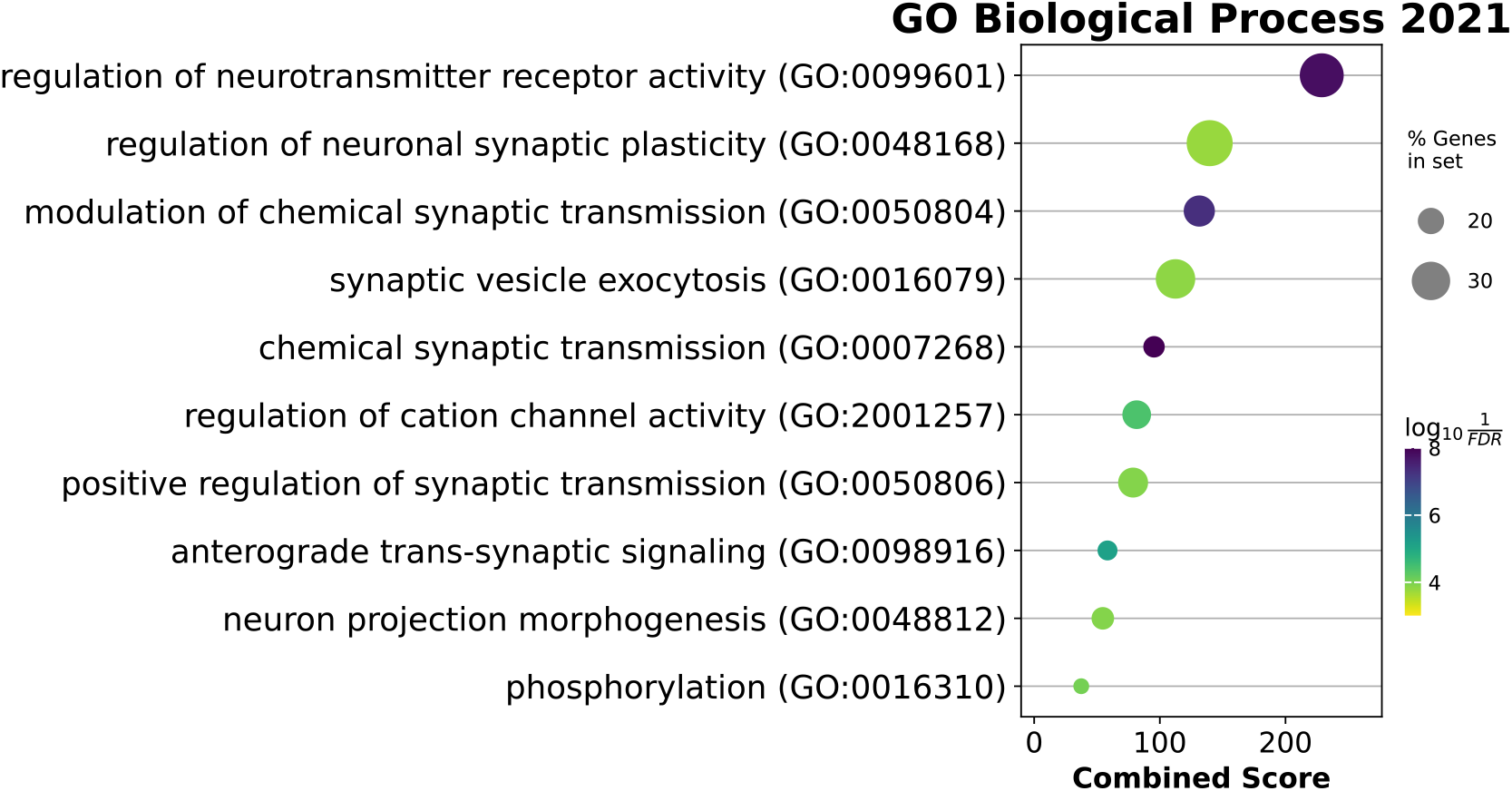
Enrichment results for genes differentially-expressed in MDD that overlap with **km3_mdd** and *EGR1* regulon. Enrichment analysis was performed with EnrichR based on the “GO Biological Process 2021” gene sets.

**Figure S16:**
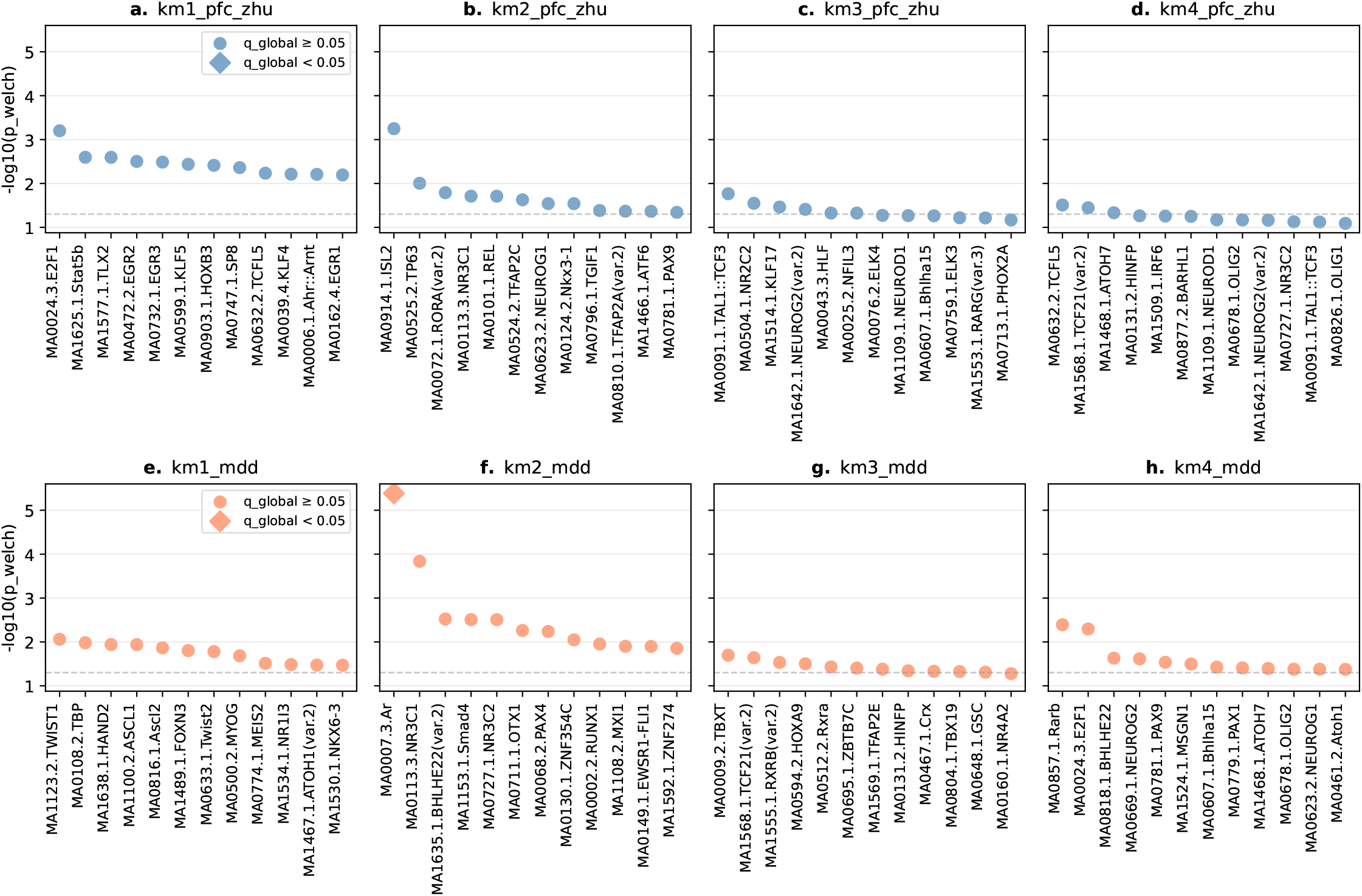
pychromVAR results for differential accessibility in MDD, by pseudotemporal cluster. A diamond-shaped marker denotes significant differential accessibility after global FDR correction, whereas a circle-shaped marker denotes non-significance after FDR correction. A dotted horizontal reference line is placed at the nominal significance level of *p* = 0.05 (i.e. −log_10_(p) ≈ 1.3). Top row shows results for pseudotemporal cluster from **PFC_Zhu** whereas bottom row shows results for clusters from MDD.

**Figure S17:**
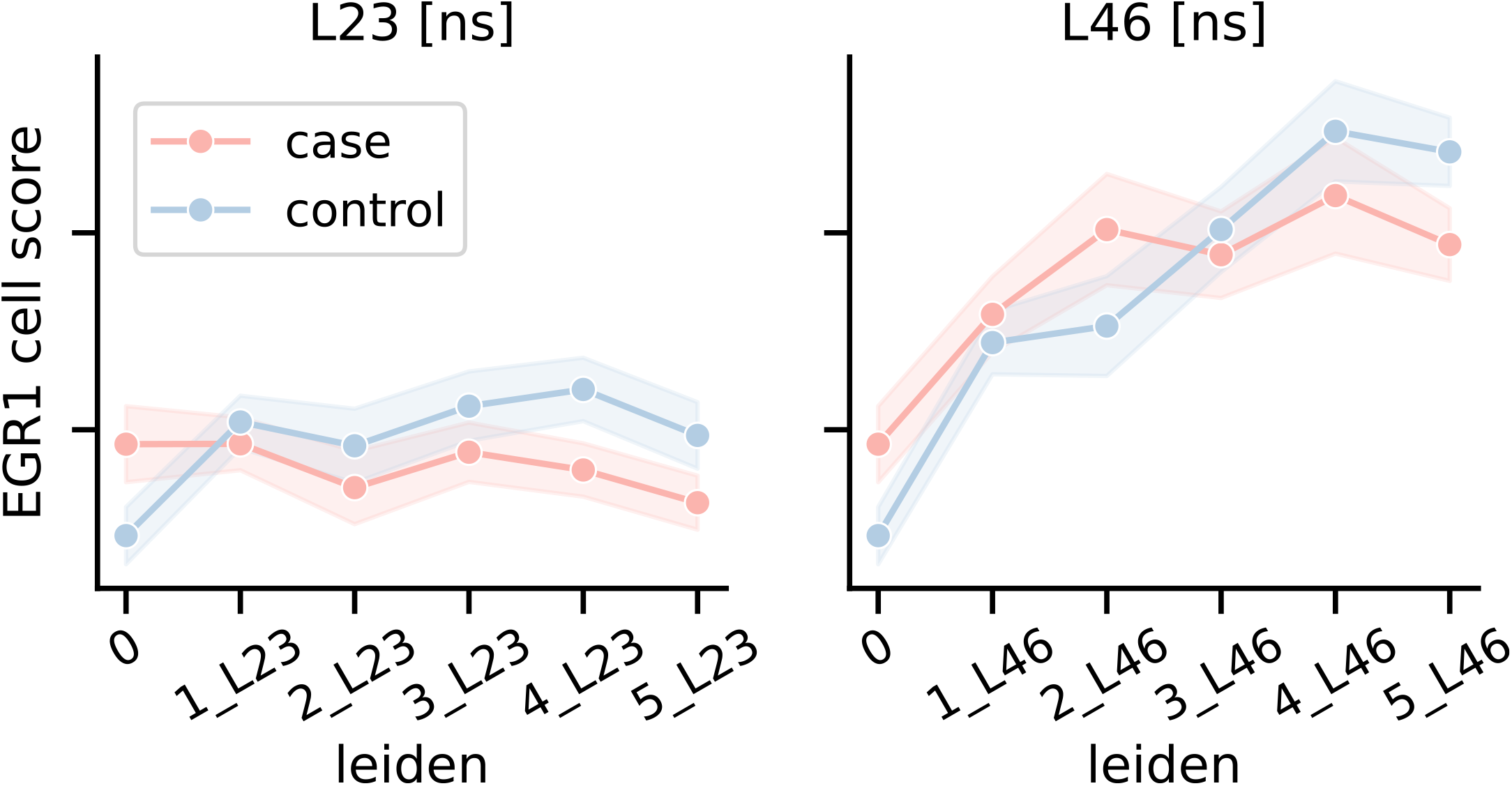
Case and control multi-omic cell regulatory scores for *EGR1* eRegulons across Leiden branches, restricted to nuclei from male donors. Branch-level significance is annotated in sub-panel title and cluster-level significance is annotated above data points. ns: non-significant.

## References

[1] Ricard Argelaguet, Anna S. E. Cuomo, Oliver Stegle, and John C. Marioni. Computational principles and challenges in single-cell data integration. Nature Biotechnology, 39(10):1202–1215, October 2021. ISSN 1087-0156, 1546-1696. doi: 10.1038/s41587-021-00895-7. URL https://www.nature.com/articles/s41587-021-00895-7.

[2] Song Chen, Blue B. Lake, and Kun Zhang. High-throughput sequencing of the transcriptome and chromatin accessibility in the same cell. Nature Biotechnology, 37(12):1452–1457, December 2019. ISSN 1546-1696. doi: 10.1038/s41587-019-0290-0. URL https://www.nature.com/articles/s41587-019-0290-0. Publisher: Nature Publishing Group.

[3] Sai Ma, Bing Zhang, Lindsay M. LaFave, Andrew S. Earl, Zachary Chiang, Yan Hu, Jiarui Ding, Alison Brack, Vinay K. Kartha, Tristan Tay, Travis Law, Caleb Lareau, Ya-Chieh Hsu, Aviv Regev, and Jason D. Buenrostro. Chromatin Potential Identified by Shared Single-Cell Profiling of RNA and Chromatin. Cell, 183(4):1103–1116.e20, November 2020. ISSN 00928674. doi: 10.1016/j.cell.2020.09.056. URL https://linkinghub.elsevier.com/retrieve/pii/S0092867420312538.

[4] Pau Badia-i Mompel, Lorna Wessels, Sophia Müller-Dott, Rémi Trimbour, Ricardo O. Ramirez Flores, Ricard Argelaguet, and Julio Saez-Rodriguez. Gene regulatory network inference in the era of single-cell multi-omics. Nature Reviews Genetics, 24(11): 739–754, November 2023. ISSN 1471-0064. doi: 10.1038/s41576-023-00618-5. URL https://www.nature.com/articles/s41576-023-00618-5. Publisher: Nature Publishing Group.

[5] The Tabula Sapiens Consortium. The Tabula Sapiens: A multiple-organ, single-cell transcriptomic atlas of humans. Science, 376(6594):eabl4896, May 2022. doi: 10.1126/science.abl4896. URL https://www.science.org/doi/10.1126/science.abl4896. Publisher: American Association for the Advancement of Science.

[6] Kai Zhang, James D. Hocker, Michael Miller, Xiaomeng Hou, Joshua Chiou, Olivier B. Poirion, Yunjiang Qiu, Yang E. Li, Kyle J. Gaulton, Allen Wang, Sebastian Preissl, and Bing Ren. A single-cell atlas of chromatin accessibility in the human genome. Cell, 184(24):5985–6001.e19, November 2021. ISSN 0092-8674, 1097-4172. doi: 10.1016/j.cell.2021.10.024. URL https://www.cell.com/cell/abstract/S0092-8674(21)01279-4. Publisher: Elsevier.

[7] Lukas Heumos, Anna C. Schaar, Christopher Lance, Anastasia Litinetskaya, Felix Drost, Luke Zappia, Malte D. Lücken, Daniel C. Strobl, Juan Henao, Fabiola Curion, Herbert B. Schiller, and Fabian J. Theis. Best practices for single-cell analysis across modalities. Nature Reviews Genetics, 24(8):550–572, August 2023. ISSN 1471-0064. doi: 10.1038/s41576-023-00586-w. URL https://www.nature.com/articles/s41576-023-00586-w. Publisher: Nature Publishing Group.

[8] Jill E. Moore, Michael J. Purcaro, Henry E. Pratt, Charles B. Epstein, Noam Shoresh, Jessika Adrian, Trupti Kawli, Carrie A. Davis, Alexander Dobin, Rajinder Kaul, Jessica Halow, Eric L. Van Nostrand, Peter Freese, David U. Gorkin, Yin Shen, Yupeng He, Mark Mackiewicz, Florencia Pauli-Behn, Brian A. Williams, Ali Mortazavi, Cheryl A. Keller, Xiao-Ou Zhang, Shaimae I. Elhajjajy, Jack Huey, Diane E. Dickel, Valentina Snetkova, Xintao Wei, Xiaofeng Wang, Juan Carlos Rivera-Mulia, Joel Rozowsky, Jing Zhang, Surya B. Chhetri, Jialing Zhang, Alec Victorsen, Kevin P. White, Axel Visel, Gene W. Yeo, Christopher B. Burge, Eric Lécuyer, David M. Gilbert, Job Dekker, John Rinn, Eric M. Mendenhall, Joseph R. Ecker, Manolis Kellis, Robert J. Klein, William S. Noble, Anshul Kundaje, Roderic Guigó, Peggy J. Farnham, J. Michael Cherry, Richard M. Myers, Bing Ren, Brenton R. Graveley, Mark B. Gerstein, Len A. Pennacchio, Michael P. Snyder, Bradley E. Bernstein, Barbara Wold, Ross C. Hardison, Thomas R. Gingeras, John A. Stamatoyannopoulos, and Zhiping Weng. Expanded encyclopaedias of DNA elements in the human and mouse genomes. Nature, 583(7818):699–710, July 2020. ISSN 1476-4687. doi: 10.1038/s41586-020-2493-4. URL https://www.nature.com/articles/s41586-020-2493-4. Publisher: Nature Publishing Group.

[9] Mengjie Chen. Capturing cell-type-specific activities of cis-regulatory elements from peak-based single-cell ATAC-seq. Cell Genomics, 5(3):100806, March 2025. ISSN 2666-979X. doi: 10.1016/j.xgen.2025.100806. URL https://www.ncbi.nlm.nih.gov/pmc/articles/PMC11960509/.

[10] Yang Eric Li, Sebastian Preissl, Xiaomeng Hou, Ziyang Zhang, Kai Zhang, Yunjiang Qiu, Olivier B. Poirion, Bin Li, Joshua Chiou, Hanqing Liu, Antonio Pinto-Duarte, Naoki Kubo, Xiaoyu Yang, Rongxin Fang, Xinxin Wang, Jee Yun Han, Jacinta Lucero, Yiming Yan, Michael Miller, Samantha Kuan, David Gorkin, Kyle J. Gaulton, Yin Shen, Michael Nunn, Eran A. Mukamel, M. Margarita Behrens, Joseph R. Ecker, and Bing Ren. An atlas of gene regulatory elements in adult mouse cerebrum. Nature, 598(7879):129–136, October 2021. ISSN 1476-4687. doi: 10.1038/s41586-021-03604-1. URL https://www.nature.com/articles/s41586-021-03604-1. Publisher: Nature Publishing Group.

[11] Tal Ashuach, Mariano I. Gabitto, Rohan V. Koodli, Giuseppe-Antonio Saldi, Michael I. Jordan, and Nir Yosef. MultiVI: deep generative model for the integration of multimodal data. Nature Methods, 20(8):1222–1231, August 2023. ISSN 1548-7105. doi: 10.1038/s41592-023-01909-9. URL https://www.nature.com/articles/s41592-023-01909-9. Publisher: Nature Publishing Group.

[12] Zhi-Jie Cao and Ge Gao. Multi-omics single-cell data integration and regulatory inference with graph-linked embedding. Nature Biotechnology, 40(10):1458–1466, October 2022. ISSN 1546-1696. doi: 10.1038/s41587-022-01284-4. URL https://www.nature.com/articles/s41587-022-01284-4. Publisher: Nature Publishing Group.

[13] Ziqi Zhang, Chengkai Yang, and Xiuwei Zhang. scDART: integrating unmatched scRNA-seq and scATAC-seq data and learning cross-modality relationship simultaneously. Genome Biology, 23(1):139, June 2022. ISSN 1474-760X. doi: 10.1186/s13059-022-02706-x. URL https://genomebiology.biomedcentral.com/articles/10.1186/s13059-022-02706-x.

[14] Yingxin Lin, Tung-Yu Wu, Sheng Wan, Jean Y. H. Yang, Wing H. Wong, and Y. X. Rachel Wang. scJoint integrates atlas-scale single-cell RNA-seq and ATAC-seq data with transfer learning. Nature Biotechnology, 40(5):703–710, May 2022. ISSN 1087-0156, 1546-1696. doi: 10.1038/s41587-021-01161-6. URL https://www.nature.com/articles/s41587-021-01161-6.

[15] Pinar Demetci, Rebecca Santorella, Björn Sandstede, William Stafford Noble, and Ritambhara Singh. SCOT: Single-Cell Multi-Omics Alignment with Optimal Transport. Journal of Computational Biology, 29(1):3–18, January 2022. doi: 10.1089/cmb.2021.0446. URL https://www.liebertpub.com/doi/10.1089/cmb.2021.0446. Publisher: Mary Ann Liebert, Inc., publishers.

[16] Lei Xiong, Tianlong Chen, and Manolis Kellis. scCLIP: Multi-modal Single-cell Contrastive Learning Integration Pre-training. October 2023. URL https://openreview.net/forum?id=KMtM5ZHxct.

[17] Tim Stuart, Andrew Butler, Paul Hoffman, Christoph Hafemeister, Efthymia Papalexi, William M. Mauck, Yuhan Hao, Marlon Stoeckius, Peter Smibert, and Rahul Satija. Comprehensive Integration of Single-Cell Data. Cell, 177(7):1888–1902.e21, June 2019. ISSN 0092-8674. doi: 10.1016/j.cell.2019.05.031. URL https://www.sciencedirect.com/science/article/pii/S0092867419305598.

[18] Jeffrey M. Granja, M. Ryan Corces, Sarah E. Pierce, S. Tansu Bagdatli, Hani Choudhry, Howard Y. Chang, and William J. Greenleaf. ArchR is a scalable software package for integrative single-cell chromatin accessibility analysis. Nature Genetics, 53(3):403–411, March 2021. ISSN 1061-4036, 1546-1718. doi: 10.1038/s41588-021-00790-6. URL https://www.nature.com/articles/s41588-021-00790-6.

[19] Christina V. Theodoris, Ling Xiao, Anant Chopra, Mark D. Chaffin, Zeina R. Al Sayed, Matthew C. Hill, Helene Mantineo, Elizabeth M. Brydon, Zexian Zeng, X. Shirley Liu, and Patrick T. Ellinor. Transfer learning enables predictions in network biology. Nature, 618(7965):616–624, June 2023. ISSN 1476-4687. doi: 10.1038/s41586-023-06139-9. URL https://www.nature.com/articles/s41586-023-06139-9. Publisher: Nature Publishing Group.

[20] Yanay Rosen, Yusuf Roohani, Ayush Agrawal, Leon Samotorčan, Tabula Sapiens Consortium, Stephen R. Quake, and Jure Leskovec. Universal Cell Embeddings: A Foundation Model for Cell Biology, October 2024. URL https://www.biorxiv.org/content/10.1101/2023.11.28.568918v2. Pages: 2023.11.28.568918 Section: New Results.

[21] Haotian Cui, Chloe Wang, Hassaan Maan, Kuan Pang, Fengning Luo, Nan Duan, and Bo Wang. scGPT: toward building a foundation model for single-cell multi-omics using generative AI. Nature Methods, 21(8):1470–1480, August 2024. ISSN 1548-7105. doi: 10.1038/s41592-024-02201-0. URL https://www.nature.com/articles/s41592-024-02201-0. Publisher: Nature Publishing Group.

[22] Minsheng Hao, Jing Gong, Xin Zeng, Chiming Liu, Yucheng Guo, Xingyi Cheng, Taifeng Wang, Jianzhu Ma, Xuegong Zhang, and Le Song. Large-scale foundation model on single-cell transcriptomics. Nature Methods, 21(8):1481–1491, August 2024. ISSN 1548-7105. doi: 10.1038/s41592-024-02305-7. URL https://www.nature.com/articles/s41592-024-02305-7.

[23] Yixuan Wang, Yimin Fan, Xuesong Wang, Tingyang Yu, Yongshuo Zong, Xinyuan Liu, Gaoyang Zhong, Meitong Liu, Qing Li, Kin Hei Lee, Khachatur Dallakyan, Zhichao Hu, Yaqian Qi, Junjie Huang, Gengjie Jia, Jiao Yuan, Ting-Fung Chan, Xin Gao, Irwin King, and Yu Li. SCMBench: benchmarking domain-specific and foundation models for single-cell multiomics data integration. Nature Communications, May 2026. ISSN 2041-1723. doi: 10.1038/s41467-026-72570-x. URL https://www.nature.com/articles/s41467-026-72570-x.

[24] Geoffrey Hinton, Oriol Vinyals, and Jeff Dean. Distilling the Knowledge in a Neural Network, March 2015. URL http://arxiv.org/abs/1503.02531. arXiv:1503.02531 [stat].

[25] Chuanguang Yang, Zhulin An, Libo Huang, Junyu Bi, Xinqiang Yu, Han Yang, Boyu Diao, and Yongjun Xu. CLIP-KD: An Empirical Study of CLIP Model Distillation, May 2024. URL http://arxiv.org/abs/2307.12732. arXiv:2307.12732.

[26] Žiga Avsec, Natasha Latysheva, Jun Cheng, Guido Novati, Kyle R. Taylor, Tom Ward, Clare Bycroft, Lauren Nicolaisen, Eirini Arvaniti, Joshua Pan, Raina Thomas, Vincent Dutordoir, Matteo Perino, Soham De, Alexander Karollus, Adam Gayoso, Toby Sargeant, Anne Mottram, Lai Hong Wong, Pavol Drotar, Adam Kosiorek, Andrew Senior, Richard Tanburn, Taylor Applebaum, Souradeep Basu, Demis Hassabis, and Pushmeet Kohli. AlphaGenome: advancing regulatory variant effect prediction with a unified DNA sequence model, June 2025. URL https://www.biorxiv.org/content/10.1101/2025.06.25.661532v1. Pages: 2025.06.25.661532 Section: New Results.

[27] Simon Grouard, Christian Esposito, Jean El Khoury, Valérie Ducret, Céline Thiriez, Loïc Herpin, Anaïs Chossegros, Caroline Hoffmann, Quentin Bayard, Genevieve Robin, Nicole Tay, Esther Baena, Mosaic Consortium, Eric Durand, Almudena Espin Perez, and Lucas Fidon. Multiple instance learning with spatial transcriptomics for interpretable patient-level predictions: application in glioblastoma, October 2025. URL https://www.biorxiv.org/content/10.1101/2025.10.13.682206v1. ISSN: 2692-8205 Pages: 2025.10.13.682206 Section: New Results.

[28] Corina Nagy, Malosree Maitra, Arnaud Tanti, Matthew Suderman, Jean-Francois Théroux, Maria Antonietta Davoli, Kelly Perlman, Volodymyr Yerko, Yu Chang Wang, Shreejoy J. Tripathy, Paul Pavlidis, Naguib Mechawar, Jiannis Ragoussis, and Gustavo Turecki. Single-nucleus transcriptomics of the prefrontal cortex in major depressive disorder implicates oligodendrocyte precursor cells and excitatory neurons. Nature Neuroscience, 23(6):771–781, June 2020. ISSN 1546-1726. doi: 10.1038/s41593-020-0621-y. URL https://www.nature.com/articles/s41593-020-0621-y. Publisher: Nature Publishing Group.

[29] Anjali Chawla, Doruk Cakmakci, Laura M. Fiori, Wenmin Zang, Malosree Maitra, Jennie Yang, Dariusz Ż urawek, Gabriella Frosi, Reza Rahimian, Haruka Mitsuhashi, Maria Antonietta Davoli, Ryan Denniston, Gary Gang Chen, Volodymyr Yerko, Deborah Mash, Kiran Girdhar, Schahram Akbarian, Naguib Mechawar, Matthew Suderman, Yue Li, Corina Nagy, and Gustavo Turecki. Single-nucleus chromatin accessibility profiling identifies cell types and functional variants contributing to major depression. Nature Genetics, 57(8):1890–1904, August 2025. ISSN 1546-1718. doi: 10.1038/s41588-025-02249-4. URL https://www.nature.com/articles/s41588-025-02249-4. Publisher: Nature Publishing Group.

[30] Alec Radford, Jong Wook Kim, Chris Hallacy, Aditya Ramesh, Gabriel Goh, Sandhini Agarwal, Girish Sastry, Amanda Askell, Pamela Mishkin, Jack Clark, Gretchen Krueger, and Ilya Sutskever. Learning Transferable Visual Models From Natural Language Supervision, February 2021. URL http://arxiv.org/abs/2103.00020. arXiv:2103.00020 [cs].

[31] Mingbo Cheng, Zhijian Li, and Ivan G Costa. MOJITOO: a fast and universal method for integration of multimodal single-cell data. Bioinformatics, 38(Supplement 1):i282–i289, June 2022. ISSN 1367-4803. doi: 10.1093/bioinformatics/btac220. URL https://doi.org/10.1093/bioinformatics/btac220.

[32] Yinlei Hu, Siyuan Wan, Yuanhanyu Luo, Yuanzhe Li, Tong Wu, Wentao Deng, Chen Jiang, Shan Jiang, Yueping Zhang, Nianping Liu, Zongcheng Yang, Falai Chen, Bin Li, and Kun Qu. Benchmarking algorithms for single-cell multi-omics prediction and integration. Nature Methods, pages 1–13, September 2024. ISSN 1548-7105. doi: 10.1038/s41592-024-02429-w. URL https://www.nature.com/articles/s41592-024-02429-w. Publisher: Nature Publishing Group.

[33] Henry E. Pratt, Gregory Andrews, Nicole Shedd, Nishigandha Phalke, Tongxin Li, Anusri Pampari, Matthew Jensen, Cindy Wen, PsychENCODE Consortium, Michael J. Gandal, Daniel H. Geschwind, Mark Gerstein, Jill Moore, Anshul Kundaje, Andrés Colubri, and Zhiping Weng. Using a comprehensive atlas and predictive models to reveal the complexity and evolution of brain-active regulatory elements. Science Advances, 10(21):eadj4452, May 2024. doi: 10.1126/sciadv.adj4452. URL https://www.science.org/doi/10.1126/sciadv.adj4452. Publisher: American Association for the Advancement of Science.

[34] Emily M. Hicks, Carina Seah, Alanna Cote, Shelby Marchese, Kristen J. Brennand, Eric J. Nestler, Matthew J. Girgenti, and Laura M. Huckins. Integrating genetics and transcriptomics to study major depressive disorder: a conceptual framework, bioinformatic approaches, and recent findings. Translational Psychiatry, 13(1):129, April 2023. ISSN 2158-3188. doi: 10.1038/s41398-023-02412-7. URL https://www.nature.com/articles/s41398-023-02412-7. Publisher: Nature Publishing Group.

[35] Malosree Maitra, Haruka Mitsuhashi, Reza Rahimian, Anjali Chawla, Jennie Yang, Laura M. Fiori, Maria Antonietta Davoli, Kelly Perlman, Zahia Aouabed, Deborah C. Mash, Matthew Suderman, Naguib Mechawar, Gustavo Turecki, and Corina Nagy. Cell type specific transcriptomic differences in depression show similar patterns between males and females but implicate distinct cell types and genes. Nature Communications, 14(1):2912, May 2023. ISSN 2041-1723. doi: 10.1038/s41467-023-38530-5.

[36] Dmitry Velmeshev, Yonatan Perez, Zihan Yan, Jonathan E. Valencia, David R. Castaneda-Castellanos, Li Wang, Lucas Schirmer, Simone Mayer, Brittney Wick, Shaohui Wang, Tomasz Jan Nowakowski, Mercedes Paredes, Eric J. Huang, and Arnold R. Kriegstein. Single-cell analysis of prenatal and postnatal human cortical development. Science, 382(6667): eadf0834, October 2023. doi: 10.1126/science.adf0834. URL https://www.science.org/doi/10.1126/science.adf0834. Publisher: American Association for the Advancement of Science.

[37] Zhana Duren, Wenhui Sophia Lu, Joseph G. Arthur, Preyas Shah, Jingxue Xin, Francesca Meschi, Miranda Lin Li, Corey M. Nemec, Yifeng Yin, and Wing Hung Wong. Sc-compReg enables the comparison of gene regulatory networks between conditions using single-cell data. Nature Communications, 12(1):4763, August 2021. ISSN 2041-1723. doi: 10.1038/s41467-021-25089-2. URL https://www.nature.com/articles/s41467-021-25089-2. Publisher: Nature Publishing Group.

[38] Megan H. Hagenauer, Yusra Sannah, Elaine K. Hebda-Bauer, Cosette Rhoads, Angela M. O’Connor, Elizabeth Flandreau, Stanley J. Watson, and Huda Akil. Resource: A curated database of brain-related functional gene sets (Brain.GMT). MethodsX, 13:102788, December 2024. ISSN 2215-0161. doi: 10.1016/j.mex.2024.102788. URL https://www.sciencedirect.com/science/article/pii/S2215016124002413.

[39] Cory Y. McLean, Dave Bristor, Michael Hiller, Shoa L. Clarke, Bruce T. Schaar, Craig B. Lowe, Aaron M. Wenger, and Gill Bejerano. GREAT improves functional interpretation of cis-regulatory regions. Nature Biotechnology, 28(5):495–501, May 2010.ISSN 1546-1696. doi: 10.1038/nbt.1630.

[40] Nancy Y. A. Sey, Brandon M. Pratt, and Hyejung Won. Annotating genetic variants to target genes using H-MAGMA. Nature Protocols, 18(1):22–35, January 2023. ISSN 1750-2799. doi: 10.1038/s41596-022-00745-z. URL https://www.nature.com/articles/s41596-022-00745-z. Publisher: Nature Publishing Group.

[41] Jernej Godec, Yan Tan, Arthur Liberzon, Pablo Tamayo, Sanchita Bhattacharya, Atul J. Butte, Jill P. Mesirov, and W. Nicholas Haining. Compendium of immune signatures identifies conserved and species-specific biology in response to inflammation. Immunity, 44(1):194–206, January 2016. ISSN 1074-7613. doi: 10.1016/j.immuni.2015.12.006. URL https://www.ncbi.nlm.nih.gov/pmc/articles/PMC5330663/.

[42] Marta Paczkowska, Jonathan Barenboim, Nardnisa Sintupisut, Natalie S. Fox, Helen Zhu, Diala Abd-Rabbo, Miles W. Mee, Paul C. Boutros, and Jüri Reimand. Integrative pathway enrichment analysis of multivariate omics data. Nature Communications, 11:735, February 2020. ISSN 2041-1723. doi: 10.1038/s41467-019-13983-9. URL https://www.ncbi.nlm.nih.gov/pmc/articles/PMC7002665/.

[43] Butian Zhou, Zhongqun Zhu, Bruce R. Ransom, and Xiaoping Tong. Oligodendrocyte lineage cells and depression. Molecular Psychiatry, 26(1):103–117, January 2021. ISSN 1476-5578. doi: 10.1038/s41380-020-00930-0. URL https://www.nature.com/articles/s41380-020-00930-0. Publisher: Nature Publishing Group.

[44] Andrew D. Grotzinger, Josefin Werme, Wouter J. Peyrot, Oleksandr Frei, Christiaan de Leeuw, Lucy K. Bicks, Qiuyu Guo, Michael P. Margolis, Brandon J. Coombes, Anthony Batzler, Vanessa Pazdernik, Joanna M. Biernacka, Ole A. Andreassen, Verneri Anttila, Anders D. Børglum, Gerome Breen, Na Cai, Ditte Demontis, Howard J. Edenberg, Stephen V. Faraone, Barbara Franke, Michael J. Gandal, Joel Gelernter, Alexander S. Hatoum, John M. Hettema, Emma C. Johnson, Katherine G. Jonas, James A. Knowles, Karestan C. Koenen, Adam X. Maihofer, Travis T. Mallard, Manuel Mattheisen, Karen S. Mitchell, Benjamin M. Neale, Caroline M. Nievergelt, John I. Nurnberger, Kevin S. O’Connell, Roseann E. Peterson, Elise B. Robinson, Sandra S. Sanchez-Roige, Susan L. Santangelo, Jeremiah M. Scharf, Hreinn Stefansson, Kari Stefansson, Murray B. Stein, Nora I. Strom, Laura M. Thornton, Elliot M. Tucker-Drob, Brad Verhulst, Irwin D. Waldman, G. Bragi Walters, Naomi R. Wray, Dongmei Yu, Phil H. Lee, Kenneth S. Kendler, and Jordan W. Smoller. Mapping the genetic landscape across 14 psychiatric disorders. Nature, pages 1–15, December 2025. ISSN 1476-4687. doi: 10.1038/s41586-025-09820-3. URL https://www.nature.com/articles/s41586-025-09820-3. Publisher: Nature Publishing Group.

[45] Christoph Anacker, Annamaria Cattaneo, Ksenia Musaelyan, Patricia A. Zunszain, Mark Horowitz, Raffaella Molteni, Alessia Luoni, Francesca Calabrese, Katherine Tansey, Massimo Gennarelli, Sandrine Thuret, Jack Price, Rudolf Uher, Marco A. Riva, and Carmine M. Pariante. Role for the kinase SGK1 in stress, depression, and glucocorticoid effects on hippocampal neurogenesis. Proceedings of the National Academy of Sciences of the United States of America, 110(21):8708–8713, May 2013. ISSN 1091-6490. doi: 10.1073/pnas.1300886110.

[46] Suzanne Paradis, Dana B. Harrar, Yingxi Lin, Alex C. Koon, Jessica L. Hauser, Eric C. Griffith, Li Zhu, Lawrence F. Brass, Chinfei Chen, and Michael E. Greenberg. An RNAi-Based Approach Identifies Molecules Required for Glutamatergic and GABAergic Synapse Development. Neuron, 53(2):217–232, January 2007. ISSN 0896-6273. doi: 10.1016/j.neuron.2006.12.012. URL https://www.cell.com/neuron/abstract/S0896-6273(06)00997-4. Publisher: Elsevier.

[47] Daniel W. M. Acker, Irene Wong, Mihwa Kang, and Suzanne Paradis. Semaphorin 4D promotes inhibitory synapse formation and suppresses seizures in vivo. Epilepsia, 59(6): 1257–1268, June 2018. ISSN 0013-9580. doi: 10.1111/epi.14429. URL https://pmc.ncbi.nlm.nih.gov/articles/PMC5990477/.

[48] Dmitrii A. Abashkin, Dmitry S. Karpov, Artemii O. Kurishev, Ekaterina V. Marilovtseva, Vera E. Golimbet, Dmitrii A. Abashkin, Dmitry S. Karpov, Artemii O. Kurishev, Ekaterina V. Marilovtseva, and Vera E. Golimbet. ASCL1 Is Involved in the Pathogenesis of Schizophrenia by Regulation of Genes Related to Cell Proliferation, Neuronal Signature Formation, and Neuroplasticity. International Journal of Molecular Sciences, 24(21), October 2023. ISSN 1422-0067. doi: 10.3390/ijms242115746. URL https://www.mdpi.com/1422-0067/24/21/15746. Company: Multidisciplinary Digital Publishing Institute Distributor: Multidisciplinary Digital Publishing Institute Institution: Multidisciplinary Digital Publishing Institute Label: Multidisciplinary Digital Publishing Institute Publisher: publisher.

[49] Florian Duclot and Mohamed Kabbaj. The Role of Early Growth Response 1 (EGR1) in Brain Plasticity and Neuropsychiatric Disorders. Frontiers in Behavioral Neuroscience, 11, March 2017. ISSN 1662-5153. doi: 10.3389/fnbeh.2017.00035. URL https://www.frontiersin.org/journals/behavioral-neuroscience/articles/10.3389/fnbeh.2017.00035/full. Publisher: Frontiers.

[50] Somi Kim, Nam-Kyung Yu, Kyu-Won Shim, Ji-Il Kim, Hyopil Kim, Dae Hee Han, Ja Eun Choi, Seung-Woo Lee, Dong Il Choi, Myung Won Kim, Dong-Sung Lee, Kyungmin Lee, Niels Galjart, Yong-Seok Lee, Jae-Hyung Lee, and Bong-Kiun Kaang. Remote Memory and Cortical Synaptic Plasticity Require Neuronal CCCTC-Binding Factor (CTCF). The Journal of Neuroscience: The Official Journal of the Society for Neuroscience, 38(22):5042–5052, May 2018. ISSN 1529-2401. doi: 10.1523/JNEUROSCI.2738-17.2018.

[51] Dongkyeong Kim, Jin-ok Choi, Chuandong Fan, Randall S. Shearer, Mohamed Sharif, Patrick Busch, and Yungki Park. Homo-trimerization is essential for the transcription factor function of Myrf for oligodendrocyte differentiation. Nucleic Acids Research, 45(9):5112–5125, May 2017. ISSN 0305-1048. doi: 10.1093/nar/gkx080. URL https://dx.doi.org/10.1093/nar/gkx080. Publisher: Oxford Academic.

[52] João R Gomes, Andrea Lobo, Renata Nogueira, Ana F Terceiro, Susete Costelha, Igor M Lopes, Ana Magalhães, Teresa Summavielle, and Maria J Saraiva. Neuronal megalin mediates synaptic plasticity—a novel mechanism underlying intellectual disabilities in megalin gene pathologies. Brain Communications, 2(2):fcaa135, July 2020. ISSN 2632-1297. doi: 10.1093/braincomms/fcaa135. URL https://doi.org/10.1093/braincomms/fcaa135.

[53] Rebecca Favaro, Menella Valotta, Anna L. M. Ferri, Elisa Latorre, Jessica Mariani, Claudio Giachino, Cesare Lancini, Valentina Tosetti, Sergio Ottolenghi, Verdon Taylor, and Silvia K. Nicolis. Hippocampal development and neural stem cell maintenance require Sox2-dependent regulation of Shh. Nature Neuroscience, 12(10):1248–1256, October 2009. ISSN 1546-1726. doi: 10.1038/nn.2397.

[54] Zhengliang Gao, Kerstin Ure, Jessica L. Ables, Diane C. Lagace, Klaus-Armin Nave, Sandra Goebbels, Amelia J. Eisch, and Jenny Hsieh. Neurod1 is essential for the survival and maturation of adult-born neurons. Nature Neuroscience, 12(9):1090–1092, September 2009. ISSN 1546-1726. doi: 10.1038/nn.2385.

[55] Shin-Young Park and Joong-Soo Han. Phospholipase D1 Signaling: Essential Roles in Neural Stem Cell Differentiation. Journal of Molecular Neuroscience, 64(3):333–340, 2018. ISSN 0895-8696. doi: 10.1007/s12031-018-1042-1. URL https://pmc.ncbi.nlm.nih.gov/articles/PMC5874277/.

[56] Sofia I. Petersen, Rachel K. Okolicsanyi, and Larisa M. Haupt. Exploring Heparan Sulfate Proteoglycans as Mediators of Human Mesenchymal Stem Cell Neurogenesis. Cellular and Molecular Neurobiology, 44:30, March 2024. ISSN 0272-4340. doi: 10.1007/s10571-024-01463-8. URL https://pmc.ncbi.nlm.nih.gov/articles/PMC10978659/.

[57] David Tse Shen Lin and Elizabeth Conibear. ABHD17 proteins are novel protein depalmitoylases that regulate N-Ras palmitate turnover and subcellular localization. eLife, 4:e11306. ISSN 2050-084X. doi: 10.7554/eLife.11306. URL https://pmc.ncbi.nlm.nih.gov/articles/PMC4755737/.

[58] John Lonsdale, Jeffrey Thomas, Mike Salvatore, Rebecca Phillips, Edmund Lo, Saboor Shad, Richard Hasz, Gary Walters, Fernando Garcia, Nancy Young, Barbara Foster, Mike Moser, Ellen Karasik, Bryan Gillard, Kimberley Ramsey, Susan Sullivan, Jason Bridge, Harold Magazine, John Syron, Johnelle Fleming, Laura Siminoff, Heather Traino, Maghboeba Mosavel, Laura Barker, Scott Jewell, Dan Rohrer, Dan Maxim, Dana Filkins, Philip Harbach, Eddie Cortadillo, Bree Berghuis, Lisa Turner, Eric Hudson, Kristin Feenstra, Leslie Sobin, James Robb, Phillip Branton, Greg Korzeniewski, Charles Shive, David Tabor, Liqun Qi, Kevin Groch, Sreenath Nampally, Steve Buia, Angela Zimmerman, Anna Smith, Robin Burges, Karna Robinson, Kim Valentino, Deborah Bradbury, Mark Cosentino, Norma Diaz-Mayoral, Mary Kennedy, Theresa Engel, Penelope Williams, Kenyon Erickson, Kristin Ardlie, Wendy Winckler, Gad Getz, David DeLuca, Daniel MacArthur, Manolis Kellis, Alexander Thomson, Taylor Young, Ellen Gelfand, Molly Donovan, Yan Meng, George Grant, Deborah Mash, Yvonne Marcus, Margaret Basile, Jun Liu, Jun Zhu, Zhidong Tu, Nancy J. Cox, Dan L. Nicolae, Eric R. Gamazon, Hae Kyung Im, Anuar Konkashbaev, Jonathan Pritchard, Matthew Stevens, Timothèe Flutre, Xiaoquan Wen, Emmanouil T. Dermitzakis, Tuuli Lappalainen, Roderic Guigo, Jean Monlong, Michael Sammeth, Daphne Koller, Alexis Battle, Sara Mostafavi, Mark McCarthy, Manual Rivas, Julian Maller, Ivan Rusyn, Andrew Nobel, Fred Wright, Andrey Shabalin, Mike Feolo, Nataliya Sharopova, Anne Sturcke, Justin Paschal, James M. Anderson, Elizabeth L. Wilder, Leslie K. Derr, Eric D. Green, Jeffery P. Struewing, Gary Temple, Simona Volpi, Joy T. Boyer, Elizabeth J. Thomson, Mark S. Guyer, Cathy Ng, Assya Abdallah, Deborah Colantuoni, Thomas R. Insel, Susan E. Koester, A. Roger Little, Patrick K. Bender, Thomas Lehner, Yin Yao, Carolyn C. Compton, Jimmie B. Vaught, Sherilyn Sawyer, Nicole C. Lockhart, Joanne Demchok, and Helen F. Moore. The Genotype-Tissue Expression (GTEx) project. Nature Genetics, 45(6):580–585, June 2013. ISSN 1546-1718. doi: 10.1038/ng.2653. URL https://www.nature.com/articles/ng.2653. Publisher: Nature Publishing Group.

[59] Yong He, Yue Wang, Heming Yu, Yu Tian, Xiangyu Chen, Chong Chen, Yikun Ren, Zhi Chen, Yi Ren, Xue Gong, Ke Cheng, Xiaolei Liu, Lianmei Zhong, Yi Guo, and Peng Xie. Protective effect of Nr4a2 (Nurr1) against LPS-induced depressive-like behaviors via regulating activity of microglia and CamkII neurons in anterior cingulate cortex. Pharmacological Research, 191:106717, May 2023. ISSN 1043-6618. doi: 10.1016/j.phrs.2023.106717. URL https://www.sciencedirect.com/science/article/pii/S1043661823000737.

[60] Ryne C. Ramaker, Kevin M. Bowling, Brittany N. Lasseigne, Megan H. Hagenauer, Andrew A. Hardigan, Nicholas S. Davis, Jason Gertz, Preston M. Cartagena, David M. Walsh, Marquis P. Vawter, Edward G. Jones, Alan F. Schatzberg, Jack D. Barchas, Stanley J. Watson, Blynn G. Bunney, Huda Akil, William E. Bunney, Jun Z. Li, Sara J. Cooper, and Richard M. Myers. Post-mortem molecular profiling of three psychiatric disorders. Genome Medicine, 9(1):72, July 2017. ISSN 1756-994X. doi: 10.1186/s13073-017-0458-5. URL https://doi.org/10.1186/s13073-017-0458-5.

[61] Sheng Zhang, Xiaoqing Zhu, Xuehong Gui, Christopher Croteau, Lanying Song, Jie Xu, Aijun Wang, Peter Bannerman, and Fuzheng Guo. Sox2 Is Essential for Oligodendroglial Proliferation and Differentiation during Postnatal Brain Myelination and CNS Remyelination. Journal of Neuroscience, 38(7):1802–1820, February 2018. ISSN 0270-6474, 1529-2401. doi: 10.1523/JNEUROSCI.1291-17.2018. URL https://www.jneurosci.org/content/38/7/1802. Publisher: Society for Neuroscience Section: Research Articles.

[62] Benedetta Foglio, Laura Rossini, Rita Garbelli, Maria Cristina Regondi, Sara Mercurio, Michele Bertacchi, Laura Avagliano, Gaetano Bulfamante, Roland Coras, Antonino Maiorana, Silvia Nicolis, Michèle Studer, and Carolina Frassoni. Dynamic expression of NR2F1 and SOX2 in developing and adult human cortex: comparison with cortical malformations. Brain Structure & Function, 226(4):1303–1322, May 2021. ISSN 1863-2661. doi: 10.1007/s00429-021-02242-7.

[63] Kaiyi Zhu, Jaroslav Bendl, Samir Rahman, James M. Vicari, Claire Coleman, Tereza Clarence, Ovaun Latouche, Nadejda M. Tsankova, Aiqun Li, Kristen J. Brennand, Donghoon Lee, Guo-Cheng Yuan, John F. Fullard, and Panos Roussos. Multi-omic profiling of the developing human cerebral cortex at the single-cell level. Science Advances, 9(41):eadg3754, October 2023. doi: 10.1126/sciadv.adg3754. URL https://www.science.org/doi/10.1126/sciadv.adg3754. Publisher: American Association for the Advancement of Science.

[64] Xi Wang, Qiwei Lian, Haoyu Dong, Shuo Xu, Yaru Su, and Xiaohui Wu. Benchmarking Algorithms for Gene Set Scoring of Single-cell ATAC-seq Data. Genomics, Proteomics & Bioinformatics, 22(2):qzae014, April 2024. ISSN 1672-0229. doi: 10.1093/gpbjnl/qzae014. URL https://doi.org/10.1093/gpbjnl/qzae014.

[65] Li Wang, Cheng Wang, Juan A. Moriano, Songcang Chen, Guolong Zuo, Arantxa Cebriáan-Silla, Shaobo Zhang, Tanzila Mukhtar, Shaohui Wang, Mengyi Song, Lilian Gomes de Oliveira, Qiuli Bi, Jonathan J. Augustin, Xinxin Ge, Mercedes F. Paredes, Eric J. Huang, Arturo Alvarez-Buylla, Xin Duan, Jingjing Li, and Arnold R. Kriegstein. Molecular and cellular dynamics of the developing human neocortex. Nature, pages 1–10, January 2025. ISSN 1476-4687. doi: 10.1038/s41586-024-08351-7. URL https://www.nature.com/articles/s41586-024-08351-7. Publisher: Nature Publishing Group.

[66] F. Alexander Wolf, Fiona K. Hamey, Mireya Plass, Jordi Solana, Joakim S. Dahlin, Berthold Göttgens, Nikolaus Rajewsky, Lukas Simon, and Fabian J. Theis. PAGA: graph abstraction reconciles clustering with trajectory inference through a topology preserving map of single cells. Genome Biology, 20(1):59, March 2019. ISSN 1474-760X. doi: 10.1186/s13059-019-1663-x. URL https://doi.org/10.1186/s13059-019-1663-x.

[67] Laleh Haghverdi, Maren Büttner, F. Alexander Wolf, Florian Buettner, and Fabian J. Theis. Diffusion pseudotime robustly reconstructs lineage branching. Nature Methods, 13(10):845–848, October 2016. ISSN 1548-7105. doi: 10.1038/nmeth.3971. URL https://www.nature.com/articles/nmeth.3971. Publisher: Nature Publishing Group.

[68] Wenzhi Cao, Vahid Mirjalili, and Sebastian Raschka. Rank consistent ordinal regression for neural networks with application to age estimation. Pattern Recognition Letters, 140: 325–331, December 2020. ISSN 0167-8655. doi: 10.1016/j.patrec.2020.11.008. URL https://www.sciencedirect.com/science/article/pii/S016786552030413X.

[69] Yanling Wang, Catherine A. Dye, Vikaas Sohal, Jason E. Long, Rosanne C. Estrada, Tomas Roztocil, Thomas Lufkin, Karl Deisseroth, Scott C. Baraban, and John L. R. Rubenstein. Dlx5 and Dlx6 Regulate the Development of Parvalbumin-Expressing Cortical Interneurons. Journal of Neuroscience, 30(15):5334–5345, April 2010.ISSN 0270-6474, 1529-2401. doi: 10.1523/JNEUROSCI.5963-09.2010. URL https://www.jneurosci.org/content/30/15/5334. Publisher: Society for Neuroscience Section: Articles.

[70] Camille de Lombares, Eglantine Heude, Gladys Alfama, Anastasia Fontaine, Rim Hassouna, Cécile Vernochet, Fabrice de Chaumont, Christophe Olivo-Marin, Elodie Ey, Sébastien Parnaudeau, François Tronche, Thomas Bourgeron, Serge Luquet, Giovanni Levi, and Nicolas Narboux-Nême. *Dlx5* and *Dlx6* expression in GABAergic neurons controls behavior, metabolism, healthy aging and lifespan. Aging, 11(17):6638–6656, September 2019. ISSN 1945-4589. doi: 10.18632/aging.102141. URL https://www.aging-us.com/article/102141/text.

[71] Silvia Domcke, Andrew J. Hill, Riza M. Daza, Junyue Cao, Diana R. O’Day, Hannah A. Pliner, Kimberly A. Aldinger, Dmitry Pokholok, Fan Zhang, Jennifer H. Milbank, Michael A. Zager, Ian A. Glass, Frank J. Steemers, Dan Doherty, Cole Trapnell, Darren A. Cusanovich, and Jay Shendure. A human cell atlas of fetal chromatin accessibility. Science, 370(6518): eaba7612, November 2020. doi: 10.1126/science.aba7612. URL https://www.science.org/doi/10.1126/science.aba7612. Publisher: American Association for the Advancement of Science.

[72] Charles A. Herring, Rebecca K. Simmons, Saskia Freytag, Daniel Poppe, Joel J. D. Moffet, Jahnvi Pflueger, Sam Buckberry, Dulce B. Vargas-Landin, Olivier Clément, Enrique Goñi Echeverría, Gavin J. Sutton, Alba Alvarez-Franco, Rui Hou, Christian Pflueger, Kerrie McDonald, Jose M. Polo, Alistair R. R. Forrest, Anna K. Nowak, Irina Voineagu, Luciano Martelotto, and Ryan Lister. Human prefrontal cortex gene regulatory dynamics from gestation to adulthood at single-cell resolution. Cell, 185(23):4428–4447.e28, November 2022. ISSN 0092-8674. doi: 10.1016/j.cell.2022.09.039. URL https://www.sciencedirect.com/science/article/pii/S0092867422012582.

[73] Jessica L. Bolton, Annabel K. Short, Shivashankar Othy, Cassandra L. Kooiker, Manlin Shao, Benjamin G. Gunn, Jaclyn Beck, Xinglong Bai, Stephanie M. Law, Julie C. Savage, Jeremy J. Lambert, Delia Belelli, Marie-Ève Tremblay, Michael D. Cahalan, and Tallie Z. Baram. Early stress-induced impaired microglial pruning of excitatory synapses on immature CRH-expressing neurons provokes aberrant adult stress responses. Cell Reports, 38(13), March 2022. ISSN 2211-1247. doi: 10.1016/j.celrep.2022.110600. URL https://www.cell.com/cell-reports/abstract/S2211-1247(22)00348-5. Publisher: Elsevier.

[74] Anna S. Fröhlich, Nathalie Gerstner, Miriam Gagliardi, Maik Ködel, Natan Yusupov, Natalie Matosin, Darina Czamara, Susann Sauer, Simone Roeh, Vanessa Murek, Chris Chatzinakos, Nikolaos P. Daskalakis, Janine Knauer-Arloth, Michael J. Ziller, and Elisabeth B. Binder. Single-nucleus transcriptomic profiling of human orbitofrontal cortex reveals convergent effects of aging and psychiatric disease. Nature Neuroscience, 27(10):2021–2032, October 2024. ISSN 1546-1726. doi: 10.1038/s41593-024-01742-z. URL https://www.nature.com/articles/s41593-024-01742-z. Publisher: Nature Publishing Group.

[75] Natan Yusupov, Linda Dieckmann, Mira Erhart, Susann Sauer, Monika Rex-Haffner, Johannes Kopf-Beck, Tanja M. Brückl, Darina Czamara, and Elisabeth B. Binder. Transdiagnostic evaluation of epigenetic age acceleration and burden of psychiatric disorders. Neuropsychopharmacology, 48(9):1409–1417, August 2023. ISSN 1740-634X. doi: 10.1038/s41386-023-01579-3. URL https://www.nature.com/articles/s41386-023-01579-3. Publisher: Nature Publishing Group.

[76] Boris Muzellec, Maria Teleńczuk, Vincent Cabeli, and Mathieu Andreux. PyDESeq2: a python package for bulk RNA-seq differential expression analysis. Bioinformatics, 39(9): btad547, September 2023. ISSN 1367-4803. doi: 10.1093/bioinformatics/btad547. URL https://www.ncbi.nlm.nih.gov/pmc/articles/PMC10502239/.

[77] Alicia N. Schep, Beijing Wu, Jason D. Buenrostro, and William J. Greenleaf. chromVAR: inferring transcription-factor-associated accessibility from single-cell epigenomic data. Nature Methods, 14(10):975–978, October 2017. ISSN 1548-7105. doi: 10.1038/nmeth.4401. URL https://www.nature.com/articles/nmeth.4401. Publisher: Nature Publishing Group.

[78] Patrick O. McGowan, Aya Sasaki, Ana C. D’Alessio, Sergiy Dymov, Benoit Labonté, Moshe Szyf, Gustavo Turecki, and Michael J. Meaney. Epigenetic regulation of the glucocorticoid receptor in human brain associates with childhood abuse. Nature Neuroscience, 12(3):342–348, March 2009. ISSN 1546-1726. doi: 10.1038/nn.2270. URL https://www.nature.com/articles/nn.2270. Publisher: Nature Publishing Group.

[79] Heonjong Han, Jae-Won Cho, Sangyoung Lee, Ayoung Yun, Hyojin Kim, Dasom Bae, Sunmo Yang, Chan Yeong Kim, Muyoung Lee, Eunbeen Kim, Sungho Lee, Byunghee Kang, Dabin Jeong, Yaeji Kim, Hyeon-Nae Jeon, Haein Jung, Sunhwee Nam, Michael Chung, Jong-Hoon Kim, and Insuk Lee. TRRUST v2: an expanded reference database of human and mouse transcriptional regulatory interactions. Nucleic Acids Research, 46(D1):D380–D386, January 2018. ISSN 0305-1048. doi: 10.1093/nar/gkx1013. URL https://doi.org/10.1093/nar/gkx1013.

[80] Ilan A. Kerman, René Bernard, William Bunney, Edward Jones, Alan F. Schatzberg, Richard Myers, Jack Barchas, Huda Akil, Stanley Watson, and Robert Thompson. Evidence for Transcriptional Factor Dysregulation in the Dorsal Raphe Nucleus of Patients with Major Depressive Disorder. Frontiers in Neuroscience, 6, October 2012. ISSN 1662-453X. doi: 10.3389/fnins.2012.00135. URL https://www.frontiersin.org/journals/neuroscience/articles/10.3389/fnins.2012.00135/full. Publisher: Frontiers.

[81] Giovana Bristot, Jacson Gabriel Feiten, Bianca Pfaffenseller, Gabriel Henrique Hizo, Gabriela Maria Pereira Possebon, Fernanda Endler Valiati, Jairo Vinícius Pinto, Marco Antonio Caldieraro, Marcelo Pio de Almeida Fleck, Clarissa Severino Gama, and Márcia Kauer-Sant’Anna. Early growth response 1 (EGR1) is downregulated in peripheral blood from patients with major psychiatric disorders. Trends in Psychiatry and Psychotherapy, 46: e20230749, November 2024. ISSN 2237-6089. doi: 10.47626/2237-6089-2023-0749. URL https://www.ncbi.nlm.nih.gov/pmc/articles/PMC11815351/.

[82] Stephen C. Gammie. Creation of a gene expression portrait of depression and its application for identifying potential treatments. Scientific Reports, 11(1):3829, February 2021. ISSN 2045-2322. doi: 10.1038/s41598-021-83348-0. URL https://www.nature.com/articles/s41598-021-83348-0. Publisher: Nature Publishing Group.

[83] Jo-Fan Chien, Hanqing Liu, Bang-An Wang, Chongyuan Luo, Anna Bartlett, Rosa Castanon, Nicholas D. Johnson, Joseph R. Nery, Julia Osteen, Junhao Li, Jordan Altshul, Mia Kenworthy, Cynthia Valadon, Michelle Liem, Naomi Claffey, Carolyn O’Connor, Luise A. Seeker, Joseph R. Ecker, M. Margarita Behrens, and Eran A. Mukamel. Cell-type-specific effects of age and sex on human cortical neurons. Neuron, 112(15):2524–2539.e5, August 2024. ISSN 0896-6273. doi: 10.1016/j.neuron.2024.05.013. URL https://www.cell.com/neuron/abstract/S0896-6273(24)00360-X. Publisher: Elsevier.

[84] Yanay Rosen, Maria Brbić, Yusuf Roohani, Kyle Swanson, Ziang Li, and Jure Leskovec. Toward universal cell embeddings: integrating single-cell RNA-seq datasets across species with SATURN. Nature Methods, 21(8):1492–1500, August 2024. ISSN 1548-7105. doi: 10.1038/s41592-024-02191-z. URL https://www.nature.com/articles/s41592-024-02191-z. Publisher: Nature Publishing Group.

[85] Tim Stuart, Avi Srivastava, Shaista Madad, Caleb A. Lareau, and Rahul Satija. Single-cell chromatin state analysis with Signac. Nature Methods, 18(11):1333–1341, November 2021. ISSN 1548-7105. doi: 10.1038/s41592-021-01282-5. URL https://www.nature.com/articles/s41592-021-01282-5. Publisher: Nature Publishing Group.

[86] Pavan Kumar Anasosalu Vasu, Hadi Pouransari, Fartash Faghri, Raviteja Vemulapalli, and Oncel Tuzel. MobileCLIP: Fast Image-Text Models through Multi-Modal Reinforced Training, April 2024. URL http://arxiv.org/abs/2311.17049. arXiv:2311.17049.

[87] Rémi Flamary, Nicolas Courty, Alexandre Gramfort, Mokhtar Z. Alaya, Aurélie Boisbunon, Stanislas Chambon, Laetitia Chapel, Adrien Corenflos, Kilian Fatras, Nemo Fournier, Léo Gautheron, Nathalie T. H. Gayraud, Hicham Janati, Alain Rakotomamonjy, Ievgen Redko, Antoine Rolet, Antony Schutz, Vivien Seguy, Danica J. Sutherland, Romain Tavenard, Alexander Tong, and Titouan Vayer. POT: Python Optimal Transport. Journal of Machine Learning Research, 22(78):1–8, 2021. ISSN 1533-7928. URL http://jmlr.org/papers/v22/20-451.html.

[88] Gabriel Peyré and Marco Cuturi. Computational Optimal Transport, March 2020. URL http://arxiv.org/abs/1803.00567. arXiv:1803.00567 [stat].

[89] Takuya Akiba, Shotaro Sano, Toshihiko Yanase, Takeru Ohta, and Masanori Koyama. Optuna: A Next-generation Hyperparameter Optimization Framework, July 2019. URL http://arxiv.org/abs/1907.10902. arXiv:1907.10902 [cs].

[90] Adam Paszke, Sam Gross, Francisco Massa, Adam Lerer, James Bradbury, Gregory Chanan, Trevor Killeen, Zeming Lin, Natalia Gimelshein, Luca Antiga, Alban Desmaison, Andreas Köpf, Edward Yang, Zach DeVito, Martin Raison, Alykhan Tejani, Sasank Chilamkurthy, Benoit Steiner, Lu Fang, Junjie Bai, and Soumith Chintala. PyTorch: an imperative style, high-performance deep learning library. In Proceedings of the 33rd International Conference on Neural Information Processing Systems, number 721, pages 8026–8037. Curran Associates Inc., Red Hook, NY, USA, 2019.

[91] Ryan K. Dale, Brent S. Pedersen, and Aaron R. Quinlan. Pybedtools: a flexible Python library for manipulating genomic datasets and annotations. Bioinformatics, 27(24):3423–3424, December 2011. ISSN 1367-4803. doi: 10.1093/bioinformatics/btr539. URL https://doi.org/10.1093/bioinformatics/btr539.

[92] Ashlyn G. Anderson, Brianne B. Rogers, Jacob M. Loupe, Ivan Rodriguez-Nunez, Sydney C. Roberts, Lauren M. White, J. Nicholas Brazell, William E. Bunney, Blynn G. Bunney, Stanley J. Watson, J. Nicholas Cochran, Richard M. Myers, and Lindsay F. Rizzardi. Single nucleus multiomics identifies ZEB1 and MAFB as candidate regulators of Alzheimer’s disease-specific cis-regulatory elements. Cell Genomics, 3(3), March 2023. ISSN 2666-979X. doi: 10.1016/j.xgen.2023.100263. URL https://www.cell.com/cell-genomics/abstract/S2666-979X(23)00019-8. Publisher: Elsevier.

[93] Levi Adams, Min Kyung Song, Samantha Yuen, Yoshiaki Tanaka, and Yoon-Seong Kim. A single-nuclei paired multiomic analysis of the human midbrain reveals age- and Parkinson’s disease–associated glial changes. Nature aging, 4(3):364–378, March 2024. ISSN 2662-8465. doi: 10.1038/s43587-024-00583-6. URL https://www.ncbi.nlm.nih.gov/pmc/articles/PMC11361719/.

[94] Shaojie Ma, Mario Skarica, Qian Li, Chuan Xu, Ryan D. Risgaard, Andrew T. N. Tebbenkamp, Xoel Mato-Blanco, Rothem Kovner, Željka Krsnik, Xabier De Martin, Victor Luria, Xavier Martí-Pérez, Dan Liang, Amir Karger, Danielle K. Schmidt, Zachary Gomez-Sanchez, Cai Qi, Kevin T. Gobeske, Sirisha Pochareddy, Ashwin Debnath, Cade J. Hottman, Joshua Spurrier, Leon Teo, Anthony G. Boghdadi, Jihane Homman-Ludiye, John J. Ely, Etienne W. Daadi, Da Mi, Marcel Daadi, Oscar Marín, Patrick R. Hof, Mladen-Roko Rasin, James Bourne, Chet C. Sherwood, Gabriel Santpere, Matthew J. Girgenti, Stephen M. Strittmatter, André M. M. Sousa, and Nenad Sestan. Molecular and cellular evolution of the primate dorsolateral prefrontal cortex. Science, 377(6614):eabo7257, September 2022. ISSN 0036-8075, 1095-9203. doi: 10.1126/science.abo7257. URL https://www.science.org/doi/10.1126/science.abo7257.

[95] Malte D. Luecken, M. Büttner, K. Chaichoompu, A. Danese, M. Interlandi, M. F. Mueller, D. C. Strobl, L. Zappia, M. Dugas, M. Colomé-Tatché, and Fabian J. Theis. Benchmarking atlas-level data integration in single-cell genomics. Nature Methods, 19(1):41–50, January 2022. ISSN 1548-7105. doi: 10.1038/s41592-021-01336-8. URL https://www.nature.com/articles/s41592-021-01336-8. Publisher: Nature Publishing Group.

[96] Laleh Haghverdi, Florian Buettner, and Fabian J. Theis. Diffusion maps for high-dimensional single-cell analysis of differentiation data. Bioinformatics, 31(18):2989–2998, September 2015. ISSN 1367-4803. doi: 10.1093/bioinformatics/btv325. URL https://doi.org/10.1093/bioinformatics/btv325.

[97] Maya U. Sheth, Wei-Lin Qiu, X. Rosa Ma, Andreas R. Gschwind, Evelyn Jagoda, Anthony S. Tan, Hjörleifur Einarsson, Bram L. Gorissen, Danilo Dubocanin, Christopher S. McGinnis, Dulguun Amgalan, Ansuman T. Satpathy, Thouis R. Jones, Lars M. Steinmetz, Anshul Kundaje, Berk Ustun, Jesse M. Engreitz, and Robin Andersson. Mapping enhancer-gene regulatory interactions from single-cell data, November 2024. URL https://www.biorxiv.org/content/10.1101/2024.11.23.624931v1. Pages: 2024.11.23.624931 Section: New Results.

[98] Jordan W. Squair, Matthieu Gautier, Claudia Kathe, Mark A. Anderson, Nicholas D. James, Thomas H. Hutson, Rémi Hudelle, Taha Qaiser, Kaya J. E. Matson, Quentin Barraud, Ariel J. Levine, Gioele La Manno, Michael A. Skinnider, and Grégoire Courtine. Confronting false discoveries in single-cell differential expression. Nature Communications, 12(1):5692, September 2021. ISSN 2041-1723. doi: 10.1038/s41467-021-25960-2. URL https://www.nature.com/articles/s41467-021-25960-2. Publisher: Nature Publishing Group.

[99] Sitara Persad, Zi-Ning Choo, Christine Dien, Noor Sohail, Ignas Masilionis, Ronan Chaligné, Tal Nawy, Chrysothemis C. Brown, Roshan Sharma, Itsik Pe’er, Manu Setty, and Dana Pe’er. SEACells infers transcriptional and epigenomic cellular states from single-cell genomics data. Nature Biotechnology, 41(12):1746–1757, December 2023. ISSN 1546-1696. doi: 10.1038/s41587-023-01716-9. URL https://www.nature.com/articles/s41587-023-01716-9. Publisher: Nature Publishing Group.

[100] Jeffrey Niu and Jiarui Ding. Single-cell multiomics data integration and generation with scPairing, January 2025. URL https://www.biorxiv.org/content/10.1101/2025.01.04.631299v1. Pages: 2025.01.04.631299 Section: New Results.

[101] Aziz Khan. pyJASPAR: a Pythonic interface to JASPAR transcription factor motifs, February 2021. URL https://zenodo.org/records/4509415.

[102] Endre Bakken Stovner and Pål Sætrom. PyRanges: efficient comparison of genomic intervals in Python. Bioinformatics, 36(3):918–919, February 2020. ISSN 1367-4803. doi: 10.1093/bioinformatics/btz615. URL https://doi.org/10.1093/bioinformatics/btz615.

[103] Prashant S. Emani, Jason J. Liu, Declan Clarke, Matthew Jensen, Jonathan Warrell, Chirag Gupta, Ran Meng, Che Yu Lee, Siwei Xu, Cagatay Dursun, Shaoke Lou, Yuhang Chen, Zhiyuan Chu, Timur Galeev, Ahyeon Hwang, Yunyang Li, Pengyu Ni, Xiao Zhou, PsychENCODE Consortium, Trygve E. Bakken, Jaroslav Bendl, Lucy Bicks, Tanima Chatterjee, Lijun Cheng, Yuyan Cheng, Yi Dai, Ziheng Duan, Mary Flaherty, John F. Fullard, Michael Gancz, Diego Garrido-Martín, Sophia Gaynor-Gillett, Jennifer Grundman, Natalie Hawken, Ella Henry, Gabriel E. Hoffman, Ao Huang, Yunzhe Jiang, Ting Jin, Nikolas L. Jorstad, Riki Kawaguchi, Saniya Khullar, Jianyin Liu, Junhao Liu, Shuang Liu, Shaojie Ma, Michael Margolis, Samantha Mazariegos, Jill Moore, Jennifer R. Moran, Eric Nguyen, Nishigandha Phalke, Milos Pjanic, Henry Pratt, Diana Quintero, Ananya S. Rajagopalan, Tiernon R. Riesenmy, Nicole Shedd, Manman Shi, Megan Spector, Rosemarie Terwilliger, Kyle J. Travaglini, Brie Wamsley, Gaoyuan Wang, Yan Xia, Shaohua Xiao, Andrew C. Yang, Suchen Zheng, Michael J. Gandal, Donghoon Lee, Ed S. Lein, Panos Roussos, Nenad Sestan, Zhiping Weng, Kevin P. White, Hyejung Won, Matthew J. Girgenti, Jing Zhang, Daifeng Wang, Daniel Geschwind, and Mark Gerstein. Single-cell genomics and regulatory networks for 388 human brains. Science, 384 (6698):eadi5199, May 2024. doi: 10.1126/science.adi5199. URL https://www.science.org/doi/10.1126/science.adi5199. Publisher: American Association for the Advancement of Science.

[104] Edward Y. Chen, Christopher M. Tan, Yan Kou, Qiaonan Duan, Zichen Wang, Gabriela Vaz Meirelles, Neil R. Clark, and Avi Ma’ayan. Enrichr: interactive and collaborative HTML5 gene list enrichment analysis tool. BMC Bioinformatics, 14(1):128, April 2013. ISSN 1471-2105. doi: 10.1186/1471-2105-14-128. URL https://doi.org/10.1186/1471-2105-14-128.

[105] Zhuoqing Fang, Xinyuan Liu, and Gary Peltz. GSEApy: a comprehensive package for performing gene set enrichment analysis in Python. Bioinformatics, 39(1):btac757, January 2023. ISSN 1367-4811. doi: 10.1093/bioinformatics/btac757. URL https://doi.org/10.1093/bioinformatics/btac757.

[106] Toni Q. Duan, Megan H. Hagenauer, Elizabeth I. Flandreau, Anne Bader, Duy Manh Nguyen, Pamela M. Maras, Randriely Merscher S. De Lima, Trevonn Gyles, Christabel Mclain, Michael. J. Meaney, Eric J. Nestler, Stanley J. Watson, and Huda Akil. A meta-analysis of the effects of early life stress on the prefrontal cortex transcriptome suggests long-term effects on myelin. bioRxiv, page 2024.11.22.624315, November 2024. ISSN 2692-8205. doi: 10.1101/2024.11.22.624315. URL https://www.ncbi.nlm.nih.gov/pmc/articles/PMC11601536/.

[107] Skipper Seabold and Josef Perktold. statsmodels: Econometric and statistical modeling with python. In statsmodels: Econometric and statistical modeling with python, 2010.

[108] V. Batagelj and M. Zaversnik. An O(m) Algorithm for Cores Decomposition of Networks, October 2003. URL http://arxiv.org/abs/cs/0310049. arXiv:cs/0310049.

[109] Aric A. Hagberg, Daniel A. Schult, and Pieter J. Swart. Exploring Network Structure, Dynamics, and Function using NetworkX. pages 11–15, Pasadena, California, June 2008. doi: 10.25080/TCWV9851. URL https://doi.curvenote.com/10.25080/TCWV9851.

[110] Jean-Luc R. Stevens, Philipp Rudiger, and James A. Bednar. HoloViews: Building Complex Visualizations Easily for Reproducible Science. SciPy 2015, June 2015. doi: 10.25080/Majora-7b98e3ed-00a. URL https://proceedings.scipy.org/articles/Majora-7b98e3ed-00a.

[111] F. Alexander Wolf, Philipp Angerer, and Fabian J. Theis. SCANPY: large-scale single-cell gene expression data analysis. Genome Biology, 19(1):15, February 2018. ISSN 1474-760X. doi: 10.1186/s13059-017-1382-0. URL https://doi.org/10.1186/s13059-017-1382-0.

[112] Will Macnair, Revant Gupta, and Manfred Claassen. psupertime: supervised pseudotime analysis for time-series single-cell RNA-seq data. Bioinformatics, 38(Supplement 1):i290–i298, June 2022. ISSN 1367-4803. doi: 10.1093/bioinformatics/btac227. URL https://doi.org/10.1093/bioinformatics/btac227.

[113] Carmen Bravo González-Blas, Seppe De Winter, Gert Hulselmans, Nikolai Hecker, Irina Matetovici, Valerie Christiaens, Suresh Poovathingal, Jasper Wouters, Sara Aibar, and Stein Aerts. SCENIC+: single-cell multiomic inference of enhancers and gene regulatory networks. Nature Methods, 20(9):1355–1367, September 2023. ISSN 1548-7105. doi: 10.1038/s41592-023-01938-4. URL https://www.nature.com/articles/s41592-023-01938-4. Publisher: Nature Publishing Group.

[114] Carmen Bravo Gonzáalez-Blas, Liesbeth Minnoye, Dafni Papasokrati, Sara Aibar, Gert Hulselmans, Valerie Christiaens, Kristofer Davie, Jasper Wouters, and Stein Aerts. cisTopic: cis-regulatory topic modeling on single-cell ATAC-seq data. Nature Methods, 16(5):397–400, May 2019. ISSN 1548-7105. doi: 10.1038/s41592-019-0367-1. URL https://www.nature.com/articles/s41592-019-0367-1. Publisher: Nature Publishing Group.

[115] The pandas development team. pandas-dev/pandas: Pandas, September 2025. URL https://zenodo.org/doi/10.5281/zenodo.3509134.

[116] Wes McKinney. Data Structures for Statistical Computing in Python. pages 56–61, Austin, Texas, 2010. doi: 10.25080/Majora-92bf1922-00a. URL https://doi.curvenote.com/10.25080/Majora-92bf1922-00a.

[117] Stéfan van der Walt, Johannes L. Schönberger, Juan Nunez-Iglesias, François Boulogne, Joshua D. Warner, Neil Yager, Emmanuelle Gouillart, and Tony Yu. scikit-image: image processing in Python. PeerJ, 2:e453, June 2014. ISSN 2167-8359. doi: 10.7717/peerj.453. URL https://peerj.com/articles/453. Publisher: PeerJ Inc.

[118] Junyue Cao, Malte Spielmann, Xiaojie Qiu, Xingfan Huang, Daniel M. Ibrahim, Andrew J. Hill, Fan Zhang, Stefan Mundlos, Lena Christiansen, Frank J. Steemers, Cole Trapnell, and Jay Shendure. The single-cell transcriptional landscape of mammalian organogenesis. Nature, 566(7745):496–502, February 2019. ISSN 1476-4687. doi: 10.1038/s41586-019-0969-x. URL https://www.nature.com/articles/s41586-019-0969-x. Publisher: Nature Publishing Group.

[119] Sergio J Rey and Luc Anselin. A.10 PySAL: A Python Library of Spatial Analytical Methods.

[120] Daniel Servén and Charlie Brummitt. pyGAM: Generalized Additive Models in Python, March 2018. URL https://zenodo.org/record/1208724.

[121] Fabian Pedregosa, Gaël Varoquaux, Alexandre Gramfort, Vincent Michel, Bertrand Thirion, Olivier Grisel, Mathieu Blondel, Peter Prettenhofer, Ron Weiss, Vincent Dubourg, Jake Vanderplas, Alexandre Passos, David Cournapeau, Matthieu Brucher, Matthieu Perrot, and Édouard Duchesnay. Scikit-learn: Machine Learning in Python. Journal of Machine Learning Research, 12(85):2825–2830, 2011. ISSN 1533-7928. URL http://jmlr.org/papers/v12/pedregosa11a.html.

[122] Isaac Virshup, Danila Bredikhin, Lukas Heumos, Giovanni Palla, Gregor Sturm, Adam Gayoso, Ilia Kats, Mikaela Koutrouli, Bonnie Berger, Dana Pe’er, Aviv Regev, Sarah A. Teichmann, Francesca Finotello, F. Alexander Wolf, Nir Yosef, Oliver Stegle, and Fabian J. Theis. The scverse project provides a computational ecosystem for single-cell omics data analysis. Nature Biotechnology, 41(5):604–606, May 2023. ISSN 1546-1696. doi: 10.1038/s41587-023-01733-8. URL https://www.nature.com/articles/s41587-023-01733-8. Publisher: Nature Publishing Group.

[123] Zuguang Gu and Daniel Hübschmann. rGREAT: an R/bioconductor package for functional enrichment on genomic regions. Bioinformatics, 39(1):btac745, January 2023. ISSN 1367-4811. doi: 10.1093/bioinformatics/btac745. URL https://doi.org/10.1093/bioinformatics/btac745.

[124] David M. Howard, Mark J. Adams, Toni-Kim Clarke, Jonathan D. Hafferty, Jude Gibson, Masoud Shirali, Jonathan R. I. Coleman, Saskia P. Hagenaars, Joey Ward, Eleanor M. Wigmore, Clara Alloza, Xueyi Shen, Miruna C. Barbu, Eileen Y. Xu, Heather C. Whalley, Riccardo E. Marioni, David J. Porteous, Gail Davies, Ian J. Deary, Gibran Hemani, Klaus Berger, Henning Teismann, Rajesh Rawal, Volker Arolt, Bernhard T. Baune, Udo Dannlowski, Katharina Domschke, Chao Tian, David A. Hinds, Maciej Trzaskowski, Enda M. Byrne, Stephan Ripke, Daniel J. Smith, Patrick F. Sullivan, Naomi R. Wray, Gerome Breen, Cathryn M. Lewis, and Andrew M. McIntosh. Genome-wide meta-analysis of depression identifies 102 independent variants and highlights the importance of the prefrontal brain regions. Nature Neuroscience, 22(3):343–352, March 2019. ISSN 1546-1726. doi: 10.1038/s41593-018-0326-7. URL https://www.nature.com/articles/s41593-018-0326-7. Publisher: Nature Publishing Group.

